# Gene Content Evolution in the Arthropods

**DOI:** 10.1101/382945

**Authors:** Gregg W.C. Thomas, Elias Dohmen, Daniel S.T. Hughes, Shwetha C. Murali, Monica Poelchau, Karl Glastad, Clare A. Anstead, Nadia A. Ayoub, Phillip Batterham, Michelle Bellair, Gretta J. Binford, Hsu Chao, Yolanda H. Chen, Christopher Childers, Huyen Dinh, HarshaVardhan Doddapaneni, Jian J. Duan, Shannon Dugan, Lauren A. Esposito, Markus Friedrich, Jessica Garb, Robin B. Gasser, Michael A.D. Goodisman, Dawn E. Gundersen-Rindal, Yi Han, Alfred M. Handler, Masatsugu Hatakeyama, Lars Hering, Wayne B. Hunter, Panagiotis Ioannidis, Joy C. Jayaseelan, Divya Kalra, Abderrahman Khila, Pasi K. Korhonen, Carol Eunmi Lee, Sandra L. Lee, Yiyuan Li, Amelia R.I. Lindsey, Georg Mayer, Alistair P. McGregor, Duane D. McKenna, Bernhard Misof, Mala Munidasa, Monica Munoz-Torres, Donna M. Muzny, Oliver Niehuis, Nkechinyere Osuji-Lacy, Subba R. Palli, Kristen A. Panfilio, Matthias Pechmann, Trent Perry, Ralph S. Peters, Helen C. Poynton, Nikola-Michael Prpic, Jiaxin Qu, Dorith Rotenberg, Coby Schal, Sean D. Schoville, Erin D. Scully, Evette Skinner, Daniel B. Sloan, Richard Stouthamer, Michael R. Strand, Nikolaus U. Szucsich, Asela Wijeratne, Neil D. Young, Eduardo E. Zattara, Joshua B. Benoit, Evgeny M. Zdobnov, Michael E. Pfrender, Kevin J. Hackett, John H. Werren, Kim C. Worley, Richard A. Gibbs, Ariel D. Chipman, Robert M. Waterhouse, Erich Bornberg-Bauer, Matthew W. Hahn, Stephen Richards

## Abstract

**Background:** Arthropods comprise the largest and most diverse phylum on Earth and play vital roles in nearly every ecosystem. Their diversity stems in part from variations on a conserved body plan, resulting from and recorded in adaptive changes in the genome. Dissection of the genomic record of sequence change enables broad questions regarding genome evolution to be addressed, even across hyper-diverse taxa within arthropods.

**Results:** Using 76 whole genome sequences representing 21 orders spanning more than 500 million years of arthropod evolution, we document changes in gene and protein domain content and provide temporal and phylogenetic context for interpreting these innovations. We identify many novel gene families that arose early in the evolution of arthropods and during the diversification of insects into modern orders. We reveal unexpected variation in patterns of DNA methylation across arthropods and examples of gene family and protein domain evolution coincident with the appearance of notable phenotypic and physiological adaptations such as flight, metamorphosis, sociality and chemoperception.

**Conclusions:** These analyses demonstrate how large-scale comparative genomics can provide broad new insights into the genotype to phenotype map and generate testable hypotheses about the evolution of animal diversity.

## Background

Arthropods (chelicerates, myriapods, crustaceans, and hexapods) constitute the most species-rich and diverse phylum on Earth, having adapted, innovated, and expanded into all major habitats within all major ecosystems. They are found as carnivores, detritivores, herbivores, and parasites. As major components of the world’s biomass, their diversity and ubiquity lead naturally to significant interactions with humanity, as crop pest, disease vectors, food sources, pollinators, and synanthropes. Despite their diversity, arthropods share a deeply conserved and highly modular body plan. They are bilaterally symmetrical, with serially repeated segments along the anterior-posterior axis. Many segments bear paired appendages, which can take the form of antennae, feeding appendages, gills, and jointed legs. Many arthropods have evolved specialized secretions such as venom or silk, extruded from dedicated structures that further capitalize on this segmental modularity. Arthropods also have a hard exoskeleton, composed mostly of chitin, which molts as the animal grows in size. One group of arthropods, the winged insects (Pterygota), took to the skies, bearing up to two pairs of wings as outgrowths of that exoskeleton.

The extraordinary diversity of arthropods is manifested in a series of genomic changes and innovations selected for throughout their evolutionary history. However, linking this phenotypic diversity to underlying genomic changes remains an elusive challenge. The major transitions in arthropod evolution include the differential grouping of body segments into morphological units with a common function (*e.g.*, head, thorax, and abdomen in the Hexapoda) in different taxa, the independent and parallel colonizations of terrestrial and freshwater habitats by ancestrally marine lineages^1,2^, the emergence of active flight in insects^3,4^, and the evolution of insect metamorphosis^5^. Multiple genomic mechanisms might be responsible for such innovations, but the underlying molecular transitions have not been explored on a broad phylogenomic scale. Tracing these transitions at the genomic-level requires mapping whole genome data to a robust phylogenetic framework. Here we explore the evolution of arthropod genomes using a phylogeny-mapped genomic resource of 76 species representing the breath of arthropod diversity.

## Results

### An Arthropod Evolution Resource

As a pilot project for the i5K initiative to sequence 5,000 arthropod genomes^6^, we sequenced and annotated the genomes of 28 arthropod species (Table S1). These include a combination of species of agricultural or ecological importance, emerging laboratory models, and species occupying key positions in the arthropod phylogeny. We combined these newly sequenced genomes with those of 48 previously sequenced arthropods creating a dataset comprising 76 species representing the four extant arthropod subphyla and spanning 21 taxonomic orders. Using the OrthoDB gene orthology database^7^, we annotated 38,195 protein ortholog groups (orthogroups/gene families) among all 76 species (Fig. 1). Based on single-copy orthogroups within and between orders, we then built a phylogeny of all major arthropod lineages (Fig. 2). This phylogeny is mostly consistent with previous arthropod phylogenies^8-10^, with the exception being that we recover a monophyletic Crustacea, rather than the generally accepted paraphyletic nature of Crustacea in respect to Hexapoda, the difference likely due to our restricted taxon sampling (see Methods). We reconstructed the gene content and protein domain arrangements for all 38,195 orthogroups in each of the lineages for the 76 species in the arthropod phylogeny. This resource (available at https://i5k.gitlab.io/ArthroFam/ and Table S11) forms the basis for the analyses detailed below and is an unprecedented tool for identifying and tracking genomic changes over arthropod evolutionary history.

**Figure 1.**
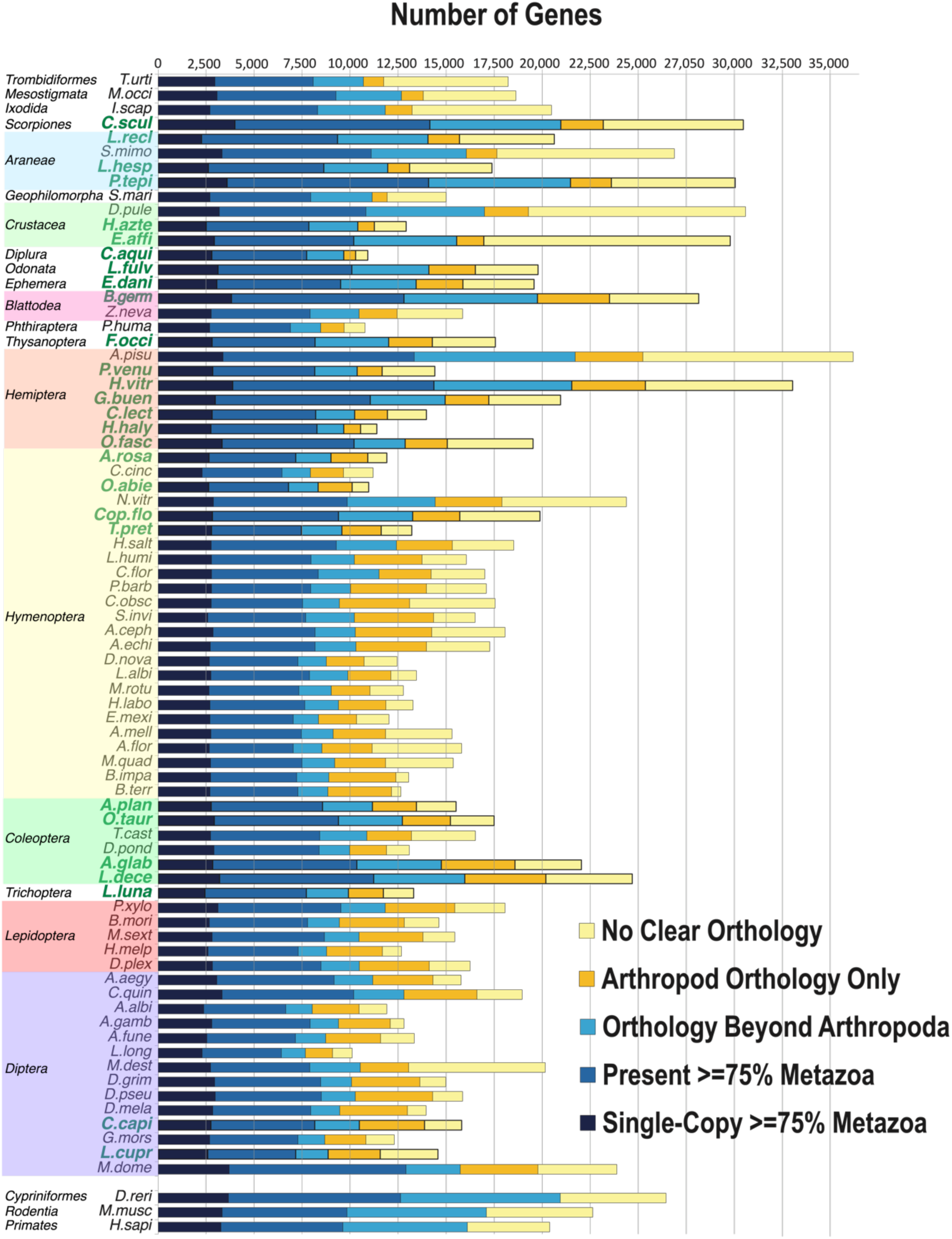
OrthoDB orthology delineation for the i5K pilot species. The bars show Metazoa-level orthologs for the 76 selected arthropods and three outgroup species (of 13 outgroup species used for orthology analysis) partitioned according to their presence and copy-number, sorted from the largest total gene counts to the smallest. The 28 i5K species generated in this study with a total of 533,636 gene models are indicated in bold green font. A total of 38,195 orthologous protein groups were annotated among the total 76 genomes.

**Figure 2:**
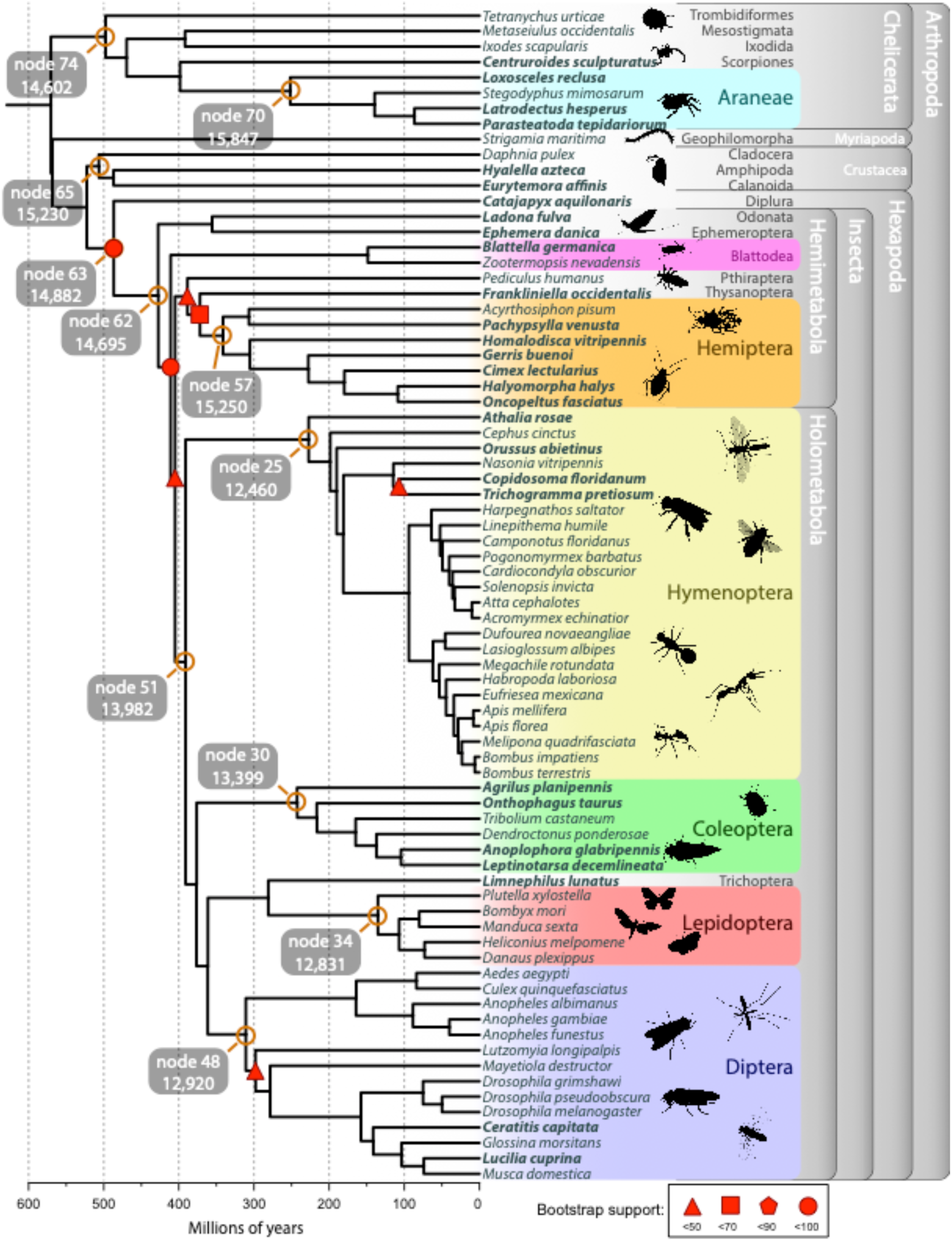
Arthropod phylogeny inferred from 569 to 4,097 single copy protein-coding genes among the six multi-species orders, crustaceans, and non-spider chelicerates (Table S13) and 150 single-copy genes for the orders represented by a single species and the deeper nodes. Divergence times estimated with non-parametric rate smoothing and fossil calibrations at 22 nodes (Table S14). Species in bold are those sequenced within the framework of the i5K pilot project. All nodes, except those indicated with red shapes, have bootstrap support of 100 inferred by ASTRAL. Nodes of particular interest are labeled in orange and referred to in the text. Larger fonts indicate multi-species orders enabling CAFE 3.0 likelihood analyses (see Methods). Nodes leading to major taxonomic groups have been labeled with their node number and the number of genes inferred at that point. See Fig. S17 and Table S12 for full node labels.

### Genomic Change Throughout Arthropod History

Evolutionary innovation can result from diverse genomic changes. New genes can arise either by duplication or, less frequently, by *de novo* gene evolution^11^. Genes can also be lost over time, constituting an underappreciated mechanism of evolution^12,13^. Protein domains are the basis of reusable modules for protein innovation and the rearrangement of domains to form new combinations plays an important role in molecular innovation^14^. Together, gene family expansions and contractions and protein domain rearrangements, may coincide with phenotypic innovations in arthropods. We therefore searched for signatures of such events corresponding with pivotal phenotypic shifts in the arthropod phylogeny.

Using ancestral reconstructions of gene counts (see Methods), we tracked gene family expansions and losses across the arthropod phylogeny. Overall, we inferred 181,157 gene family expansions and 87,505 gene family contractions. 68,430 gene families were inferred to have gone extinct in at least one lineage, and 9,115 families emerged in different groups. We find that, of the 268,662 total gene family changes, 5,843 changes are statistically rapid (see Methods), with the German Cockroach having the most rapid gene family changes (Fig. 3E). The most dynamically-changing gene families encode proteins involved in functions of xenobiotic defense (cytochrome P450s, sulfotransferases), digestion (peptidases), chitin exoskeleton structure and metabolism, multiple zinc finger transcription factor types, HSP20 domain stress response, fatty acid metabolism, chemosensation, and ecdysteroid (molting hormone) metabolism (Table S15). Using the estimates of where in the phylogeny these events occurred, we can infer characteristics of ancestral arthropods. For example, we identified 9,601 genes in the last insect common ancestor (LICA) and estimate ∼14,700 LICA genes after correcting for unobserved gene extinctions (Fig. 2, Fig. S5, and Table S16). We reconstructed similar numbers for ancestors of the six well-represented arthropod taxa in our sample (Fig. 2 and Table S16). Of the 9,601 genes present in LICA, we identified 147 emergent gene families (*i.e.*, lineage-restricted families with no traceable orthologs in other clades) which appeared concurrently with the evolution of insects (Fig. 3A, Fig. 2 node 62, Table S18). GO-term analysis of these 147 gene families recovered multiple key functions, including cuticle and cuticle development (suggesting changes in exoskeleton development), visual learning and behavior, pheromone and odorant binding (suggesting the ability to sense in terrestrial/aerial environments rather than aquatic), ion transport, neuronal activity, larval behavior, imaginal disc development, and wing morphogenesis. These emergent gene families likely allowed insects to undergo substantial diversification by expanding chemical sensing, such as an expansion in odorant binding to locate novel food sources and fine-tune species self-recognition^15-17^. Others, such as cuticle proteins underlying differences in exoskeleton structure, may enable cuticle properties optimized for diverse environmental habitats or life history stages^18^. In contrast, the data reveal only ten gene families that arose along the ancestral lineage of the Holometabola (Fig. 3B, Table S19), implying that genes and processes required for the transition to holometabolous development, such as imaginal disc development, were already present in the hemimetabolous ancestors. This is consistent with Truman and Riddiford’s model that the holometabolous insect larva corresponds to a late embryonic state of hemimetabolous insects^19^.

**Figure 3:**
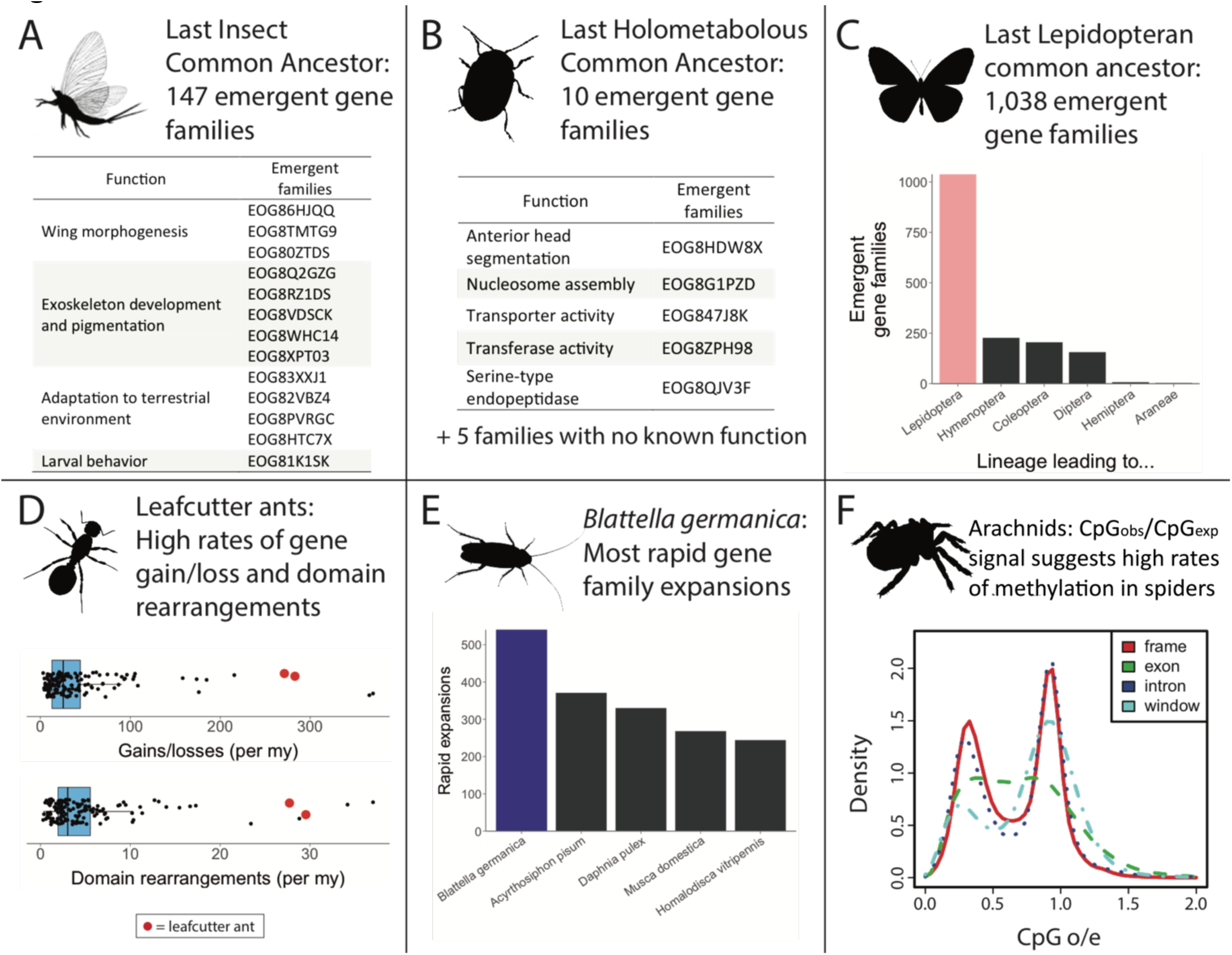
Summary of major results from gene family, protein domain, and methylation analyses. **A.** We identify 147 gene families emerging during the evolution of insects, including several which may play an important role in insect development and adaptation. **B.** Contrastingly, we find only ten emergent gene families during the evolution of holometabolous insects, indicating many gene families were already present during this transition. **C.** Among all lineage nodes, we find that the node leading to Lepidoptera has the most emergent gene families. **D.** We find that rates of gene gain and loss are highly correlated with rates of protein domain rearrangement. Leafcutter ants have experienced high rates of both types of change. **E.** *Blattella germanica* has experienced the highest number of rapid gene family changes, possibly indicating its ability to rapidly adapt to new environments. **F.** We observe signals of CpG methylation in all Araneae (spiders) genomes investigated (species shown: the brown recluse spider, *Loxosceles reclusa*) and the genome of the bark scorpion, *Centruroides exilicauda*. The two peaks show different CG counts in different gene features, with depletion of CG sequences in the left peak due to methylated C’s mutating to T. This suggests epigenetic control of a significant number of spider genes. Additional plots for all species in this study are shown in Fig. S8.

We identified numerous genes that emerged in specific orders of insects. Strikingly, we found 1,038 emergent gene families in the first ancestral Lepidoptera node (Fig. 3C). This node has by far the most emergent gene families, with the next highest being the node leading to the bumble bee genus *Bombus* with 860 emergent gene families (Fig S10). Emergent lepidopteran gene families show enrichment for functional categories such as peptidases and odorant binding. Among the other insect orders, we find 227 emergent families in the node leading to the Hymenoptera, 205 in that leading to Coleoptera, and 156 in that leading to Diptera. Though our sampling is extensive, it is possible that gene families we have classified as emergent may be present in unsampled lineages.

Similarly, we reconstructed the protein domain arrangements for all nodes of the arthropod phylogeny, that is, the permutations in protein domain type per (multi-domain) gene. In total, we can explain the underlying events for more than 40,000 domain arrangement changes within the arthropods. The majority of domain arrangements (48% of all observable events) were formed by a fusion of two ancestral arrangements, while the fission of an existing arrangement into two new arrangements accounts for 14% of all changes. Interestingly, 37% of observed changes can be explained by losses (either as part of an arrangement (14%) or the complete loss of a domain in a proteome (23%)), while emergence of a novel protein domain is a very rare event, comprising only 1% of total events.

We observe high concordance between rates of gene family dynamics and protein domain rearrangement (Figs. 4 & S28). In some cases, we find specific examples of overlap between gene family and protein domain evolution. For example, spiders have the characteristic ability to spin silk and are venomous. Correspondingly, we identify ten gene families associated with venom or silk production that are rapidly expanding within Araneae (spiders, Table S20). In parallel, we find a high rate of new protein domains in the subphylum Chelicerata, including a large number within Araneae associated with venom and silk production. For example, ‘spider silk protein 1’ (Pfam ID: PF16763), ‘Major ampullate spidroin 1 and 2’ (PF11260), ‘Tubuliform egg casing silk strands structural domain’ (PF12042), and ‘Toxin with inhibitor cystine knot ICK or Knottin scaffold’ (PF10530) are all domains that emerged within the spider clade. Venom domains also emerged in other venomous chelicerates, such as the bark scorpion, *Centruroides sculpturatus*.

**Figure 4:**
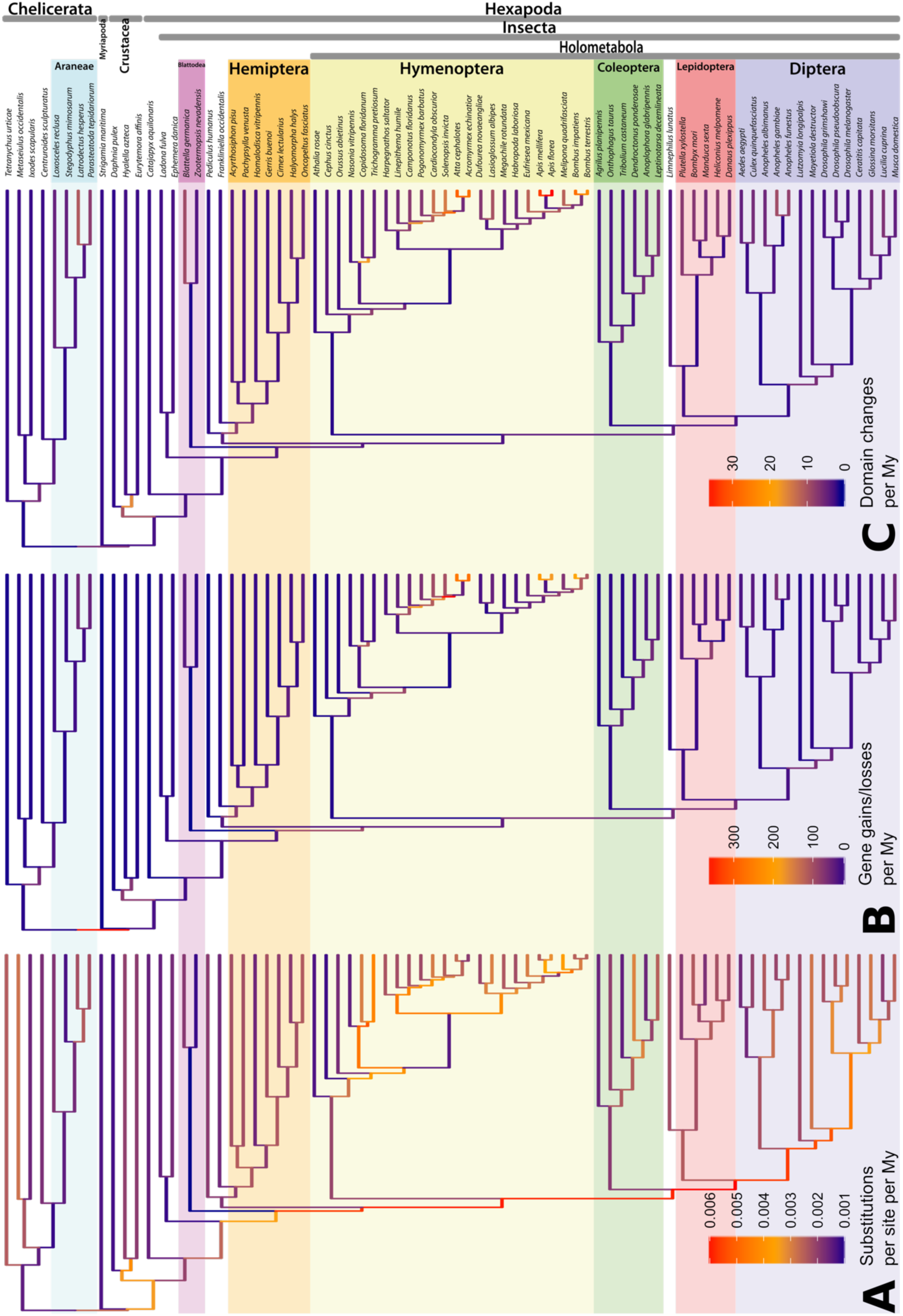
Rate of genomic change along the arthropod phylogeny: **A**, frequency of amino acid substitutions per site, **B**, gene gains/losses and **C**, domain changes. All rates are averaged per My and color-indicated as branches of the phylogenetic tree. Species names are shown on the right; specific subclades are highlighted by colors according to the taxonomic groups noted in Fig. 2.

We identified gene family changes that may underlie unique phenotypic transitions. The evolution of eusociality among three groups in our study, bees and ants (both Hymenoptera), and termites (Blattodea), requires these insects to be able to recognize other individuals of their colony (such as nest mates of the same or different caste), or invading individuals (predators, slave-makers and hosts) for effective coordination. We find 41 functional terms enriched for gene family changes in all three groups, with multiple gene family gains related to olfactory reception and odorant binding (Table S21) in agreement with previous chemoreceptor studies of these species^20,21^.

Finally, we observe species-specific gene family expansions that suggest biological functions under selection. The German cockroach *Blattella germanica*, a pervasive tenant in human dwellings across the world, has experienced the highest number of rapidly evolving gene families among the arthropods studied here, in agreement with a previously reported major expansion of chemosensory genes^22^. We also find the largest number of domain rearrangement events in *B. germanica*. The impressive capability of this cockroach to quickly adapt to changing environments could be linked to these numerous and rapid evolutionary changes at the genomic level and warrants more detailed investigation.

### Evolutionary Rates Within Arthropod History

The rate of genomic change can reflect key events during evolution along a phylogenic lineage. Faster rates might imply small population sizes or strong selective pressure, possibly indicative of rapid adaptive radiations, and slower rates may indicate stasis. Studying rates of change requires a time-calibrated phylogeny. For this, we used 22 fossil calibration points^8,23^ and obtained branch lengths for our phylogeny in millions of years (My) (Fig. 2) that are very similar to those obtained by Misof et al.^8^ and Rota-Stabelli et al.^9^.

We examined the rates of three types of genomic change: (i) amino acid substitutions, (ii) gene duplications and gene losses, and (iii) protein domain rearrangements, emergence, and loss. While clearly not changing in a clock-like manner, all types of genomic change have a strikingly small amount of variation in rate among the investigated species (Fig. 4). We estimate an average amino acid substitution rate of 2.54 x 10^−3^ substitutions per site per My with a standard deviation of 1.11 x 10^−3^. The slowest rate is found in the branch leading to the insect order Blattodea (cockroaches and termites), while the fastest rates are found along the short branches during the early diversification of Holometabola, suggesting a period of rapid evolution, a pattern similar to that found for amino acid sequence evolution during the Cambrian explosion^24^. Other branches with elevated amino acid divergence rates include those leading to Acarina (mites), and to the Diptera (flies).

Though we observe thousands of genomic changes across the arthropod phylogeny, they are mostly evenly distributed (Fig. 3D). Rates of gene duplication and loss show remarkably little variation, both across the tree and within the six multi-species orders (Table S13). Overall, we estimate an average rate of 43.0 gains/losses per My, but with a high standard deviation of 59.0 that is driven by a few lineages with greatly accelerated rates. Specifically, the terminal branches leading to the leafcutter ants *Atta cephalotes* and *Acromyrmex echinatior* along with the internal node leading to the leafcutter ants and the red fire ant (node HY29) have exceptionally high gene gain/loss rates of 266, 277, and 370 per My, respectively (Fig. 3D). This is an order of magnitude higher than average, as previously reported among leafcutter ants^25^. Removing these nodes, the average becomes 27.2 gains/losses per My (SD 19.7). Interestingly, the high gain/loss rates observed in these ants, in contrast to other arthropods, are not due to large gene content change in a small number of gene families. They are instead due mostly to single gene gains or losses in a large number of gene families.

Regarding protein domain rearrangements, which mainly arise from duplication, fusion and terminal losses of domains^26^, we estimate an average rate of 5.27 events per My, approximately eight-fold lower than the rate of gene gain/loss. Interestingly, we discovered a strong correlation between rates of gene gain/loss and domain rearrangement (Figs. 3D, 4, & S28). For example, terminal branches within the Hymenoptera have an accelerated rate of domain rearrangement, which coincides with the increased rate of gene gains and losses observed along those branches. This novel finding is surprising, given that these processes follow largely from different underlying genetic events.

Our examination found no correlation between variation in amino acid substitution rates and rates of gene gain/loss or domain rearrangement rates (Figs. 4 & S28). Branches with accelerated rates of amino acid substitution, such as the lineage leading to the most recent common ancestor of the insect superorder Holometabola, do not show corresponding increases in gene gain/loss rates. Similarly, the hymenopteran lineages displaying the fastest rate of gene gain/loss in our analysis do not display higher rates of amino acid substitutions.

### Control of Novel Genes: Methylation Signals in Arthropod Genomes

Our description of gene family expansions in arthropods by gene duplication naturally suggests the need for differential control of duplicated genes. Insect epigenetic control by CpG methylation is important for caste development in honey bees^27^ and polyphenism in aphids^28^. However, signals of methylation are not seen in every insect, and the entire Dipteran order appears to have lost the capacity for DNA methylation. Given this diversity in the use of, and capacity for epigenetic control by DNA methylation, we searched for signals of CpG methylation in our broader sampling of arthropod genomes. We find several independent losses of the DNA methylation machinery across the arthropods (Fig. S7)^29^. This indicates that DNA methylation is not universally necessary for development and that the DNA methyltransfereases in insects may function in ways not previously appreciated^30^. Additionally, putative levels of DNA methylation vary considerably across arthropod species (Figs. S7, S8). Notably, the hemimetabolous insects and non-insect arthropods show higher levels of DNA methylation signals than the holometabolous insects^29^. Araneae (spiders), in particular, show clear bimodal patterns of methylation (Figs. 3F & S8); with some genes displaying high methylation signals and others not. A possible connection between spider bimodal gene methylation and their proposed ancestral whole genome duplication will require additional investigation. This pattern is also found in some holometabolous insects, suggesting that the division of genes into methylated and unmethylated categories is a relatively ancient trait in Arthropoda, although many species have since lost this clear distinction. Finally, some taxa, particularly in Hymenoptera, show higher levels of CpG di-nucleotides than expected by chance alone, which may be a signal of strong effects of gene conversion in the genome^31^.

### Concluding remarks and future directions

The i5K pilot initiative has assembled an unparalleled genomic dataset for arthropods research and conducted a detailed phylogenetic analysis of evolutionary changes at the genomic level within this diverse and fascinating phylum. The combined research output of species-level i5K work has been substantial and wide-ranging, addressing pests of agricultural crops^32,33^ and animals^34^, urban^20,35^ and forest^36^ pests, biocontrol species^37^, along with developmental models^18,38,39^, indicators of water quality and models for toxicology^15,40^ (Table S1).

Here, in contrast, we take a broad overview generating a comparative genomics resource for a phylum with an evolutionary history of over 500 million years. Our analysis identifies multiple broad patterns such as the very small number of novel protein domains, and a surprising lack of variation in the rates of some types of genomic change. We pinpoint the origin of specific gene families and trace key transitions during which specific gene families or protein domains have undergone rapid radiations or contractions. An overview of the diversity and evolution of TEs found large intra-and inter-lineage variation in both TE content and composition^41^.

Nonetheless, drawing functional biological conclusions from these data is not straightforward. In some cases, the link between specific gene families and their biological function is clear. This is true for genes related to specific physiological function (*e.g.*, olfaction), or to the production of specific compounds (*e.g.*, silk or venom). However, for many gene families, there is no known function, highlighting the need for functional genomic studies. For example, emergent gene families, such as those identified in the Lepidoptera and for rapidly evolving and diverging gene families, cannot be studied in the dipteran *Drosophila* model.

A key consequence of the relatively stable rate of gene family and protein domain change across the arthropod tree is that major morphological transitions (e.g. full metamorphosis, wing emergence, Table S17) could not easily be identified by surges in gene content or protein domain change. There are two possible exceptions in our data. We see an increased rate of gene family extinction along the ancestral nodes from the ancestor of the cockroach and termites and hemimetabolous insects to the ancestor of Lepidoptera and Diptera (Fig S20), suggesting evolution by gene loss^12,42^. This rate increase is not seen in wing evolution. The second possible exception is that of whole genome duplications (as proposed in spiders^39^), when there is a temporary opening of the “evolutionary search space” of gene and protein domain content. This overall finding is in line with the emerging understanding that morphology is effected by complex gene networks, which are active mostly during ontogenetic processes^43^, rather than by individual “morphology genes”. Morphological innovations are often based on modulating the timing and location of expression, rewiring of existing gene networks, and assembling new networks using existing developmental toolkit genes^44^. The current study was unable to address the evolutionary non-coding sequences such as enhancers, promoters and small and other non-coding RNAs underlying these networks due to the lack of sequence conservation over large evolutionary distances, however our results underscore their evolutionary importance.

The advent of affordable and widely transferable genomics opens up many avenues for evolutionary analyses. The genome is both the substrate and record of evolutionary change, and it encodes these changes, but the connection is far from simple. A better understanding of the genotype-phenotype map requires in-depth experimental studies to test hypotheses generated by genomic analyses, such as those presented here. The diversity of arthropods provides unparalleled taxonomic resolution for phenotypic change, which, combined with the experimental tractability of many arthropods, suggests a productive area of future research using and building upon the resource established herein.

## Methods

### Sequencing, assembly and annotation

Twenty-eight arthropod species were sequenced using Illumina short read technology. In total, 126 short read libraries were generated and sequenced to generate 4.9 Tb of raw nucleotide sequence (Table S2). For individual species, reads were assembled using AllpathsLG^45,46^ followed by refinements employing Atlas-Link^47^ and Gapfill^48^. Version 1.0 assemblies had minimum, mean, and maximum scaffold N50 lengths of 13.8 kb, 1.0 Mb and 7.1 Mb (Table S3). Following re-assembly and collapsing of unassembled haplotypes using Redundans^49^, version 2.0. assemblies had minimum, mean and maximum contig N50 lengths of 11.1 kb, 166.2 kb and 857.0 kb with a mean scaffold N50 lengths of 619 kb (Table S3). The redundans software and new assemblies became available late in the project timeline, and thus automated gene annotations, orthologous gene family identification in OrthoDB and analysis were performed on the Version 1 ALLPATHS-LG based assemblies.

To support the annotation, RNAseq data were generated from 25 species for which no data were available (Table S4). A MAKER^50^ based automated annotation pipeline was applied to the 1.0 assembly of each species with species-specific input RNAseq data and alignment data from a non-redundant metazoan protein sequence set containing all available arthropod protein sequences (see Supplementary methods). This pipeline was applied to 28 species with annotatable genome assemblies generating 533,636 gene models, with minimum, mean, and maximum gene model numbers of 10,901, 19,058, and 33,019 per species (Table S5, see Table S7 for completeness statistics). Many of these gene models were manually curated using the i5k Workspace@NAL^51^. Given the magnitude of this manual task, the greatest fraction of gene models manually confirmed for a species was 15 %. The analyses presented here were performed on the automatically generated gene models.

### Orthology prediction

Orthology delineation is a cornerstone of comparative genomics, offering qualified hypotheses on gene function by identifying “equivalent” genes in different species. We used the OrthoDB^7^ (www.orthodb.org) orthology delineation process that is based on the clustering of best reciprocal hits (BRHs) of genes between all pairs of species. Clustering proceeds first by triangulating all BRHs and then subsequently adding in-paralogous groups and singletons to build clusters of orthologous genes. Each of these ortholog groups represent all descendants of a single gene present in the genome of the last common ancestor of all the species considered for clustering^52^.

The orthology datasets computed for the analyses of the 28 i5K pilot species, together with existing sequenced and annotated arthropod genomes were compiled from OrthoDB v8^53^, which comprises 87 arthropods and an additional 86 other metazoans (including 61 vertebrates). Although the majority of these gene sets were built using MAKER (Table S6), variation in annotation pipelines and supporting data, introduce a potential source of technical gene content error in our analysis.

Orthology clustering at OrthoDB included ten of the i5K pilot species (*Anoplophora glabripennis, Athalia rosae, Ceratitis capitata, Cimex lectularius, Ephemera danica, Frankliniella occidentalis, Ladona fulva, Leptinotarsa decemlineata, Orussus abietinus, Trichogramma pretiosum*). The remaining 18 i5K pilot species were subsequently mapped to OrthoDB v8 ortholog groups at several major nodes of the metazoan phylogeny. Orthology mapping proceeds by the same steps as for BRH clustering, but existing ortholog groups are only permitted to accept new members, *i.e.*, the genes from species being mapped are allowed to join existing groups if the BRH criteria are met. The resulting ortholog groups of clustered and mapped genes were filtered to select all groups with orthologs from at least two species from the full set of 76 arthropods, as well as retaining all orthologs from any of 13 selected outgroup species for a total of 47,281 metazoan groups with orthologs from 89 species. Mapping was also performed for the relevant species at the following nodes of the phylogeny: Arthropoda (38,195 groups, 76 species); Insecta (37,079 groups, 63 species); Endopterygota (34,614 groups, 48 species); Arachnida (8,806 groups, 8 species); Hemiptera (8,692 groups, 7 species); Hymenoptera (21,148 groups, 24 species); Coleoptera (12,365 groups, 6 species); and Diptera (17,701, 14 species). All identified BRHs, amino acid sequence alignment results, and orthologous group classifications were made available for downstream analyses: http://ezmeta.unige.ch/i5k.

### Arthropod phylogeny

We reconstructed the arthropod phylogeny (Fig. 2) using protein sequences from the 76 genomes. Six different phylogenetic reconstruction approaches generated a consistent relationship among the orders (see Supplemental Methods), corresponding with previously inferred arthropod phylogenies^8-10^.

Of the six orders in our dataset represented by multiple species (Figs. S11-16), relationships within the Araneae, Hemiptera, Coleoptera, and Lepidoptera were identical, regardless of the tree building method used. Within the Hymenoptera, the only disagreement between methods concerned the position of the parasitoid wasps within the Chalcidoidea, with three methods placing *Copidosoma floridanum* as sister to *Nasonia vitripennis* (in agreement with recent phylogenomic research^54^), and the three other methods placing *C. floridanum* as sister to *Trichogramma pretiosum* (Fig. S13). Within the Diptera, we obtained a sister group relationship between the sand fly, *Lutzomyia longipalpis*, and the Culicidae, but this was not a stable topology across methods (Fig. S16).

The most contentious nodes in the phylogeny involve the relationship of crustaceans and hexapods. We recover a monophyletic Crustacea that represent the sister clade to Hexapoda (Fig. 2), in contrast to recent analyses suggesting this group is paraphyletic in respect to Hexapoda^55^. However, an extensive phylogenetic investigation (Supplementary Results, Fig. S9) shows that regardless of the inference method used, the relationships among the crustacean and hexapod lineages remain uncertain. Aside from these few discrepancies, branch support values across the tree were high for all tree building methods used. Even when bootstrap support was < 100 %, all methods still inferred the same topology among the species included. The most likely reason for the difference from the current consensus is poor taxon sampling. Importantly, remipedes (the possible sister group of the hexapods) are missing from our taxon sampling, as are mystacocarids, ostracods and pentatomids, and may change this result to the current consensus when added as was seen in^55^.

### Divergence time estimation

Phylogenetic branch lengths calibrated in terms of absolute time are required to study rates of evolution and to reconstruct ancestral gene counts. We used a non-parametric method of tree smoothing implemented in the software r8s^56^ to estimate these divergence times. Fossil calibrations are required to scale the smoothed tree by absolute time. We relied on Wolfe et al.’s^23^ aggregation of deep arthropod fossils with additional recent fossils used by Misof et al.^8^ (Table S14). The results indicate that the first split within arthropods (the chelicerate-mandibulate split) occurred ∼570 million years ago (mya). We estimate that within the chelicerates, arachnids radiated from a common ancestor ∼500 mya. Within the mandibulates, myriapods split from other mandibulates ∼570 mya. Crustaceans started radiating ∼506 mya, and insects started radiating ∼430 mya.

### Substitution rate estimation

To estimate substitution rates per year on each lineage of the arthropod phylogeny, we divided the expected number of substitutions (the branch lengths in the unsmoothed tree) by the estimated divergence times (the branch lengths in the smoothed tree) (Fig. 4).

### Gene family analysis

With the 38,195 orthogroups and the ultrametric phylogeny we were able to perform the largest gene family analysis of any group of taxa to date. In this analysis we were able to estimate gene turnover rates (*λ*) for the six multi-species taxonomic orders, to infer ancestral gene counts for each taxonomic family on each node of the tree, and to estimate gene gain/loss rates for each lineage of the arthropod phylogeny. The size of the dataset and the depth of the tree required several methods to be utilized.

Gene turnover rates (λ) for the six multi-species orders were estimated with CAFE 3.0, a likelihood method for gene-family analysis^57^. CAFE 3.0 is able to estimate the amount of assembly and annotation error (ε) present in the input gene count data. This is done by treating the observed gene family counts as distributions rather than certain observations. CAFE can then be run repeatedly on the input data while varying these error distributions to calculate a pseudo-likelihood score for each one. The error model that is obtained as the minimum score after such a search is then used by CAFE to obtain a more accurate estimate of λ and reconstruct ancestral gene counts throughout the tree (Table S12). However, with such deep divergence times of some orders, estimates of ε may not be accurate. CAFE has a built-in method to assess significance of changes along a lineage given an estimated λ and this was used to identify rapidly evolving families within each order. We partitioned the full dataset of 38,195 orthogroups for each order such that taxa not in the order were excluded for each family and only families that had genes in a given order were included in the analysis. This led to the counts of gene families seen in Table S11.

For nodes with deeper divergence times across Arthropoda, likelihood methods to reconstruct ancestral gene counts such as CAFE, become inaccurate. Instead, a parsimony method was used to infer these gene counts across all 38,195 orthogroups^58^. Parsimony methods for gene family analysis do not include ways to assess significant changes in gene family size along a lineage. Hence, we performed a simple statistical test procedure for each branch to assess whether a given gene family was changing significantly: under a stochastic birth-death process of gene family evolution, and within a given family, the expected relationship between any node and its direct ancestor is that no change will have occurred. Therefore, we took all differences between nodes and their direct descendants in a family and compared them to a one-to-one linear regression. If any of the points differ from this one-to-one line by more than two standard deviations of the variance within the family, it was considered a significant change and that family is rapidly evolving along that lineage. Rates of gene gain and loss were estimated in a similar fashion to substitution rates. We counted the number of gene families inferred to be changing along each lineage and divided that by the estimated divergence time of that lineage (Fig. 4). To quantify the effect of any single species on the parsimony gene family reconstructions, we performed 100 jackknife replicates while randomly removing 5 species from each replicate. We find that ancestral gene counts are not greatly impacted by the presence or absence of any single genome (Supplemental Fig S32).

To estimate ancestral gene content (*i.e.*, the number of genes at any given node in the tree), we had to correct for gene losses that are impossible to infer given the present data. To do this, we first regressed the number of genes at each internal node with the split time of that node and noticed the expected negative correlation of gene count and time (Fig. S5) (*r*^*2*^=0.37; *P*=4.1 x 10^−9^). We then took the predicted value at time 0 (present day) as the number of expected genes if no unobserved gene loss occurs along any lineage and shifted the gene count of each node so that the residuals from the regression matched the residuals of the 0 value.

### Protein domain evolution analysis

We annotated the proteomes of all 76 arthropod species and 13 outgroup species with protein domains from the Pfam database (v30)^59^. Thereby, every protein was represented as a domain arrangement, defined by its order of domains in the amino acid sequence. To prevent evaluating different isoforms of proteins as additional rearrangement events, we removed all but the longest isoform. Repeats of a same domain were collapsed to one instance of the domain (A-B-B-B-C → A-B-C), since copy numbers of some repeated domains can vary strongly even between closely related species^60,61^. To be able to infer all rearrangement events over evolutionary time, we reconstructed the ancestral domain content of all inner nodes in the phylogenetic tree via the DomRates tool (http://domainworld.uni-muenster.de/programs/domrates/) based on a combined parsimony approach (see Supplementary Methods). Six different event types were considered in this study (Fig. S6): fusion, fission, terminal loss/emergence and single domain loss/emergence. For the rate calculation just all arrangement changes were considered that could be explained by exactly one of these event types, while all arrangements were ignored that could not be explained by one of these events in a single step or if multiple events could explain a new arrangement.

## Supporting information

Large supplemental tables

## Acknowledgments

We thank the staff at the Baylor College of Medicine Human Genome Sequencing Center for their contributions. Genome sequencing, assembly and annotation was funded by National Human Genome Research Institute grant U54 HG003273 to R.A.G. GWCT and MWH are funded by NSF DBI-1564611. ED was funded by the Deutsche Forschungsgemeinschaft (DFG, German Research Foundation) – 281125614 / GRK2220. RMW, PI and EMZ were funded by The Swiss National Science Foundation (PP00P3_170664 to RMW, 31003A_143936 to EMZ). Contributions by DDM and AW were supported in part by NSF-DEB grant 1355169 and USDA-APHIS Cooperative Agreement 15-8130-0547-CA to DDM. BM and ON acknowledge the German Research foundation (NI 1387/3-1, MI 649/12-1) and the Leibnitz Graduate School on Genomic Biodiversity Research. CS was supported by the Blanton J. Whitmire endowment, Housing and Urban Development NCHHU-0007-13, National Science Foundation 1557864 and Alfred P. Sloan Foundation 2013-5-35 MBE. Funding from Australian Wool Innovation (to P.B. and R.B.G.) and the Australian Research Council (to R.B.G.) is gratefully acknowledged. Support to R.B.G.’s laboratory by YourGene Bioscience and Melbourne Water Corporation is gratefully acknowledged. This project was also supported by a Victorian Life Sciences Computation Initiative (VLSCI; grant number VR0007) on its Peak Computing Facility at the University of Melbourne, an initiative of the Victorian Government (R.B.G.). C.A.A. holds an NSERC Postdoctoral Fellowship. N.D.Y. holds an NHMRC Early Career Research Fellowship. P.K.K. is the recipient of a scholarship (STRAPA) from the University of Melbourne.

## Author Contributions

GWCT performed phylogenetic analyses/reconstructions and designed the website. GWCT, ED and RMW performed gene content and protein domain analysis and interpretation and contributed data to the website. KG and MADG performed methylation analysis. MB, HC, HD, HVD, SD, YH, JCJ, SLL, MM, NO-L, DMM, RAG and SR managed and performed sequence library and sequence generation. SCM, JQ, DSTH, KCW and SR performed genome assemblies, DSTH generated automated annotations, and DK, ES and SR submitted data to public databases. MP, CC, MM-T performed and supported manual annotation. NAA, JBB, DB, HC, JJD, LE, CEL, JG, RBG, CAA, PKK, NDY, PB, TP, DEG-R, AMH, MH, LH, WBH, AK, ARIL, GM, APM, DDM, BM, ON, SRP, KAP, MP, RSP, HCP, N-MP, DR, CS, SDS, EDS, DBS, RS, MRS, NUS, and EEZ provided species materials and expertise. RMW, PI and EMZ performed orthology analysis. YL and MEP performed GO analysis, AW, DDM and MF assessed coleopteran and dipteran gene content change. JG summarized chelicerate gene families. MEP, KJH, JHW, KCW, GWCT, ED, RMW, ADC, EBB, MWH and SR prepared the manuscript contributed to project management and provided leadership.

