## Supplemental text for "Gene Content Evolution in the Arthropods"

#### Supplementary Materials

#### Table of Contents

**1. SUPPLEMENTAL METHODS** 3

*1.01 Species Selection and DNA Isolation* 3
*1.02 Genome sequencing and assembly strategy* 4
*1.03 DNA sequencing library preparation* 4
*1.04 DNA sequencing* 5
*1.05 Genome assembly* 5
*1.06 Platanus assembly of a dipluran* 6
*1.07 Redundans genome assembly improvement* 6
*1.08 Simple GC content analysis* 7
*1.09 Sequence read K-mer distributions* 7
*1.10 RNA sequencing* 8
*1.11 Automated gene model annotation* 8
*1.12 Community gene curation and annotation* 9
*1.13 Gene model quality assessment* 10
*1.14 Orthology prediction* 10
*1.15 Phylogeny inference* 12
*1.16 Phylogeny inference – an additional point brought up during review.* 14
*1.17 Divergence time estimation* 15
*1.18 Substitution rate estimation* 15
*1.19 Gene family analysis* 15
*1.20 Jackknifing tests on parsimony reconstructions of gene families* 16
*1.21 GO enrichment tests* 17
*1.22 Protein Domain evolution analysis* 17

**2. SUPPLEMENTAL RESULTS** 20

*2.01 DNA methylation across the arthropods* 20
*2.02 Pancrustacea phylogeny* 22
*2.03 Gene families evolving on the most lineages* 23
*2.04 Coleopteran gene family evolution summary* 23
*2.05 Diptera gene family evolution summary* 24
*2.06 Protein domain analysis* 26
*2.07 Protein Innovation: Silk and venom domain emergences in Chelicerates* 27

**3. SUPPLEMENTARY TABLES.** 28

*3.01. List of Large Supplementary Tables as worksheets in Microsoft Excel file "Large Supplementary Tables".* 28

*3.02. Calculated rates of rearrangement events.* 30
*3.03. Calculated exact numbers of rearrangement events.* 30

**4. SUPPLEMENTARY FIGURES. 31**

**Figure S1. Counts of 195 i5K Nominated Species by Order.** 31
**Figure S2. Assembly and Maker 2.0 CDS GC content for Redundans assemblies.** 32
**Figure S3. Kmer analysis of i5K pilot species 500bp read libraries at 17, 21 and 31 bp.** 33
**Figure S4. OrthoDB orthology delineation for the i5K pilot species.** 38
**Figure S5: Estimating gene counts at ancestral nodes.** 39
**Figure S6. Protein domain reconstruction and rearrangement event inference.** 40
**Figure S7. Presence of DNA methylation machinery across the arthropods.** 41
**Figure S8. Patterns of DNA methylation CpG 0/e signals of 72 arthropod species.** 42
**Figure S9. Support for 15 alternate crustacean topologies with 3 gene sets.** 62
**Figure S10: Novel gene family expansions and extinctions.** 63
**Figure S11: Araneae tree.** 64
**Figure S12: Hemiptera tree.** 65
**Figure S13: Hymenoptera tree.** 66
**Figure S14: Coleoptera tree.** 67
**Figure S15: Lepidoptera tree.** 68
**Figure S16: Diptera tree.** 69
**Figure S17: Fig 2. with all nodes labeled.** 70
**Figure S18: Gene family emergences vs. gene family extinctions.** 71
**Figure S19: Rapid gene family expansions vs. rapid gene family contractions.** 72
**Figure S20. Rates of gene family emergences and extinctions.** 73
**Figure S21. Distribution of domain rearrangement events.** 74
**Figure S22. Distribution of fusion events** 75
**Figure S23. Distribution of fission events.** 76
**Figure S25. Distribution of terminal emergence events** 78
**Figure S29. Significant GO terms in gained domain arrangements.** 82
**Figure S30. Dipteran gene content descriptive statistics.** 83
**Figure SG2_31: Comparison of branch lengths from 150 backbone or 1,000's of genes.** 84
**Figure SG1_32: Jackknife test of gene gains/losses after randomly removing species.** 85
**Figure SG3_33: Jackknife resampling test of phylogeny branch divergence times.** 86
**Figure SE1_34: Jackknife resampling test of total protein domain change events** 87

**5. SUPPLEMENTAL REFERENCES 88**

#### 1. Supplemental Methods

##### *1.01 Species Selection and DNA Isolation*

As the genome sequencing aspect of this project was a pilot for the i5K project[1], a community genomic infrastructure initiative for arthropods, we took a community approach to species selection. A species nomination page on the i5K wiki website (now at http://i5K.github.io/legacy_i5K_nominations), combined with significant community outreach via multiple large email lists solicited community nominations for 193 species for genomic sequencing at the time of selection. The nomination list continued to grow to 783 species. The nominated species were highly focused on the four holometabolous orders and the Hemiptera (Fig. S1). Narrowing of this nomination list to the sequenced species was based on several factors: 1. Genome size (and thus cost) - initial budgeting for the pilot was based on 500Mb genome sizes as seen previously in Holometabola, but genome sizes are larger outside these orders. Mantids for example have genome sizes around 5Gb, many Crustacea around 3Gb, and spiders 1.2-1.5Gb (all sizes from the animal genome size database[2]). Many species were removed based on this size/cost criterion alone. 2. An active research community increasing the probability of analysis completion and publication and maximizing the number of researchers impacted. 3. The first sequenced representative of an order, both to sample widely in the arthropods, and to increase the probability of changes in gene content being representative of different life history. 4. Scientific significance - for example scientific model species such as the house spider or the milkweed bug, urban pest such as the bed bug and German cockroach, agricultural pest such as the Colorado potato beetle, etc. 5. Some sampling of non-insect arthropods. The Arachnid community in particular narrowed down the list to the four chelicerates chosen. 6. Availability of high quality DNA (50µg was a requested ideal given size cuts for larger insert mate pair libraries, and backup material) and ability to generate inbred lines for better sequence read assembly (although this requirement was often impossible to fulfill). 7. We additionally sought out "basal" insect orders in collaboration with Bernhard Misof and Oliver Niehuis and the 1Kite project[3] to better understand insect evolution. 8. The addition of the velvet worm *E. rowelli* as an outgroup to the arthropods, although the large genome size prevented high quality draft assembly.

DNA was isolated by collaborators using a variety of methods, the most common of which was the Blood & Cell Culture DNA Midi Kit (G/100) (Qiagen Inc., Valencia, California, USA). Genomic DNA was most often isolated from individual adults of both sexes, with additional RNA isolated most often using the TRIzol Reagent (Invitrogen/Thermo Fisher Scientific, Waltham). There was variation in DNA isolation protocols reflecting the variety of difficulties in dealing with the different species. RNA and genomic DNA was shipped to the Baylor College of Medicine Human Genome Sequencing Center on dry ice for library construction, sequencing, assembly and annotation.

##### *1.02 Genome sequencing and assembly strategy*

It is critical that sequence generation be designed with the assembly strategy in mind. We used an Illumina-ALLPATHS-LG[4,5] sequencing and assembly strategy enhanced with Atlas-link and Atlas-gapfill (https://www.hgsc.bcm.edu/software/). This enabled multiple species to be approached in parallel at reduced costs. For most species, we sequenced four libraries of nominal insert sizes 180bp, 500bp, 3kb and 8kb at 40X, 40X, 40X and 20X estimated genome coverage respectively. The amount of sequence generated from each of these libraries is noted in Table S2, with NCBI SRA accessions. In some cases additional libraries with nominal insert sizes of 1kb or 2kb were prepared using the same methods as for the 3kb insert libraries and sequenced for an improved assembly, however the additional sequencing was not found to significantly improve the genome assembly for the additional effort, and the 4 insert library strategy was the primary sequencing dataset for assembly. In one case (the Dipluran *Catajapyx aquilonaris*) the small amount of input DNA precluded the use of the 4 insert DNA library / ALLPATHS-LG strategy so a PLATANUS[6] assembly strategy based on sequencing two libraries of nominal insert size 400bp and 800bp generated from ~25ng DNA isolated from a single individual. Where possible efforts were made to generate at least some sequence from either sex, for example the 180bp, 500bp, and 3kb inserts might come from one sex, and the 8kb insert from the other sex. In three cases, (Hhal, Mhra and Lcup), an additional library was sequenced to generate sequence from the second sex. Finally, whilst the ALLPATHS-LG with the Atlas enhancements can be very successful in our hands[7], the tools struggle on polymorphic input sequence data and approximately half of the genome assemblies had contig N50s < 10kb. Towards the end of this project new assembly tools designed to improve genome assemblies on polymorphic input sequence data became available, and one (REDUNDANS[8]) was successful enough to merit assembly improvement to be attempted on all applicable species. The REDUNDANS software and new assemblies became available late in the project timeline, and thus automated gene annotations, orthologous gene family identification in OrthoDB and analysis were all performed on the Version 1 ALLPATHS-LG based assemblies.

##### *1.03 DNA sequencing library preparation*

To prepare the 180bp and 500bp libraries, we used a gel-cut paired end library protocol. Briefly, 1 µg of the DNA was sheared using a Covaris S-2 system (Covaris, Inc. Woburn, MA) using the 180-bp or 500-bp program. Sheared DNA fragments were purified with Agencourt AMPure XP beads, end-repaired, dA-tailed, and ligated to Illumina universal adapters. After adapter ligation, DNA fragments were further size selected by agarose gel and PCR amplified for 6 to 8 cycles using Illumina P1 and Index primer pair and Phusion® High-Fidelity PCR Master Mix (New England Biolabs). The final library was purified using Agencourt AMPure XP beads and quality assessed by Agilent Bioanalyzer 2100 (DNA 7500 kit) determining library quantity and fragment size distribution before sequencing.

Long mate pair libraries with 3kb or 8kb insert sizes were constructed according to the manufacturer's protocol (Mate Pair Library v2 Sample Preparation Guide art # 15001464 Rev. A PILOT RELEASE). Briefly, 5 µg (for 2 and 3-kb gap size library) or 10 µg (8-10 kb gap size library) of genomic DNA was sheared to desired size fragments by Hydroshear (Digilab, Marlborough, MA), then end repaired and biotinylated. Fragment sizes between 3-3.7 kb (3kb) or 8-10 kb (8kb) were purified from 1% low melting agarose gel and then circularized by blunt-end ligation. These size selected circular DNA fragments were then sheared to 400-bp (Covaris S-2), purified using Dynabeads M-280 Streptavidin Magnetic Beads, end-repaired, dA-tailed, and ligated to Illumina PE sequencing adapters. DNA fragments with adapter molecules on both ends were amplified for 12 to 15 cycles with Illumina P1 and Index primers. Amplified DNA fragments were purified with Agencourt AMPure XP beads. Quantification and size distribution of the final library was determined before sequencing as described above.

##### *1.04 DNA sequencing*

Sequencing was performed on Illumina HiSeq2000s generating 100bp paired end reads according to the manufacturer's specifications. Briefly, sequencing libraries were quantified with an Agilent 2100 Bioanalyzer. Cluster generations were performed on an Illumina cluster station. A total of 101 cycles of sequencing were carried out with varying numbers of barcoded libraries per flow cell lane depending on the amount of sequence required for that library species combination. Sequencing analysis was first done with Illumina analysis pipeline. Sequencing image files were processed to generate base calls and phred-like base quality scores and to remove low-quality reads. All sequence reads generated for this project have been deposited in the sequence read archive (SRA) at NCBI under umbrella bioproject number PRJNA163973. All of the individual bioprojects and SRA accessions for the sequencing are described in Tables S1 and S2.

##### *1.05 Genome assembly*

Sequence reads were prepared using SeqPrep (https://github.com/jstjohn/SeqPrep/) to remove stray Illumina adapters, and mate pair libraries trimmed to 60bp using a custom perl script. Sequence reads were assembled using ALLPATHS-LG (v35218)[4,5] on a large memory computer with 1Tbyte of RAM. We have previously found that ALLPATHS-LG assembly stats can be incrementally improved (20-30% increase in contig and scaffold N50s) with additional efforts to gap fill and scaffold[7]. We used in-house tools Atlas-Link (v.1.0) and Atlas gap-fill (v.2.2) (https://www.hgsc.bcm.edu/software/) to incrementally improve these assemblies. In one case the size of the genome assembly prevented ALLPATHS run completion. Running the estimated 4.5Gb velvet worm assembly required small modifications to the ALLPATHS-LG code enabling input of more than 4 billion input reads. It then required more than 1Tbyte of RAM which was enabled by allowing swap to happen on an SSD drive. Despite these efforts the resulting assembly was not satisfactory. The resulting assembly statistics and NCBI Accessions are shown in Table S3.

##### *1.06 Platanus assembly of a dipluran*

The very small amount of DNA able to be isolated from *Caqu* individuals prevented the use of our standard strategy. Instead we prepared two paired end Illumina libraries with 450 bp and 1,000 bp insert sizes, and generated 300 bp reads using an Illumina MiSeq at ~40X coverage for each library. These reads were then assembled using Platanus version 1.2.4., a program well suited to long reads due to varying k-mer sizes, and efforts made to deal with polymorphism[6]. Platanus assembly produced reasonable contig sizes (N50: 12.8 Kb), but relatively small scaffolds (N50: 32.4 Kb) due to the lack of long range sequence data (Table S3).

##### *1.07 Redundans genome assembly improvement*

Many of the species assemblies had less than desired contiguity due primarily to sequence polymorphism in the input sequence datasets. These datasets were often generated from wildtype individuals as opposed to strains bred for genome homozygosity, and/or from multiple individuals in species with physically small members rather than a single individual due to the DNA requirements. During the course of the project we tried to improve our assemblies using two assembly software packages that attempt to assemble such polymorphic sequence datasets: Platanus[6] and Redundans[8]. In our experiments Platanus often gave worse, equivalent or slightly better assemblies than our initial ALLPATHS-LG/Atlas results with smaller computational demands. However, it most improved the better assemblies, and had less effect on the worst assemblies which were of most concern (results not shown). Overall Platanus improvements were too incremental to justify the community work required of a new reference version. Given its ease of use, we do recommend trying Platanus with polymorphic data, especially with longer Illumina reads as used for our very low input species (see above).

The Redundans assembly improvement tool more fully collapsed un-assembled haplotypes, generating greater improvement in the biological utility of the assemblies. We ran Redundans version 0.12c on all of the species assemblies. As Redundans is an assembly improvement tool it required a genome assembly to reduce by overlapping previously assembled sequences which is detailed for each species in Table S3. Later versions of Redundans do not have this limitation. For each assembly it was fed the 1.0 assembly (allpaths/atlas) contigs, the 1.0 assembly scaffolds, and/or Platanus re-assembly contigs or Platanus re-assembly scaffolds, with the most successful assembly noted in Table S3. We were unable to get the Redundans assembly to complete for time and memory consumption reasons in three cases, Lrec, Mhra and Erow, likely due to the size of these genomes. The REDUNDANS software and new assemblies became available late in the project timeline, and thus automated gene annotations, orthologous gene family identification in OrthoDB and analysis were all performed on the Version 1 ALLPATHS-LG based assemblies.

##### *1.08 Simple GC content analysis*

Fig S2, and Table S9 show GC content for genome assemblies and Maker gene coding sequences. The minimum assembly GC content was 0.272 for the black widow spider, and the maximum 0.512 for the western flower thrips. It is possible the GC content for the spider is low due to assembly issues and it could increase with a higher quality assembly, as happened for the common house spider assembly moving from the 1.0 ALLPATHS to the 2.0 Redundans assembly - although perhaps low GC repeats were collapsed with the Redundans method. Note that in general GC content was very similar between the two assemblies of the same species reflecting the identical input data. For the CDS GC content, the extremes at the low end were not as pronounced with a number of species having 0.37 GC content, but three species had CDS GC content > 0.5 (Hazt:0.5384, Caqu:0.5198, Focc:0.5109).

##### *1.09 Sequence read K-mer distributions*

We assessed sequence read k-mer distributions for k-mer lengths 17, 21 and 31 bp for all of the i5K pilot species using Jellyfish[9] on 500bp sequence data and custom python scripts to generate the plots, (Fig. S3). Genome size was estimated from k-mers using the relationship genome size = (total k-mers - error k-mers)/k-mer peak (A nice methodological description is available here: http://koke.asrc.kanazawa-u.ac.jp/HOWTO/kmer-genomesize.html).

The species can be divided into groups depending on whether there is a classic single diploid k-mer abundance peak as would be expected for highly inbred species or otherwise low polymophism or isogenic species, (Aros, Caqu, Ccap, Dpse - *Drosophila pseudoobscura*, a control species with 20 generations of sib-sib inbreeding, Eaff, Erow, Focc, Hazt, Hhal, Hpun - another control species, Lcup, Oabi, Ofas, Otau, Tpre), two peaks, with polymorphism partially separating haplotypes in the diploid peak (Agla, Bger, Clec, Edan, Hvit, Lhes, Llun, Mhra, Ptep, Pven) (sometimes the two peaks can be barely seen as most k-mers are very low coverage due to more extensive polymorphism) and finally no peaks, where more extensive polymorphism makes large numbers of identical k-mers less likely (Apla, Cexi, Cflo, Gbue, Ldec, Lful, Lrec). Sometimes, but not always, these categories of k-mer plot roughly correspond to the quality of the genome assembly. For example, species with a single diploid peak at high coverage suggesting low polymorphism include the parasitic wasps Aros, Oabi, Tpre, the inbred Medfly and sheep blowfly Ccap and Lcup, all of which assembled relatively well. Interestingly the parasitic wasp Cflo had no k-mer peak, but perhaps the edge of one appearing can be seen in the 17mer plot, this was the worst of the wasp genome 1.0 assemblies with a contig N50 of only 14kb - unusual given the occurrence of haploid males in the Hymenoptera. Reassembly of Cflo using Redundans improved the contig N50 to 42kb. At the other end of the spectrum, species with no peaks in the k-mer plots generally assembled poorly, with some exceptions - for example the dragonfly Lful has reasonable assembly contig N50s of 16 and 70kb for the version 1.0 and

2.0 assemblies respectively. Accurate genome size estimation using k-mer analysis is also dependent on peak definition. The absence of a peak prevents estimation of genome size, and determination of whether a peak is haploid or diploid can be challenging. Finally, original estimates of genome size identified at the initiation of the project may not have been available for the exact species sequence, and instead be derived from nearby taxa, in some cases leading to significant differences between k-mer and previous genome size estimates.

##### *1.10 RNA sequencing*

RNA-seq was performed following standard protocols on an Illumina HiSeq 2000 platform. Briefly, poly- A+ messenger RNA (mRNA) was extracted from 1 mg total RNA using Oligo(dT)25 Dynabeads (Life Technologies, cat. no. 61002) followed by fragmentation of the mRNA by heat at 94 for 3 min (for samples with RNA Integrity Number (RIN) from 3–6) or 4 min (for samples with RIN of ≥6.0). First-strand complementary DNA (cDNA) was synthesized using the Superscript III reverse transcriptase (Life Technologies, cat. no. 18080-044) and purified using Agencourt RNAClean XP beads (Beckman Coulter, cat. no. A63987). During second-strand cDNA synthesis, deoxynucleoside triphosphate (dNTP) mix containing deoxyuridine triphosphate was used to introduce strand specificity. For Illumina paired-end library construction, the resultant cDNA was processed through end repair and A-tailing, ligated with Illumina PE adapters, and then digested with 10 units of uracil–DNA glycosylase (New England Biolabs, Ipswich, MA; cat. no. M0280L). Amplification of the libraries was performed for 13 PCR cycles using the Phusion High-Fidelity PCR Master Mix (New England Biolabs, cat. no. M0531L); 6-bp molecular barcodes were also incorporated during this PCR amplification. These libraries were then purified with Agencourt AMPure XP beads after each enzymatic reaction, and after quantification using the Agilent Bioanalyzer 2100 DNA Chip 7500 (cat. no. 5067-1506), libraries were pooled in equimolar amounts for sequencing. Sequencing was performed on Illumina HiSeq2000s, generating 100-bp paired-end reads. Table S4 shows the RNA sequence stats and NCBI accessions.

##### *1.11 Automated gene model annotation*

Twenty eight of the i5K pilot Version 1 genome assemblies were subjected to automatic gene annotation using a MAKER 2.0[10] annotation pipeline tuned for arthropods (Table S5). The two remaining assemblies for Mhra and Erow were considered too fragmented and incomplete to allow useful gene model annotation. The REDUNDANS software and new assemblies became available late in the project timeline, and thus automated gene annotations, orthologous gene family identification in OrthoDB and analysis were all performed on the Version 1 ALLPATHS-LG based assemblies.

The pipeline was designed to be systematic, providing a single consistent procedure for the species in the pilot study, scalable to handle 100s of genome assemblies, evidence guided using both protein and RNA-seq evidence to guide gene models and targeted to utilize extant information on arthropod gene sets. The core of the pipeline was a Maker 2 instance, modified slightly to enable efficient running on BCM-HGSC computational resources. The genome assemblies were first subjected to de novo repeat prediction and CEGMA analysis[11] to generate gene models for initial training of the ab initio gene predictors. Three rounds of training of the Augustus[12] and SNAP[13] gene predictors within Maker were used to bootstrap to a high-quality training set. Input protein data included 1 million peptides from a non-redundant (nr) reduction (90% identity) of Uniprot Ecdysozoa (1.25 million peptides) supplemented with proteomes from 18 additional species (*Strigamia maritima, Tetranychus urticae*, *Caenorhabditis elegans*, *Loa loa*, *Trichoplax adhaerens*, *Amphimedon queenslandica*, *Strongylocentrotus purpuratus*, *Nematostella vectensis*, *Branchiostoma floridae*, *Ciona intestinalis*, *Ciona savignyi*, *Homo sapiens*, *Mus musculus*, *Capitella teleta*, *Helobdella robusta*, *Crassostrea gigas*, *Lottia gigantea* and *Schistosoma mansoni*) leading to a final nr peptide evidence set of 1.03 million peptides. When available, species specific RNA-seq data either generated as described above, or culled from publically available sources was used judiciously to identify exon–intron boundaries, but with a heuristic script to identify and split erroneously joined gene models. Finally, the pipeline used a nine-way homology prediction with human, *Drosophila melanogaster*, and *Caenorhabditis elegans*, and InterPro Scan5 to allocate gene names. To assess the comprehensiveness of the gene models we identified a set of 1,977 arthropod specific CEGMA genes and assessed the number that could be identified in both the underlying assembly and the final automated gene model set. The automated gene sets, and all evidence tracks are available as gff from the BCM-HGSC website and as gff and browser interface at the National Agricultural Library i5K workspace (https://i5K.nal.usda.gov). Table S5 lists the automated gene model count, average transcript length, average CDS length, average protein length, total number of exons, average number of exons per gene, and the numbers and percentages of the 1,977 CEGMA genes found in the assemblies and final gene sets.

##### *1.12 Community gene curation and annotation*

Twenty-seven genomes were made available for manual annotations, and all genomes have had some level of manual annotation performed. Out of these twenty-seven, eleven genomes have completed an initial round of manual annotation. Annotation communities ranging from 9 to 41 individuals coalesced around each of these nine genomes. Community annotation was performed using the Web Apollo manual annotation software[14], which enables collaborative community curation of genome assemblies, and was hosted by the National Agricultural Library's i5K Workspace[15] (https://i5K.nal.usda.gov/). Annotators received training on manual annotation via webinars (AP for Ccap, MCMT for all others), and were instructed to adhere to a set of rules for manual annotation (https://i5K.nal.usda.gov/content/rules-web-apollo-annotation-i5K-pilot-project ). In some cases, additional RNA-seq tracks were supplied to support manual annotation inference. Community annotation efforts focused on gene and gene families relevant to each community's individual interests, rather than a set of universal annotations across i5K pilot species.

Annotation efforts were ‘frozen’ once each community completed their manual annotation efforts. After basic quality control of the manual annotations in GFF3 format via the NAL and the annotators, manual annotations were merged with the respective MAKER annotations into single, non-redundant ‘Official Gene Sets’ (OGS) (Agla, Clec, Ofas – via in-house scripts by DSTH; Ccap – by AP, used EVM to fold in several gene sets in addition to MAKER[16]; Ldec, Aros, Oabi, Focc, Hazt Gbue, and Bger – by MJMC and MFP (for details on the merge procedure, see https://github.com/NAL-i5K/I5KNAL_OGS/wiki/Merge-phase). Completed Official Gene Sets are available for download at the Ag Data Commons (links in Table S8). On average, 8.47% or ~1,400 genes were manually annotated for each OGS (Table S8).

##### *1.13 Gene model quality assessment*

Because the CEGMA gene annotation quality assessment tool is being discontinued, we additionally assessed the quality of the assemblies, gene sets and proteomes with BUSCO[17] and DOGMA[18]. For the assemblies BUSCO was used and the percentage of Complete BUSCOs, Single Copy BUSCOs, Duplicated BUSCOs, Fragmented BUSCOs and Missing BUSCOs is reported based on the BUSCO Arthropoda dataset (arthropoda_odb9) in Table S7. To compare the influence of assembly versions on the gene family and domain analysis the above mentioned BUSCO measurements are provided for the ALLPATH-LG and Redundans assemblies. The gene set quality for every species was determined with BUSCO based on the BUSCO Arthropoda dataset (arthropoda_odb9) as well and can be found in Table S7. For the domain analysis we used the domain-based quality assessment method DOGMA [18] to assess proteome completeness of every species used in this study. Completeness scores, based on the DOGMA eukaryotes coreset are provided in Table S7. For more information about the quality assessment methods BUSCO and DOGMA used here, please see the related publications.

##### *1.14 Orthology prediction*

Orthology delineation is a cornerstone of comparative genomics, offering qualified hypotheses on gene function by identifying “equivalent” genes in different species, as well as highlighting shared and unique genes that offer clues to understanding species diversity. The OrthoDB hierarchical catalog of orthologs (www.orthodb.org) offers the most comprehensive orthology resource for arthropod comparative genomics[19]. OrthoDB orthology delineation follows a multi-step process that is based on the clustering of best reciprocal hits (BRHs) of genes between all pairs of species. Clustering proceeds first by triangulating all BRHs and then subsequently adding in-paralogous groups and singletons to build clusters of orthologous genes. Each of these orthologous groups represent all descendants of a single gene present in the genome of the last common ancestor of all the species considered for clustering[20]. Thus orthology delineation is inherently hierarchical and sets of rather closely related species will generally produce finely-resolved orthologous groups (*i.e.*, with many one-to-one relationships) while sets that span large evolutionary distances will produce larger groups that capture the longer-term evolutionary histories of all the genes in the extant species.

The orthology datasets computed for the analyses of the 28 i5K pilot species together with existing sequenced and annotated arthropod genomes were compiled from OrthoDB v8[21], which comprises 87 arthropods and an additional 86 other metazoans (including 61 vertebrates). The OrthoDB v8 resource was compiled using available arthropod species in 2014 (hence some more recently published species are absent from our full set of 76 selected arthropods). Selection of species for inclusion in the 76-species dataset also required permission from data providers who at that stage had not yet published their results. Furthermore, species with particularly fragmented genome assemblies were excluded and only representative samples of the numerous available dipterans were selected as extensive studies on gene family evolution in *Drosophila* had already been published[22,23] and large-scale analyses were underway for the mosquitoes[24]. Although more than half of these genesets were built using MAKER, they were generated by different groups and with different levels of supporting data, and others were annotated using various different methods, and the amount of manual curation varies amongst the species from none to many years of extensive work (Table S6). This clearly introduces a potential source of technical variation rather than true biological variation in our analysis. Orthology clustering at OrthoDB included 10 of the i5K pilot species (*Anoplophora glabripennis, Athalia rosae, Ceratitis capitata, Cimex lectularius, Ephemera danica, Frankliniella occidentalis, Ladona fulva, Leptinotarsa decemlineata, Orussus abietinus, Trichogramma pretiosum*). The remaining 18 i5K pilot species were subsequently mapped to OrthoDB v8 orthologous groups at several major nodes of the metazoan phylogeny. Orthology mapping proceeds by the same steps as for BRH clustering, but where existing orthologous groups are only permitted to accept new members, *i.e.*, the genes from species being mapped are allowed to join existing groups when the BRH criteria are met. The resulting orthologous groups of clustered and mapped genes were filtered to select all groups with orthologs from at least two species from the full set of 76 arthropods, as well as retaining all orthologs from any of 13 selected outgroup species for a total of 47,281 metazoan groups with orthologs from 89 species. Mapping was also performed for the relevant species at the following nodes of the phylogeny: Arthropoda (38,195 groups, 76 species); Insecta (37,079 groups, 63 species); Holometabola (34,614 groups, 48 species); Arachnida (8,806 groups, 8 species); Hemiptera (8,692 groups, 7 species); Hymenoptera (21,148 groups, 24 species); Coleoptera (12,365 groups, 6 species); and Diptera (17,701, 14 species). All identified BRHs, protein sequence alignment results, and orthologous group classifications were made available for downstream analyses: http://ezmeta.unige.ch/i5K.

The i5K species together with those already available at OrthoDB extend genomic sampling to many different arthropod clades, even if overall sampling remains biased towards orders of particular interest with many more completed sequencing projects to date (Fig. S4 A). Partitioning Metazoa-level orthologs according to their presence and copy-number across the 76 arthropod and the outgroup species identifies a set of about 2,500 genes per species from orthologous groups that comprise single-copy orthologs from the majority of species (Fig. S4 B). A further 5,000-10,000 genes per species are widespread but are found in orthologous groups that have experienced more frequent gene duplications. These two major categories represent ancient genes that are evolving under ‘single-copy control’ or with a ‘multi-copy licence’[25]. The remaining genes belong to orthologous groups with less widespread species distribution, so-called ‘patchy’ orthologs that may be more frequently lost or lineage-restricted orthologs that may have emerged recently within the lineage or have simply diverged beyond recognition, *e.g.*, those that appear as arthropod-restricted orthologous groups for which no confident orthologs could be found beyond Arthropoda. Finally, each species exhibits a fraction of genes for which no confident orthologs could be identified.

##### *1.15 Phylogeny inference*

The arthropod phylogeny serves as the framework in which we can ask questions about species relationships, evolutionary rate, gene families, and protein domains among our 76 species. We aimed to utilize as much information from the 38,195 orthogroups as possible for reconstruction of the arthropod phylogeny, while minimizing the effect of gene-tree incongruence. The general strategy to accomplish this is to select orthogroups that are represented by a single gene in all taxa in the dataset, thus minimizing incongruence due to gene duplication. These orthogroups are often referred to as single-copy genes or one-to-one orthologs. However, as the number of taxa in a dataset increases the number of single-copy genes tends to decrease because of gene loss or mis-annotation of genes in any given lineage. Because of this trade-off between number of species and number of single-copy genes, we found no single-copy genes in all species of our species rich dataset. Three gene families (EOG8BZQH7, EOG8BZQK0, EOG8DFS3J) are single-copy in all species except for one.

This lack of single-copy orthologs necessitates a different strategy for species tree reconstruction. We decided to first make a backbone tree at the order level and then make separate trees for the six orders comprised of multiple species in our dataset. These separate order trees can then be pasted on to the backbone tree. This allows us to maximize the amount of data for most of our species while still being able to infer the placement of more ancestral nodes in the tree. Constructing the backbone tree required us to change our perspective of what we call ‘single-copy’ genes. In this case, we want gene families that are represented by at least one species in each order with a single copy. This means that the 15 single species orders in our dataset must be single copy in every orthogroup we choose, but the 6 multi-species orders can be represented by different combinations of species for each gene, so long as they are single-copy. We found 150 such single-copy genes at the order level, with varying species present between them such that we were able to infer the full 76 species phylogeny, but with a high amount of missing data. For the six multi-species order trees we simply counted genes that were single-copy in all species in that order (Table S13). Additionally, we constructed a separate phylogeny of the Chelicerates and Crustaceans based on 569 and 1,107 single-copy genes, respectively. Branch lengths from these group phylogenies were preserved across divergence time and evolutionary rate analyses, ensuring that the maximum amount of data was used. Inferences from the 150 single copy backbone genes were restricted to only a few species and deep internal nodes (Table S13). Then once we have the full 76 species backbone phylogeny based off of 150 genes in hand, we can simply replace the topology and branch lengths of the 6 multi-species order and Chelicerate and Crustacea sub-trees with those estimated from hundreds to thousands of genes.

Branch lengths for the entire phylogeny were estimated from concatenated alignments of the dataset that maximized the number of orthologs used for a given lineage. For example, we found 569 single-copy orthlogs among Chelicerates and used a concatenated alignment of these genes to estimate their branch lengths in the larger Arthropod phylogeny. However, among spiders (a subgroup of Chelicerates) we found 1,627 single-copy orthlogs from which we estimated branch lengths for the larger phylogeny. This was done for all groups mentioned in Figure S11 and remaining nodes had branch lengths estimated from the 150 single-copy backbone genes. Groups were chosen based on taxonomic classifications, mostly being orders represented by multiple species in our data, with the exception of Chelicerates and Crustacea which are both subphyla. This nesting of phylogenetic analyses ensured that we maximized the amount of data used for each lineage, though branch lengths in the final phylogeny do not vary drastically whether using only the 150 single-copy backbone genes for all species or the maximized number of single-copy genes among groups ($r^2$ = 0.86, Fig. S31)

For downstream analyses, rooted trees are required. The backbone tree was simply rooted at the mid-point of the branch between chelicerates and mandibulates. For each multi-species order gene tree we required an outgroup species to be present so we could root the species trees after their inferences. For outgroup selection we ranked every species outside the order based on whether it was single copy in a given family. For example, given the 3,932 one-to-one orthologs in the order Coleoptera, *Pediculus humanus* from the order Psocodea is present as single copy in 3,880 and was chosen as the outgroup for those families. For the remaining 52 groups the second most common single-copy outgroup was used, though we simply labeled the outgroup taxa as "Outgroup" in our alignments.

To ensure that our inferences were not influenced by the method of reconstruction we employed two alignment methods (MUSCLE and PASTA) along with 3 species tree reconstruction methods (concatenation with RAxML, average consensus implemented in SDM, and the coalescent quartet method ASTRAL) for inferring our trees. This resulted in 6 species trees for each multi-species order and the backbone tree which we could compare to one another. Two of these species tree methods (average consensus and ASTRAL) require gene trees, which we used RAxML to infer. All reported trees are run with the PROTGAMMAJTTF amino acid substitution model; however the trees were robust to changes in the specified model.

Because of the consistency of results between methods (See Main Text), we used species trees inferred from alignments with PASTA and species tree inference with ASTRAL as our final set of trees. Because ASTRAL version 2.0 does not estimate branch lengths we were forced to use the ASTRAL topology as a constraint tree in RAxML with the concatenated alignment to estimate branch lengths. This means that, while the topology inferred by ASTRAL should be robust to incomplete lineage sorting (ILS), the branch lengths and estimated number of substitutions may be inflated because of SPILS (Substitutions Produced by ILS[26]). After estimating branch lengths on the 6 multi-species order trees and the backbone tree and then pasting the 6 multi-order trees onto the backbone tree, we were left with a single tree to use in all subsequent analyses (Fig. 2).

##### *1.16 Phylogeny inference – an additional point brought up during review.*

During the review process, a reviewer brought up the low number of universal single-copy orthologs identified, especially when compared with Misof et al.[3] Our response is placed here for readers with the same question:

We were also disappointed with the low number of universal single-copy orthologs identified, however, detailed analysis of our orthology data explains this low recovery rate: (i) even having just a few missing orthologs from each additional species considered gradually reduces the total number of universal orthologs, but this reduction is much more dramatic when filtering for duplicated orthologs i.e. (ii) duplications in each additional species considered are more frequent and result in more discarded orthologous groups. We examined Misof et al in detail to understand the differences: Misof et al identified 1478 single-copy orthologs from 12 reference species with genomes (not the full set of their 103 transcriptome species), then they searched for and identified, on average, 98% of these genes in the 103 de novo sequenced transcriptomes, and they recovered fewer from their outgroup non-insect arthropods. Using the same 12 reference species from our dataset we identify 3621 single-copy-in-all orthologs, on average, 96% of these are found in our other 53 insects, and 89% in our other 11 non-insect arthropods. I.e. a very similar result to that from the Misof et al study, which used HaMStRad and "chose the single best or the best set of non-overlapping transcripts in case multiple transcripts had been assigned to a given OG (option -representative).". In our analysis however, when applying the additional filter of requiring the identified orthologs to be single-copy then retained orthologous groups drop to, on average, 76% for the insects and 59% for the non-insect arthropods. Thus gene duplications, especially in species where this has been extensive like the cockroach, mean that the strict criterion of single-copy-in-all results in very few such orthologs across the entire phylogeny. We have added these findings to the supplement as they do indeed explain what some readers might at first interpret as a worrying inconsistency between our study and that of Misof et al. In addition we have edited the text and figure legend to clarify that only the backbone phylogeny relied on this low number (150) of strictly universal-single-copy orthologs, the separate multi-species order phylogenies used many more (from 1331 to 4097, as given in Table S13 single copy orthologs column).

##### *1.17 Divergence time estimation*

The methods of phylogeny inference outlined above result in a tree with branch lengths in terms of relative number of substitutions or coalescent units. In order to study rates of evolution and to reconstruct ancestral gene counts, branch lengths in terms of absolute time are required. We used a simple non-parametric method of tree smoothing implemented in the software r8s[27] to estimate these divergence times. r8s was chosen for its speed on large datasets and because we were able to utilize all data for branch length calculations across the whole tree while Bayesian methods would require us to re-estimate branch lengths from only the 150 backbone genes. We also observe consistency between r8s and more complex Bayesian models. To examine this, we compared divergence times estimated by r8s to those estimated by MCMCtree[28] and found a strong correlation ($r^2 = 0.91$). Fossil calibrations are also required to scale the smoothed tree by absolute time. We relied on the aggregation of deep arthropod fossils done by Wolfe et al.[29] along with a few more recent fossils used by Misof et al.[3] (Table S14).

Because r8s does not provide a confidence interval on divergence times, we employed a jackknife approach on fossil calibration points to generate a distribution of ages around each node. We randomly removed 5 fossil calibration points and re-estimated divergence times and repeated this process 100 times. We find that the original divergence time estimate for each node always falls within 1 standard deviation of the mean of the distribution generated by the replicate process (Fig S33). In other words, the removal of any fossil calibration point has very little effect on inferred node ages and that node ages are very stable with regards to the choice of fossils used to calibrate the tree.

##### *1.18 Substitution rate estimation*

To estimate substitution rates per year on each lineage of the arthropod phylogeny we simply divided the expected number of substitutions (the branch lengths in the unsmoothed tree) by the estimated divergence times (the branch lengths in the smoothed tree) (Fig. 4).

##### *1.19 Gene family analysis*

With the 38,195 orthogroups and ultrametric phylogeny we were able to perform the largest gene family analysis of any group of taxa to date. In this analysis we were able to estimate gene turnover rates ($\lambda$) for the six multi-species orders, infer ancestral gene counts for each family on each node of the tree, and estimate gene gain and loss rates for each lineage of the arthropod phylogeny. The size of the dataset and the depth of the tree required several methods to be utilized.

Gene turnover rates (λ) for the six multi-species orders were estimated with CAFE 3.0, a likelihood method for gene-family analysis[30]. CAFÉ 3.0 is able to estimate the amount of assembly and annotation error (ε) present in the input gene count data. This is done by treating the observed gene family counts as distributions rather than certain observations. CAFE can then be run repeatedly on the input data while varying these error distributions to calculate a pseudo-likelihood score for each one. The error model that is obtained as the minimum score after such a search is then used by CAFE to obtain a more accurate estimate of λ and reconstruct ancestral gene counts throughout the tree (Table S12). However, with such deep divergence times of some orders, estimates of ε may not be accurate. CAFE has a built-in method to assess significance of changes along a lineage given an estimated λ and this was used to identify rapidly evolving families within each order. We partitioned the full dataset of 38,195 orthogroups for each order such that taxa not in the order were excluded for each family and only families that had genes in a given order were included in the analysis. This led to the counts of gene families seen in Table S12.

For nodes with deeper divergence times across Arthropoda, likelihood methods to reconstruct ancestral gene counts such as CAFE become inaccurate. Instead, a parsimony method was used to infer these gene counts across all 38,195 orthogroups[31]. Parsimony methods for gene family analysis do not include ways to assess significant changes in gene family size along a lineage, so we performed a simple statistical test for each branch to assess significance. Under a stochastic birth-death process of gene family evolution, within a given family the expected relationship between any node and its direct ancestor is that no change will have occurred. Therefore, we took all differences between nodes and their direct descendants in a family and compared them to a one-to-one linear regression. If any of the points differ from this one-to-one line by more than two standard deviations of the variance within the family it is considered a significant change and that family is rapidly evolving along that lineage. Rates of gene gain and loss per year were estimated in a similar fashion to substitution rates per year. We simply counted the number of gene families that are inferred to be changing along each lineage and divided that by the estimated divergence time of that lineage (Fig. 4).

Finally, to estimate ancestral gene content (*i.e.*, the number of genes at any given node in the tree), we had to correct for gene losses that are impossible to infer given the present data. To do this, we first regressed the number of genes at each internal node with the split time of that node and noticed the expected negative correlation of gene count and time (Fig. S5) ($r^2$=0.37; $P$=4.1 x $10^{-9}$). We then took the predicted value at time 0 (present day) as the number of expected genes if no unobserved gene loss occurs along any lineage and shifted the gene count of each node so that the residuals from the regression matched the residuals of the 0 value.

##### *1.20 Jackknifing tests on parsimony reconstructions of gene families*

In order to assess the impact of any single lineage on the parsimony reconstructions of ancestral states of gene families and protein domains we performed a series of jackknifing tests.

We randomly removed 5 of the 76 species from our dataset and re-ran the parsimony reconstructions to get new estimates for the number of genes gained and lost and the number of protein domain rearrangements. This process was repeated 100 times to generate a distribution of events (Figs. S32 and S34 for gene family and protein domain events, respectively). In this manner, we can ensure that our conclusions from the full dataset are not overly influenced by any error-prone assemblies. We find that, regardless of the species removed, the total number of gene family changes we observe with the full data always falls within one standard deviation of the mean observed from 100 replicates with randomly removed species. Paying special attention to known low quality genomes, *Latrodectus hesperus* (LHESP), *Loxosveles recules* (LRECL), and *Limnephilus lunatus* (LLUNA), we find that their removal only impacts reconstructions on very closely related lineages. For example, the removal of the two spiders means we infer many more gene family changes in *Stegodyphus mimosarum* (SMIMO), and many fewer in *Centruroides sculpturatus* (CSCUL; Fig S31). Among protein domains, the removal of LHESP, LRECL and LLUNA affects only the estimated Single Domain Losses, which drop around 2% (leading to a slight percental increase of all other events; Fig S34). This suggests that missing domains due to low quality of assemblies leads to an overestimation of losses in these species, while all other event types are very robust to quality differences or sampling effects. Since this overestimation does not affect other nodes in the tree and is limited to the event type of Single Domain Loss, including these species in the analysis still adds a lot of information, while introducing just a minor bias. This bias can be taken into account by not over-interpreting Single Domain Losses in low-quality species (for quality scores see Table S7).

##### *1.21 GO enrichment tests*

GO enrichment tests were done with the R v3.4.2[32] package "GOstats"[33] using a hyper-geometric test. For each node, background gene sets were composed of every gene present at that node and every gene that was inferred to have gone extinct along the lineage leading to that node. Test sets consisted of each category of gene family change: gain, loss, emergence, and extinction. Any gene without a known GO term was removed from the GO analysis.

##### *1.22 Protein Domain evolution analysis*

Proteins are the workhorse of the cell and, at the same time, the most basic devices which can be phenotypically related to selection and adaptation. Since biologists in general, and geneticists in particular, have traditionally related genes to observable traits, and genome sequences are available in large quantities, many evolutionary studies approximated growth and birth-death processes of proteins and protein families by analyzing the analogous processes of their underlying genes.

However, the further towards the root of a tree we try to reconstruct the evolutionary history of proteins, the more we need to consider that a major mode of protein evolution results from protein domains changing their combination and order within a protein. Protein domains can be seen as the functional and structural building blocks of proteins[34,35]. Some proteins contain just one single domain, while others consist of multiple domains and sometimes contain consecutive copies of domains, so called tandem repeats. The N- to C-terminal order of domains in the amino acid sequence defines a domain arrangement[36].

While the majority of proteins in large clades of organisms are composed of a relatively small set of domains, the number of unique arrangements is tremendous and long domain arrangements are often species-specific[37]. After the occasional formation of a novel domain, it can rapidly be recombined in different arrangements and functional contexts, and in this way explain the vast protein diversity we can observe in nature[38-40]. Much of this molecular biodiversity is often overlooked because orthology detection methods by construction need to compromise between the accurate capturing of many fragments of family members and the comprehensive representation of a few.

Protein domains are defined in databases such as Pfam[41] (based on sequence similarity) or Superfamily[42] (based on structure similarity), which provide a highly standardized way to study proteome evolution based on domains and their arrangements.

We want to assess and evaluate the evolutionary paths and patterns of modular domain rearrangements at a high resolution and determine the rates (*e.g.*, in terms of events along a lineage or in a genome per million year). Previous research has demonstrated that it is not necessary to postulate concepts such as recombination or ”domain shuffling” to explain the vast majority of events. In fact, there are just four biological events that can explain the coming about of virtually all domain arrangements: fusion of existing arrangements (also of single domain proteins; this amounts to gene fusion), fission of existing domain arrangements, terminal loss of one or more domains (*i.e.*, there are no traces left as the underlying DNA sequence *e.g.*, is no longer transcribed) and terminal gain of one domain. This was found to be the case for a set of 29 plants dating back as far as 800 mya and 20 pancrustaceans available and dating back ca. 430 mya[37,43].

Since ca 40% of all domain rearrangement events could so far not be accurately classified and the classified ones were in total number just a few thousand, we created an improved method to classify a significantly larger number of events. We do so by adding two additional event types in our analysis: single domain loss and single domain emergence (Fig S6). These can be seen as conceptual event types, since they can be explained by the underlying mechanisms for terminal losses and gains, but have not been (separately) addressed in the studies before. Single domain gains refer here to the first emergence of a domain in a phylogenetic tree. Likewise, a terminal emergence is also just considered as such, if the added domain emerged newly at this node in the tree.

We annotated proteomes for all 76 arthropod species and 13 outgroup species with Pfam domains in version 30 using the PfamScan.pl script provided by Pfam[41]. To prevent evaluating different isoforms of proteins as additional rearrangement events, we removed all but the longest isoform. Repeats of a same domain were collapsed to one instance of the domain (A-B-B-B-C → A-B-C), since copy numbers of repeated domains can vary strongly even between closely related species[44,45].

Reconstruction of ancestral protein domain states was carried out in C++ using our DomRates tool (http://domainworld.uni-muenster.de/programs/domrates/). All ancestral states for full arrangements were reconstructed with Fitch parsimony and in a separate run using Dollo parsimony for all single domains. The inferred single domain states were then used for reconstruction of all terminal loss/emergence and single domain loss/emergence events. This approach differs from previous studies in the usage of both Dollo and Fitch parsimony to determine separately the ancestral states of single domains and arrangements. While a Fitch parsimony principle models ancestral states for arrangements much better than Dollo parsimony, this method was previously also applied to all single domains. However, since we expected domains to emerge only once, the ancestral presence/absence states of single domains are best modeled by Dollo parsimony[46].

Based on these reconstructed ancestral states for all inner nodes in the phylogeny (Fig. S6 A) every node was compared to its ancestor (Fig. S6 B). If the presence/absence state of an arrangement changed compared to an ancestor, we checked if we could explain it with one of the following six event types (Fig. S6 C): (1) fusion, if a gained arrangement can be explained by a fusion of two arrangements from the direct ancestor: (2) fission, if an ancestral arrangement can be split to form with one part the newly gained arrangement in the descendant and the second part is also still present; (3) terminal loss, if just one split product exists in the descendant; (4) terminal emergence, if a single domain that was not present in the ancestor, was added at the C- or N-terminal end of an arrangement; (5) single domain loss, if a domain that existed in a single state in the ancestor is not present in the descendant anymore, either alone or as part of an arrangement; (6) single domain emergence, if a new domain emerges in a single state and was not present in the phylogenetic tree before.

In this study we just considered all events for the rate calculation that could be unambiguously explained by one of the above defined event types in a single step. Arrangements and single domains specific to the outgroup species were not taken into account for rate calculation.

#### 2. Supplemental Results

##### *2.01 DNA methylation across the arthropods*

Epigenetic inheritance plays a fundamentally important role in mediating gene regulation[47] and DNA methylation is one of the most widespread forms of epigenetic information[48] implicated in several important biological processes[49]. However, the function of DNA methylation in arthropods remains unclear. We analyzed the genomes of 76 species of arthropods in order to understand the evolution of DNA methylation.

We queried the genomes to determine copy number of the two putative DNA methyltransferases, DNMT1 and DNMT3, as well as the DNA-methylation-associated genes MBD2 and UHRF1. Gene existence was evaluated by using phylogenetically distributed animal orthologs to query each species' gene set and genome. We first scanned all the translated coding sequences using blastp with a cutoff of 1e-20. In order to avoid counting fragmented models of the same gene twice, we merged nonoverlapping significant hits. We then scanned the genomes of each species using the same representative orthologs using tblastn and an initial e-value cutoff of < 1e-5. We merged adjacent hits (those without a gene model between them on the same strand) from the same scaffold that did not overlap in their identity to our query orthologs, taking the lowest e-value as the "gene-level" e-value. We then added up the number of amino acids with matches across all such hits and filtered these composite genes by requiring that the composite gene had an e-value < 1e-10 and had at least 150 matched amino acids across the whole gene. Finally, we compared the copy counts from the coding sequence scan to those from the genome scan, and manually inspected all those that did not match between the two to arrive at our final estimated gene copy number.

We then analyzed patterns of CpG dinucleotide depletion in the genomes in order to investigate putative patterns of CpG DNA methylation. In principle, if a genomic feature is methylated, then the observed frequency of CpG dinucleotides should be lower than that expected based on frequency of C and G nucleotides due to an increased rate of mutation from methylated C's to T (*i.e.*, CpG o/e will be less than 1.0). However, if a genomic feature is un-methylated then the observed frequency of CpG dinucleotides should equal that expected (*i.e.*, CpG o/e will equal 1.0). As described in Simola et al[50], normalized CpG dinucleotide content was calculated as CpG o/e = ($length^2$/length)*(CpG count/(C count * G count)).

We find that the molecular machinery putatively involved in DNA methylation shows a patchy distribution among the arthropods. A putative DNMT1 ortholog was identified in the majority of taxa examined (Fig. S7). However, DNMT1 is not present in the Diptera. DNMT3 seems to be less well-conserved and could not be confidently identified in several of the Holometabola. For example, DNMT3 is apparently not found in the Diptera, Lepidoptera, and many, but not all, Coleoptera. DNMT3 identity was also less well conserved in the non-insect arthropods analyzed in this study[51]. Interestingly, many taxa exhibited strong evidence of genomic methylation (i.e., strongly bimodal CpG o/e distribution, distinct high- and low-CpG o/e profiles across genes, etc.), despite the apparent loss of DNMT3. This may indicate that DNMT3 is a less fundamental component of the insect DNA methylation toolkit than previously assumed. The presence of UHRF1 generally coincided with the presence of DNMT1, as expected given the function of the Uhrf1 in recruiting Dnmt1 in cells. In contrast, MBD2 appears to be present in most arthropod taxa, which is not surprising given that Mbd2 is thought to have functions beyond those associated with DNA methylation. Thus, overall, there appear to be several independent losses of DNA methylation machinery with associated changes in genic patterns and levels of CpG depletion across arthropods (e.g. MOCCI -- mites, LREC – chelicerates, PVEN -- psylid, DPOND – beetle), and the presence of the DNMT3 genes is surprisingly variable.

Average levels of DNA methylation, as judged by CpG o/e, showed remarkable variation across the Arthropoda (Fig. S7). The holometabolous insects tended to show high levels of CpG o/e and, therefore, lowest levels of inferred DNA methylation. In contrast, the hemimetabolous insects and non-insect arthropods tended to show higher levels of DNA methylation than the holometabolous insects[51,52]. However, there were some species outside the Holometabola, such as *Pediculus humanus*, *Strigamia maritima*, and *Metaseiulus occidentalis*, that did not show strong evidence of DNA methylation in their genomes based on average genic CpG o/e levels [52]. Thus DNA methylation seems to have been lost or diminished in the Holometabola and in a few other non-Holometabola arthropod taxa. Such a trend runs counter to the prediction that DNA methylation contributes strongly to developmental plasticity[52,53], which tends to be more well developed in holometabolous insects. Overall, these results indicate that methylation levels were highest in ancestral arthropod taxa, and have remained relatively high in most of the basal arthropods and hemimetabolous insects, but have dropped substantially in the holometabolous insects. In addition, several arthropod taxa have apparently lost the ability to methylate their genomes.

Different arthropod clades show large differences in CpG o/e profiles indicating differences in patterns of methylation among species (Fig. S8). Bimodal distributions, when present, reflect that genes fall into two discrete groups which are either methylated or un-methylated. Some basal arthropods, such as the Araneae, show a clear signal of bimodal patterns of genic methylation, whereby some genes are relatively highly methylated and others are not. Finally, some taxa, particularly in the Hymenoptera, show levels of CpG o/e higher than expected by chance alone, which may be a signal of the strong effects of gene conversion in the genome[54].

Exons often show the strongest evidence of CpG depletion in Holometabola, suggesting that exons are the most highly methylated regions of the genome (Fig. S8). However, introns and exons both show relatively high levels of CpG depletion in more basal taxa in the Arthropoda indicating that methylation is more uniform across genes in these species (Fig. S8). In addition, many of the Holometabola appear to have DNA methylation restricted to the 5' region of genes, while the Hemimetabola and more basal taxa appear to have methylation throughout gene bodies. The Coleoptera and Diptera showed fairly unimodal distributions of CpG o/e suggesting an absent or diminished DNA methylation system.

We conclude that DNA methylation in the arthropods shows remarkable variation. The non-insect arthropods and hemimetabolous insects have high levels of DNA methylation whereas the holometabolous insects have low levels of DNA methylation. There appear to be several independent losses of DNA methylation machinery. In addition, patterns of DNA methylation vary considerably among taxa indicating that the effects of DNA methylation on gene function may differ across species. Thus, arthropods may be a very good taxon in which to study the causes and consequences of DNA methylation.

##### *2.02 Pancrustacea phylogeny*

Recent large-scale phylogenetic analyses have suggested that the subphylum "Crustacea" is paraphyletic, with Hexapoda as an in-group, sister to the branchiopods (Cladocera), thus forming a monophyletic Pancrustacea. This grouping is widely accepted[55-57], even though it is often accompanied by low statistical support, suggesting a more critical analysis is required. The tree shown in Fig. 2 indicates a monophyletic crustacean group sister to the hexapods, though with low bootstrap support of the nodes involved in the presumed paraphyly. We find that the topology of the three crustacean orders is the most contentious in our phylogeny. For the four branches of interest (the three crustacean orders and the branch leading to Hexapoda), there are a total of 15 possible topologies (Fig. S9). Among our six tree construction methods, we recover four different topologies.

To investigate the crustacean topology further, we decreased the number of species in our phylogenetic reconstruction in order to increase the number of sequences we could use to infer the topology of these lineages. We used three datasets including the original 22 orders with 150 genes (Dataset 1). Then we decreased the number of orders from 22 to 17 which allowed us to identify 408 single-copy orthologs (Dataset 2). Finally, we reduced the number of orders to 12 leading to 1,107 single-copy orthologs (Dataset 3). We also tested different amino acid models along with our original framework of two alignment methods and three species tree methods. All together, we reconstructed 24 species trees for each dataset, resulting in 72 total species trees.

In total, we recovered 12 of the 15 possible topologies (Fig. S9), and half of our methods support a monophyletic crustacean clade. We find no clear source for systematic error in the discordance between species tree reconstruction methods. Changing the alignment software, amino acid model, or species tree method does not seem to affect the clarity with which this topology is inferred. In Datasets 1 and 2 there does seem to be a tendency for concatenation/consensus methods to support the monophyletic grouping of the crustacean orders (topology 3 in Fig. S9) while coalescent methods result in non-monophyletic groupings. However, the coalescent method itself seems sensitive to changes in alignment method and amino acid model. In general, a monophyletic Crustacea seems to be supported better than a non-monophyletic one, with average bootstrap support of the relevant nodes being 83.27 and 51.10, respectively, with monophyletic topology 3 being recovered 31 out of 72 times.

The sensitivity to small changes in inference methods and low statistical support of crustacean nodes could point to true biological discordance among gene trees. Gene duplication and loss, introgression, and incomplete lineage sorting are all sources of such discordance. Gene tree discordance can lead mis-inferences of mutational events along a resulting species tree. In light of our whole genome data and the resulting disagreement between phylogeny inference methods regarding Crustacea, we conclude that the relationships between the crustacean orders and their insect sister species remain unresolved. As such we advise and practice caution when drawing conclusions about evolutionary events in these lineages. The major drawback of this work is the sampling of only three crustacean species. The addition of other crustaceans with whole genome data may increase confidence in the Crustacea-Hexapoda relationships in the future.

##### *2.03 Gene families evolving on the most lineages*

The identification of rapidly evolving gene families can provide insight into the molecular and biological functions that arose on different branches of the arthropod tree. We identified the 30 gene families that were inferred to be rapidly changing along the most lineages (Table S15). After excluding transposable element families that showed large changes in size (supplement), these gene families include cytochrome 450s involved in xenobiotic defense, digestive enzymes, non-p450 detoxification enzymes, cuticle components and enzymes involved in molting, zinc finger transcription factors, and genes involved in fatty acid metabolism possibly for energy production for flight, and chemosensation. With the exception of the zinc finger transcriptases, all these genes are likely related to the day-to-day survival of arthropods across many hostile environments.

##### *2.04 Coleopteran gene family evolution summary*

Fourteen gene families, together containing 707 genes in the 4 species of beetles studied, showed large expansions compared to other arthropods. Of these, EOG8Z91BB, which contains genes encoding proteins with a Ribonuclease H-like domain, had the most genes (105). These genes are putatively involved in nucleic acid metabolism, including DNA repair and replication, and RNA interference. EOG8CRPGZ, which contains PiggyBac transposable element-derived proteins, and EOG8KWN7R, which comprises proteins with putative transposase activities, are notably expanded in both *A. glabripennis* (37 and 25 genes, respectively) and *L. decemlineata* (52 and 26 genes, respectively), suggesting that these phytophagous beetle genomes have high levels of transposable element activity. *Leptinotarsa* notably has 31 genes in EOG8GTNT0 (integrase catalytic domain; important for integrating viral genomes into host genomes), which are typically found in proteins with transposase activities. EOG8WDGTK was expanded in *A. glabripennis* (37 genes) and contains proteins with Parvovirus coat VP1. Four families that contain genes encoding proteins with potential transmembrane transporter activity (EOG8284G8, EOG80P6ND, EOG80S2WJ, EOG89W4VJ) show a similar degree of expansion in the beetles studied. McKenna[58] proposed that the acquisition of new genes via HGT, followed by gene copy number amplification and functional divergence, contributed to the addition, expansion, and enhancement of the metabolic repertoire of beetles, perhaps especially those that feed on living plants. Our results are compatible with this view, and further reveal that selfish DNA, including transposable elements and viruses, may also play a key role in the generation of adaptive novelty (metabolic and other) in beetle genomes. Notably, all beetle genomes studied to date have been from the species rich suborder Polyphaga (more than 350,000 described extant species). The other 3 suborders of beetles (Adephaga, Archostemata, and Myxophaga), remain un-sampled.

***2.05 Diptera gene family evolution summary***

Gene family evolution analyses in flies are particularly attractive given the large number of functionally characterized genes in *D. melanogaster*. Gene family expansions in the lineage to *Drosophila*, for instance, have the potential to inform about adaptive events in the more recent history of *Drosophila* evolution likely with higher accuracy and efficiency given the increased accuracy of orthology calls, thus leveraging the phylogenetic closeness of *Drosophila* genetic data. Vice versa, also insights into family extinctions in the lineage leading to *Drosophila* are of exceptional use as this information informs about gene families that cannot be studied in *Drosophila*, hence calling for studies in arthropod/insect satellite model species. Finally, previously documented gene family expansions and extinctions in flies can serve as benchmarks/gold standard to characterize the accuracy and sensitivity of genome-wide gene family studies.

The dipteran portion of our gene content analysis tree only captures a small glimpse of fly diversity, which is documented to exceed 150,000 species[59]. Moreover, the majority of the corresponding 14 sampled dipteran species are comparatively closely related with 4 species representing mosquitoes and 3 species representing the genus *Drosophila*. At the same time, given the distant relation of these two preferentially sampled clades, the taxon sample covers close to the maximum of dipteran species divergence which reaches back approximately 170 million years[3].

In total, the dipteran i5K subtree contains 15,424 expansions and 43,539 contractions, suggesting that gene loss has been more frequent than gain (Fig. S29 A & B). In part, however, this indicated trend could be an artifact due to poor gene coverage in the sand fly and Hessian fly genomes, which stand out by accounting for 12,905, *i.e.*, 30%, of the gene family contractions (Fig. S30 B). Among the internal nodes, the root note and the close to terminal note DL 1, which represents the presumptive last common ancestor to the mosquito species *Aedes aegypti* and

*Culex quinquefasciatus*, stand out with the highest number of gene family expansions, 709 and 700 respectively. Only DL 1, however, also stands out with the highest number of rapidly evolving gene families, *i.e.*, 38, while the dipteran root node accounts for only 4 rapidly evolving gene families, ranking among the bottom 4 internal branches in this variable (Fig. S30 A & B). In general, however, there is also a pronounced bias in the taxonomic representation of the 889 dipteran rapidly evolving gene families: 148 are associated with 13 internal nodes (1.5%) while 741 (84%) are associated with 14 OTUs (Fig. S30 C). In the latter, remarkably the house fly *Musca domestica* accounts for 203 (23%) rapidly evolving gene families alone compared to an average of 41 in the other OTUs.

Dipteran relationships in the i5K tree are largely consistent with the flytree of life[59], except for the closer relatedness of the Mediterranean fruitfly, a representative of the family Tephritidae to the Drosophilidae instead of calyptrate Diptera.

Rapidly evolving gene family content was analyzed in more detail for 4 important, consistently recovered nodes. This includes the dipteran root node DL 48 and two higher up nodes that are associated with major radiations in the dipteran tree of life. One of them is the node DL 17 at the base of schizophoran fly diversity and one of them is DL 13 at the base of calyptrate diversity (Tables S22, S26, and at https://i5k.gitlab.io/ArthroFam/). The dipteran root node is characterized by two rapidly expanding gene families, both of which include functionally characterized *Drosophila* genes: The gene family of *grauzone* (*grau*) related zinc finger transcription factors, which are involved in the regulation of oogenesis[60], and the Niemann-Pick type C-2d gene family of sterol transport involved lipid associated proteins (Table S23, worksheet DL48 Dipteran Root). These two expansions at the dipteran root are complemented with three diagnosed contractions, two of which, however, concern the same gene family of Activity-regulated cytoskeleton associated (Arc) proteins (Table S23, worksheet 'DL48 Dipteran Root').

The approximately 65 million years old higher taxon Schizophora represents one of the most dramatic radiations within Diptera, with over 50,000 species[59]. CAFE analysis detected 14 rapidly evolving gene families at the base of this group, the large majority of which (12) have been expanding (Table S25, worksheet 'DL17 Schizophora'). The majority of genes within these gene families lack functional characterization in *Drosophila*. Notable exceptions include the Lysozyme gene family, which has been previously reported for having experienced an adaptive expansion in flies[61], the Serine proteinase inhibitor gene family[62], the Halo-related gene family of Halo kinesin cofactors of lipid droplet trafficking protein complexes (ref), the 64a subfamily of gustatory receptors for sugar taste[63], and the pickpocket family of Na+ channel proteins[64]. It will thus be interesting to explore the role of these family expansions in the staggering diversification of schizophoran Diptera.

By comparison to the schizophoran node DL 17, the Calyptrate-specific node DL 13 only highlights a much smaller number of gene family expansions, *i.e.*, 3 (Table S26, worksheet 'DL13 Calyptrates'). This number is strikingly small give that calyptrate Diptera represent approximately 50% of the vast diversity of schizophoran Diptera. Further notable, all of the three gene families are missing from *Drosophila*, thus lacking functional annotation. Two of them code for zinc finger transcription factors, while the third codes for a protein with Ribonuclease H-like, DNA polymerase, and Recombination endonuclease VII domains. No specific studies exist yet on the role of these gene families in calyptrate evolution.

As a first attempt to utilize previously documented fly-specific gene family expansions to characterize the accuracy and sensitivity of the CAFE i5K tree results, we explored whether the data recovers the dramatic expansion of Methuselah/Methuselah-like G-protein coupled receptor (GPCR) family during recent *Drosophila* evolution[65]. While drosophilids possess 10 to over 15 members of this gene family, distantly related Diptera possess on average 5 due to recent expansions preceding and within the genus *Drosophila*[66,67]. None of 74 OrthoDB Methuselah/Methuselah-like orthology groups is associated with significant expansions or contractions in the CAFE i5K tree results, suggesting that future analyses of the i5K gene family tree are poised to detect larger numbers of clade-specific gene family changes.

##### *2.06 Protein domain analysis*

A complementary analysis to gene families is the reconstruction of ancestral protein domain content. Based on the ancestral domain content we can then infer all domain rearrangement events and measure domain emergence and loss rates. These types of changes define the biochemical and cellular capacity of proteins encoded by the gene family changes identified above, and the emergence of a completely novel protein domain can be an important evolutionary event. In total we can explain the underlying events for more than 40,000 domain arrangement changes within the arthropods (Tables S3.02 and S3.03 this document, and Fig S21). The majority of novel arrangements (48% of all observable events) were formed by a fusion of two ancestral arrangements, while the fission of an existing arrangement into two new ones accounts for only 14% of all changes. Comparison of emergence and loss rates shows another interesting signal: 37% of observable changes can be explained by losses (either as part of an arrangement (14%) or the complete loss of a domain in a proteome (23%)), while emergence is a very rare event, summing up to only 1% (Fig S21 to S27). Although domain emergence is such a rare event we find in total more than 500 domains that originate within the arthropods. A detailed look into their function can give interesting insights into arthropod-specific adaptations and innovations. A GO-term enrichment analysis of all rearrangement events results in 'chitin metabolic process' and 'chitin binding' as two of the most functionally relevant terms. This is additional evidence for the importance of exoskeleton development and restructuring during arthropod evolution.

We find the highest total amount of rearrangement events at a single node in *Blattella germanica*, the German cockroach, confirming again the signals we find in the gene family expansion analysis. The impressive capability of the cockroach to adapt to quickly changing environments was apparently the result of a huge amount of evolutionary changes on the genomic level we can trace back with multiple different approaches. The fastest changes over evolutionary time however, we find in the bees (*Apis* and *Bombus*). This signal matches again the results from the gene family analysis.

In general, the high resemblance of signals in this study found with different methods and analyses gives a reliable basis for conclusions about adaptations in the arthropod evolution (Fig. 1).

##### *2.07 Protein Innovation: Silk and venom domain emergences in Chelicerates*

At the root of the order Araneae (spiders; node 70 Fig. 2) we find the first emergence of a domain in the chelicerate phylogeny. The domain refers to a 'Major ampullate spidroin 1' and is also called 'spider silk protein 1' (Pfam-ID: PF16763). Spider silk is a fiber made of specific proteins, spidroins, and is used by spiders for web construction, prey capturing and immobilization, and to build protective egg sacs. Dependent on the task, spiders can change the composition and properties of their silk[68]. The general structure of spider silk consists usually of repetitive segments flanked by a non-repetitive domain at the N- and C-terminus. The major ampullate spidroin 1 is the conserved N-terminal domain that prevents premature aggregation and accelerates and directs the self-assembly[69]. This specific domain is mainly involved in formation of the dragline. Another related domain associated to 'Major ampullate spidroin 1 and 2' (PF11260) emerges at the following node in the Araneae phylogeny together with a domain that is structurally relevant for another type of silk - 'Tubuliform egg casing silk strands structural domain' (PF12042). This domain is found in repeats of Tubuliforms that are used to protect egg cases[70]. In addition to these very specialized domains for spider silks we find venom related domains that emerge in several spider lineages. In *Lactrodectus hesperus*, the western black widow spider, a 'Toxin with inhibitor cystine knot ICK or Knottin scaffold' (PF10530) emerges that is an important part of the neurotoxin produced by this species[71]. New venom related domains can be found in other chelicerates as well. In total three new domains show up in the Arizona bark scorpion, *Centruroides sculpturatus*, that are part of its venom, more precisely 'Scorpion short toxin, BmKK2' (PF00451), 'CagA exotoxin' (PF03507) and 'Ergtoxin family' (PF08086). These examples indicate that we can identify genomic signatures, with corresponding phenotypic and behavioral observations.

#### 3. Supplementary Tables.

##### *3.01. List of Large Supplementary Tables as worksheets in Microsoft Excel file "Large Supplementary Tables".*

| | |
|---|---|
| Table S1. | Species Abbr. & NCBI Accessions. |
| Table S2. | DNA Sequence Stats. |
| Table S3. | Assemblies. |
| Table S4. | RNAseq Stats. |
| Table S5. | Automated Annotation. |
| Table S6. | OrthoDB Geneset Sources. |
| Table S7. | Busco and Dogma Completeness Scores for 76 Arthropod Species. |
| Table S8. | OGS Manual Annotation. |
| Table S9. | GC Content. |
| Table S10. | Kmer Analysis. |
| Table S11. | All Gene Family Data. |
| Table S12. | Species Gene Family Data. |
| Table S13. | Multi Species Orders. |
| Table S14. | Fossil Calibrations. |
| Table S15. | Top30 Changing Gene Fams. |
| Table S16. | Ancestral Gene Counts. |
| Table S17. | Major Transistions |
| Table S18. | Novel LICA Fams. |
| Table S19 | Novel Holometabola Fams. |
| Table S20. | Spider Venom Silk Fams. |
| Table S21. | Enriched GO-ZNEVA-25. |
| Table S22. | Dipteran Overview. |
| Table S23. | Dipteran Root. |
| Table S24. | DL11 Lucilia-Musca. |
| Table S25. | DL17 Schizophora. |
| Table S26. | DL13 Calyptrates. |

##### *3.02. Calculated rates of rearrangement events.*

| set | fusion | fission | term loss | term gain | single domain loss | single domain gain |
|---|---|---|---|---|---|---|
| Arthropods (i5K) | 49.01% | 13.67% | 13.53% | 0.47% | 22.37% | 0.95% |

##### *3.03. Calculated exact numbers of rearrangement events.*

| Event type | Arthropod set |
|---|---|
| Exact solutions | 39018 |
| Non-ambiguous solutions | 2587 |
| Ambiguous solutions | 1472 |
| Complex solutions | 47135 |
| Fusions | 20392 |
| Fissions | 5687 |
| Terminal loss | 5628 |
| Terminal gain | 195 |
| Single domain loss | 9309 |
| Single domain gain | 394 |
| Exact and non-ambigous solutions | 41605 |

#### 4. Supplementary Figures.

**Figure S1. Counts of 195 i5K Nominated Species by Order.**

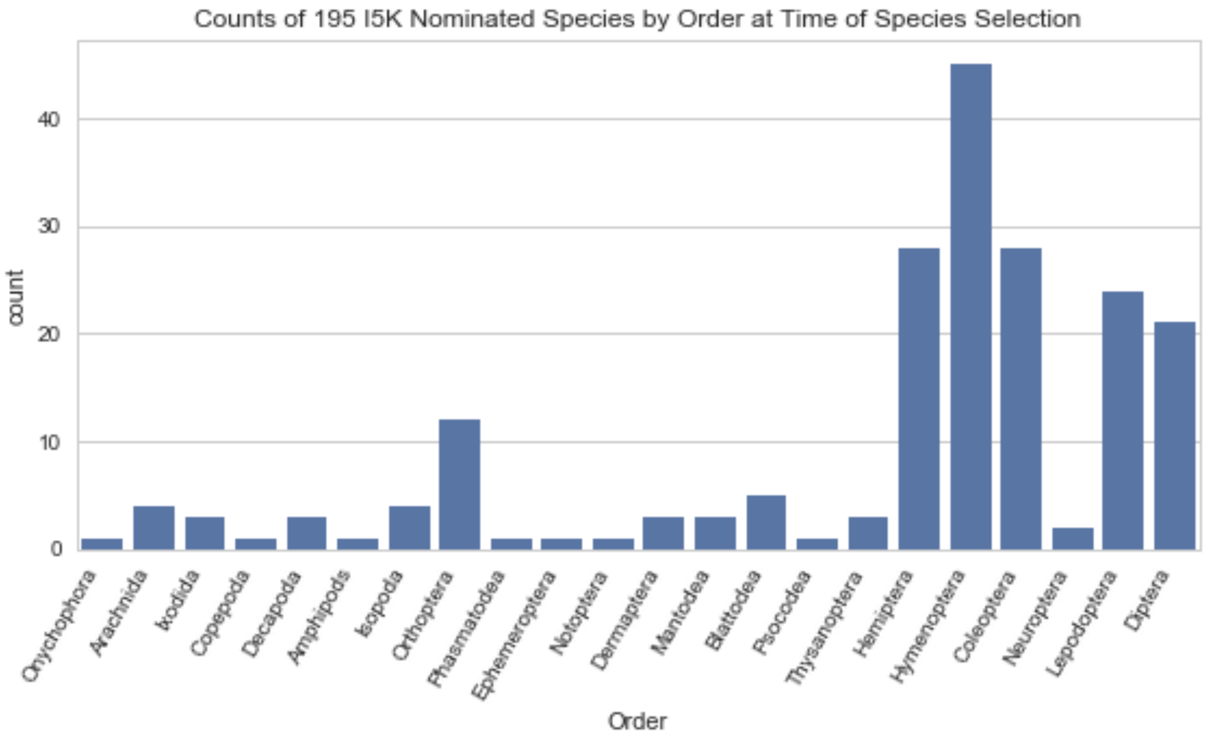

**Figure S2. Assembly and Maker 2.0 CDS GC content for Redundans assemblies.**

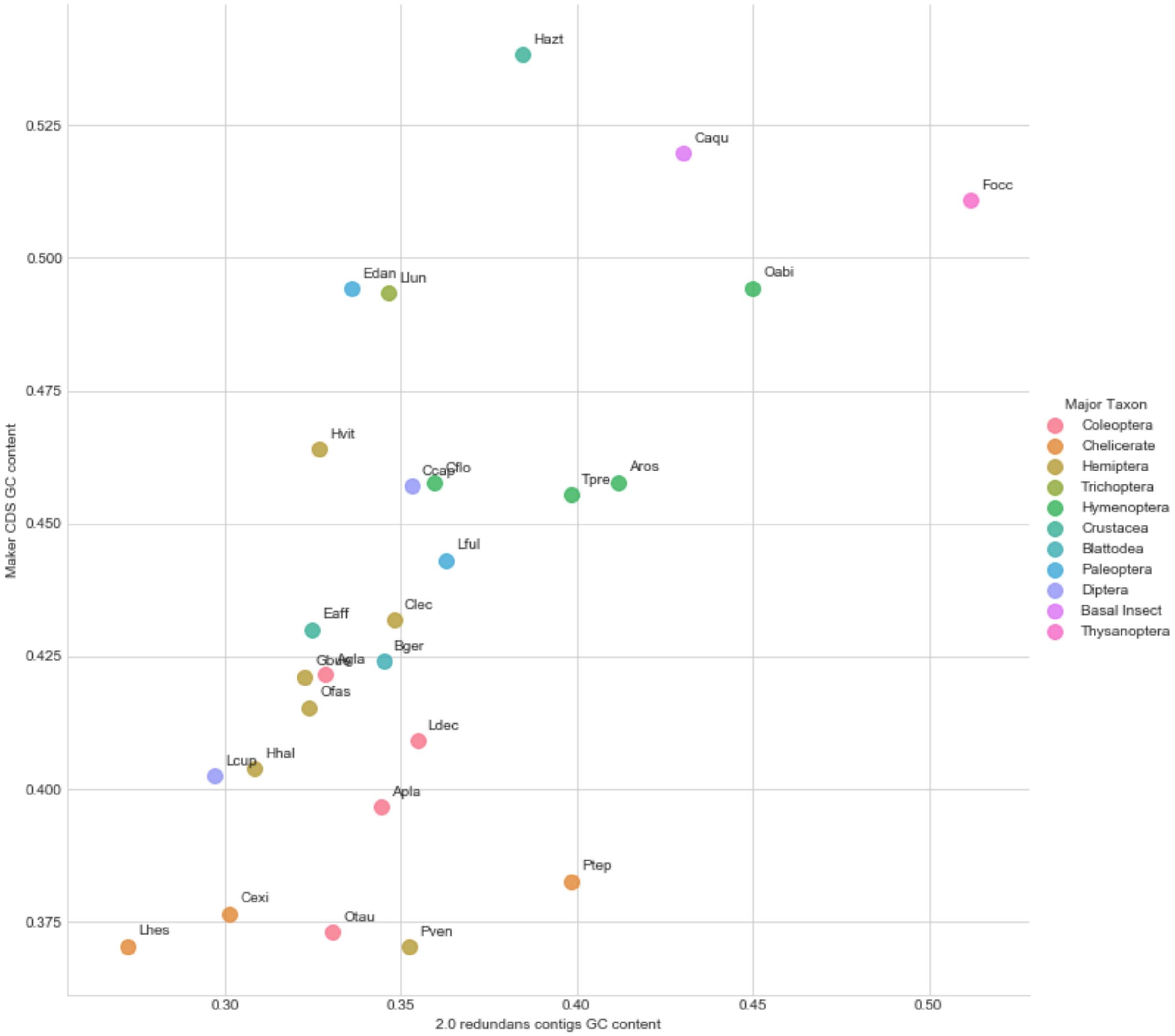

### Figure S3. Kmer analysis of i5K pilot species 500bp read libraries at 17, 21 and 31 bp.

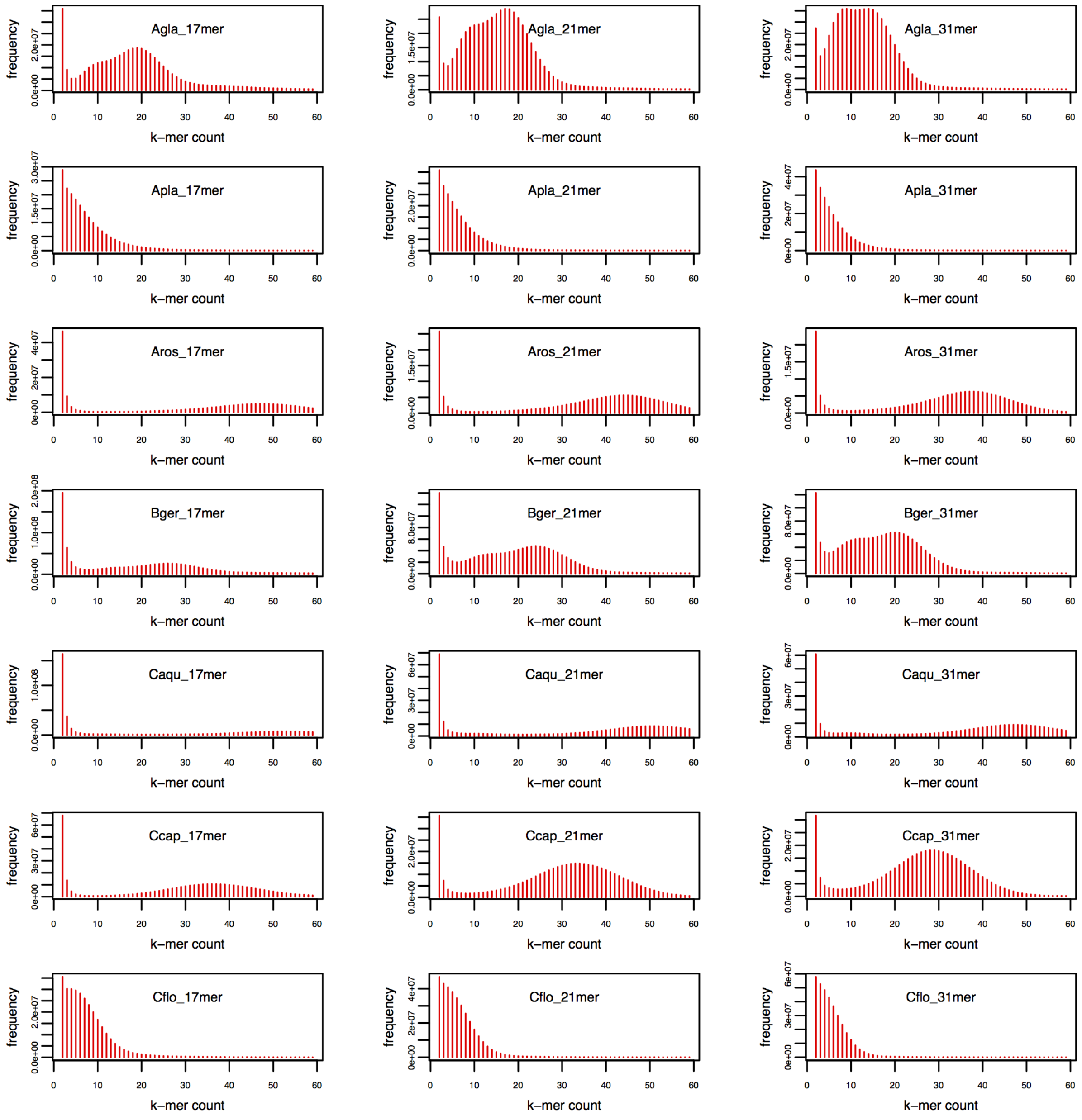

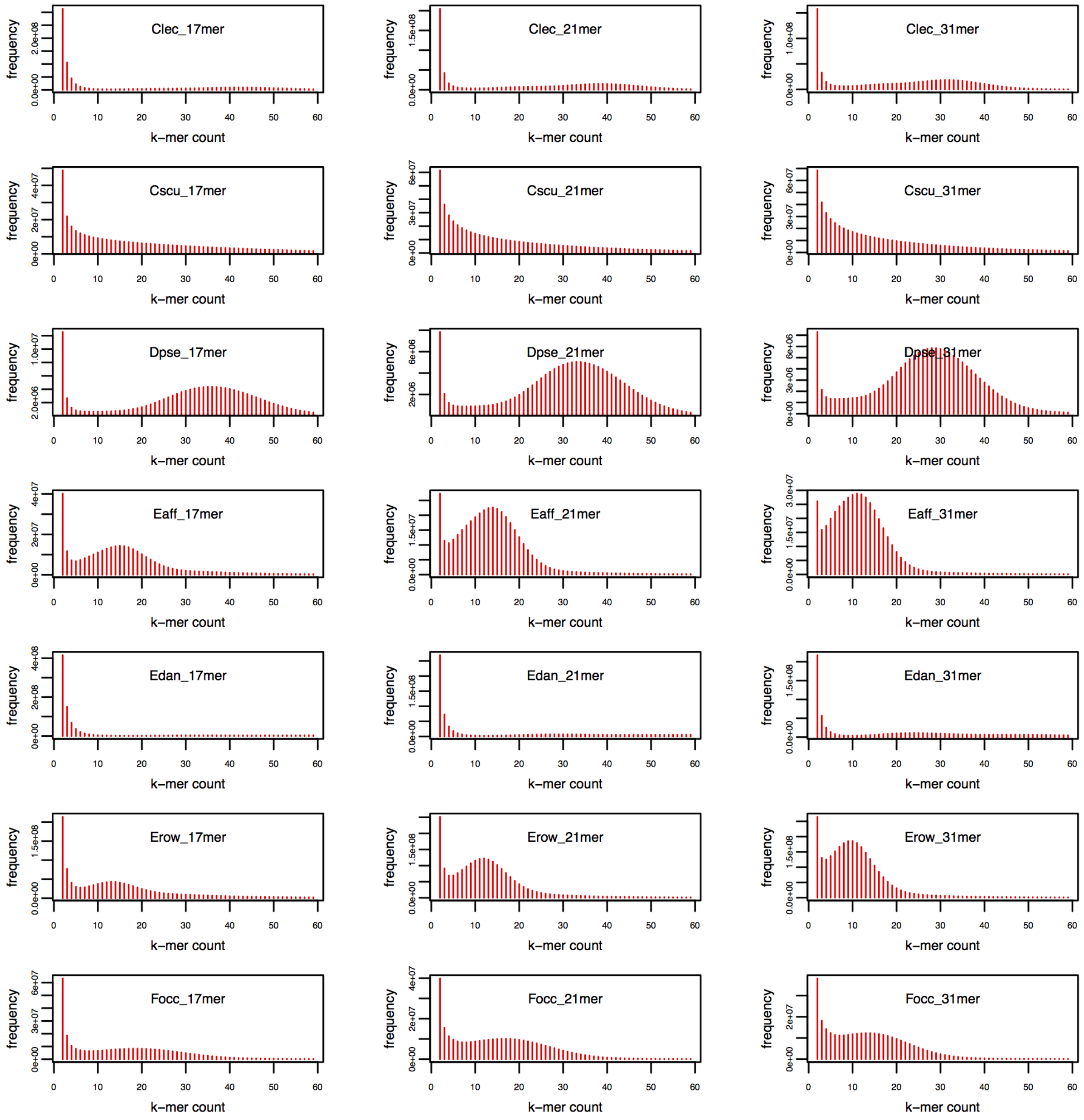
Clec_17mer
Clec_21mer
Clec_31mer
Cscu_17mer
Cscu_21mer
Cscu_31mer
Dpse_17mer
Dpse_21mer
Dpse_31mer
Eaff_17mer
Eaff_21mer
Eaff_31mer
Edan_17mer
Edan_21mer
Edan_31mer
Erow_17mer
Erow_21mer
Erow_31mer
Focc_17mer
Focc_21mer
Focc_31mer
frequency
k–mer count

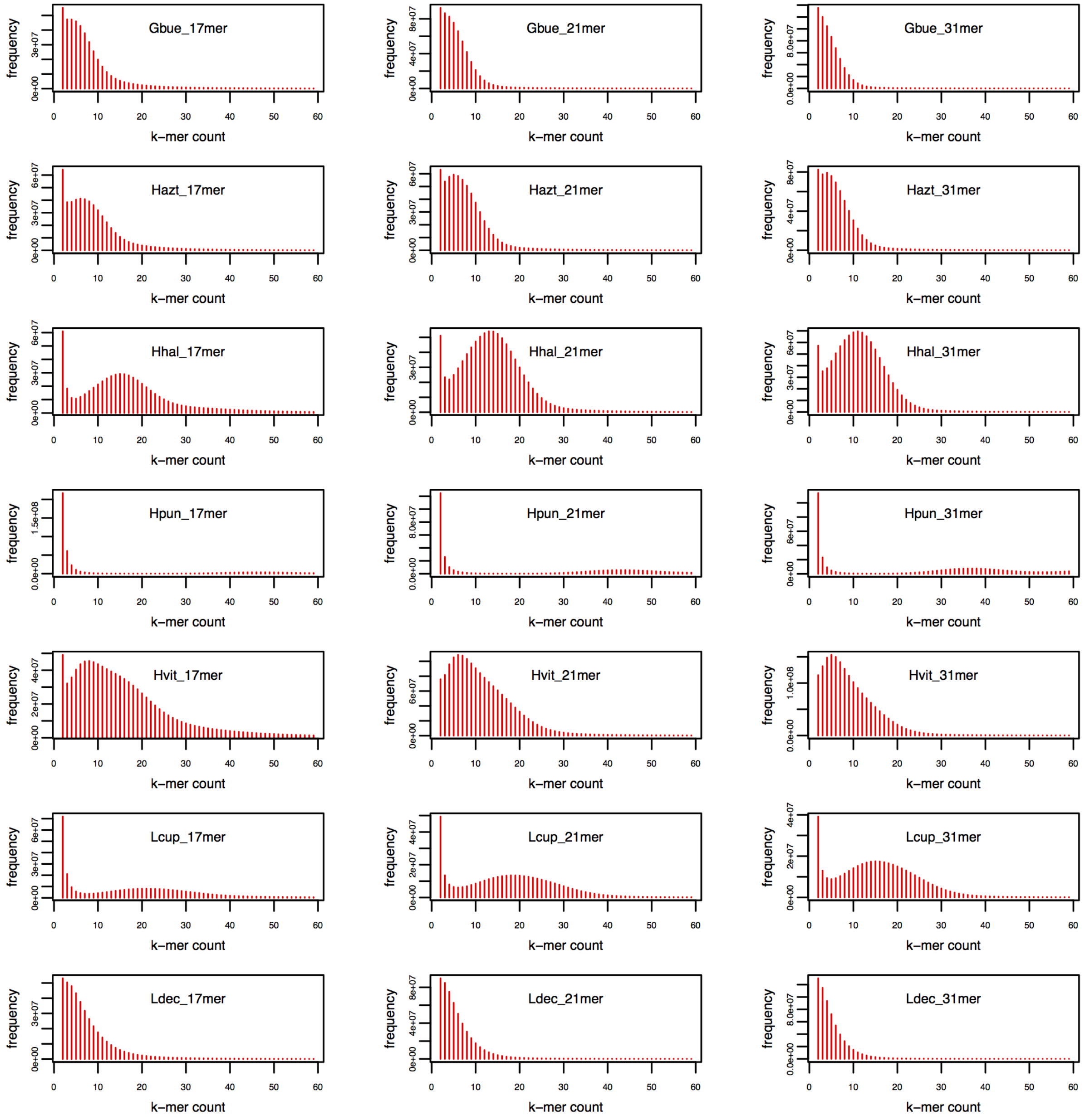
Gbue_17mer
Gbue_21mer
Gbue_31mer
Hazt_17mer
Hazt_21mer
Hazt_31mer
Hhal_17mer
Hhal_21mer
Hhal_31mer
Hpun_17mer
Hpun_21mer
Hpun_31mer
Hvit_17mer
Hvit_21mer
Hvit_31mer
Lcup_17mer
Lcup_21mer
Lcup_31mer
Ldec_17mer
Ldec_21mer
Ldec_31mer
frequency
k–mer count
0 10 20 30 40 50 60

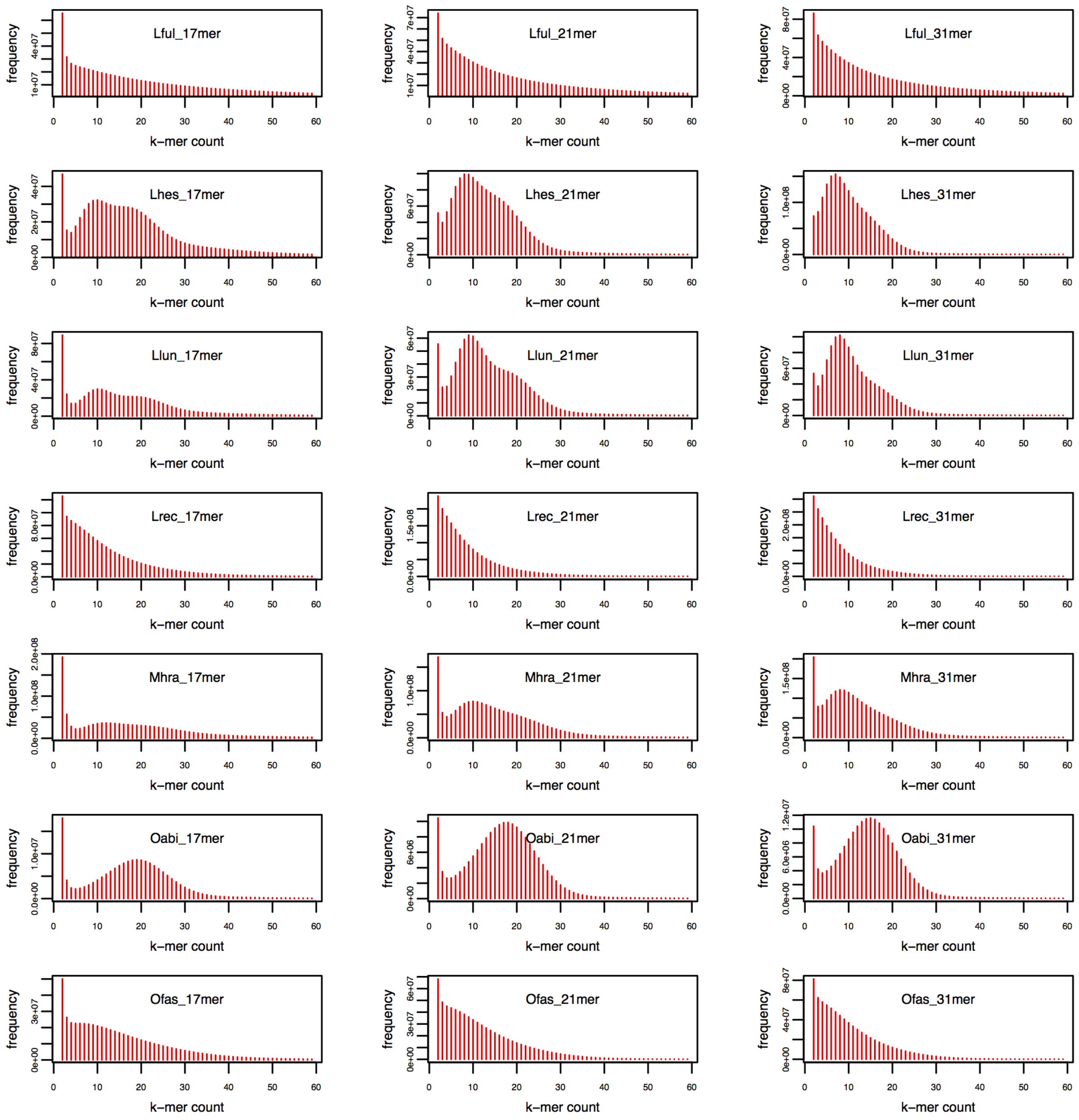
Lful_17mer
Lful_21mer
Lful_31mer
Lhes_17mer
Lhes_21mer
Lhes_31mer
Llun_17mer
Llun_21mer
Llun_31mer
Lrec_17mer
Lrec_21mer
Lrec_31mer
Mhra_17mer
Mhra_21mer
Mhra_31mer
Oabi_17mer
Oabi_21mer
Oabi_31mer
Ofas_17mer
Ofas_21mer
Ofas_31mer
frequency
k–mer count

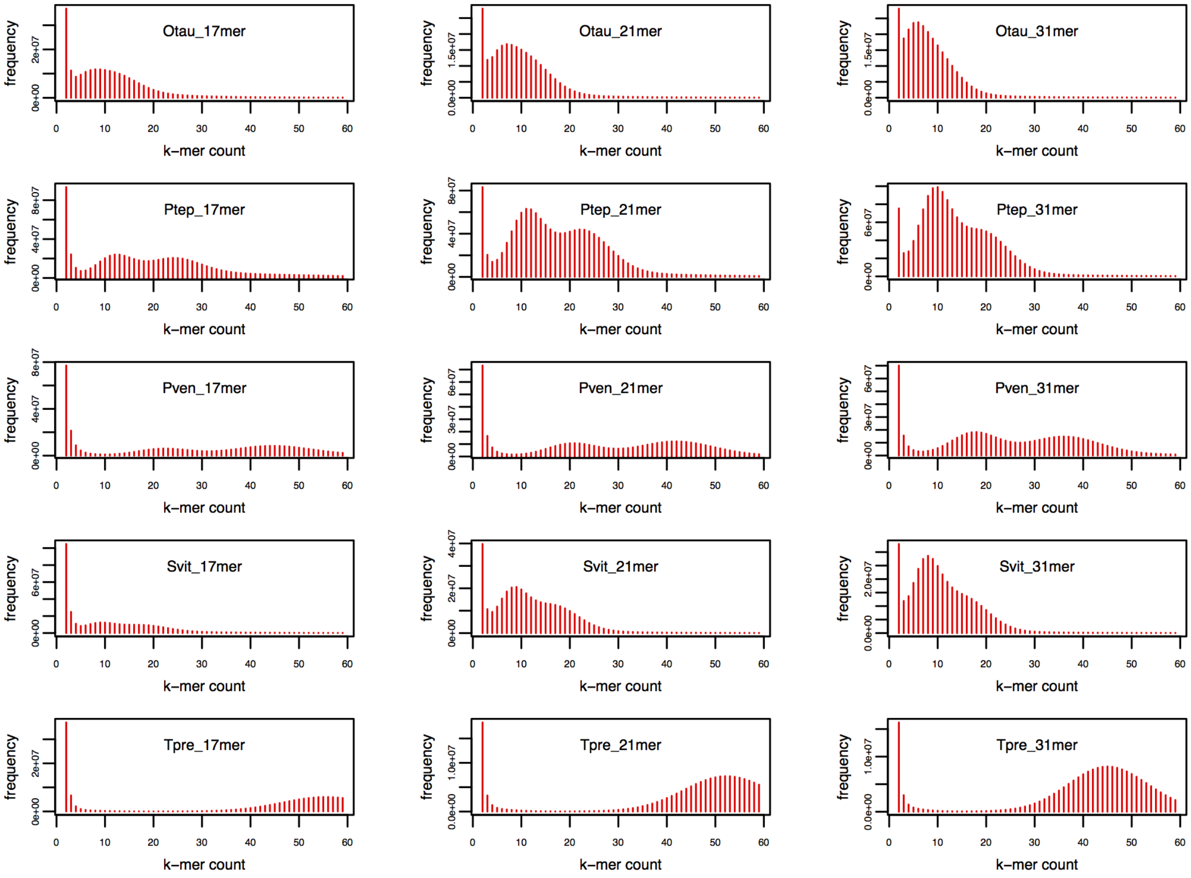
Otau_17mer
Otau_21mer
Otau_31mer
Ptep_17mer
Ptep_21mer
Ptep_31mer
Pven_17mer
Pven_21mer
Pven_31mer
Svit_17mer
Svit_21mer
Svit_31mer
Tpre_17mer
Tpre_21mer
Tpre_31mer
frequency
k−mer count

**Figure S4. OrthoDB orthology delineation for the i5K pilot species.**

(**A**) The area-proportional pie charts show the proportions of the total of 105 clustered and mapped arthropod species belonging to the sampled subclades for each of five major nodes of the arthropod phylogeny. (**B**) The bars show Metazoa-level orthologs for the 76 selected arthropods and three outgroup species partitioned according to their presence and copy-number, sorted from the largest total gene counts to the smallest. The 28 i5K species abbreviations are indicated in bold font with species images positioned above each bar.

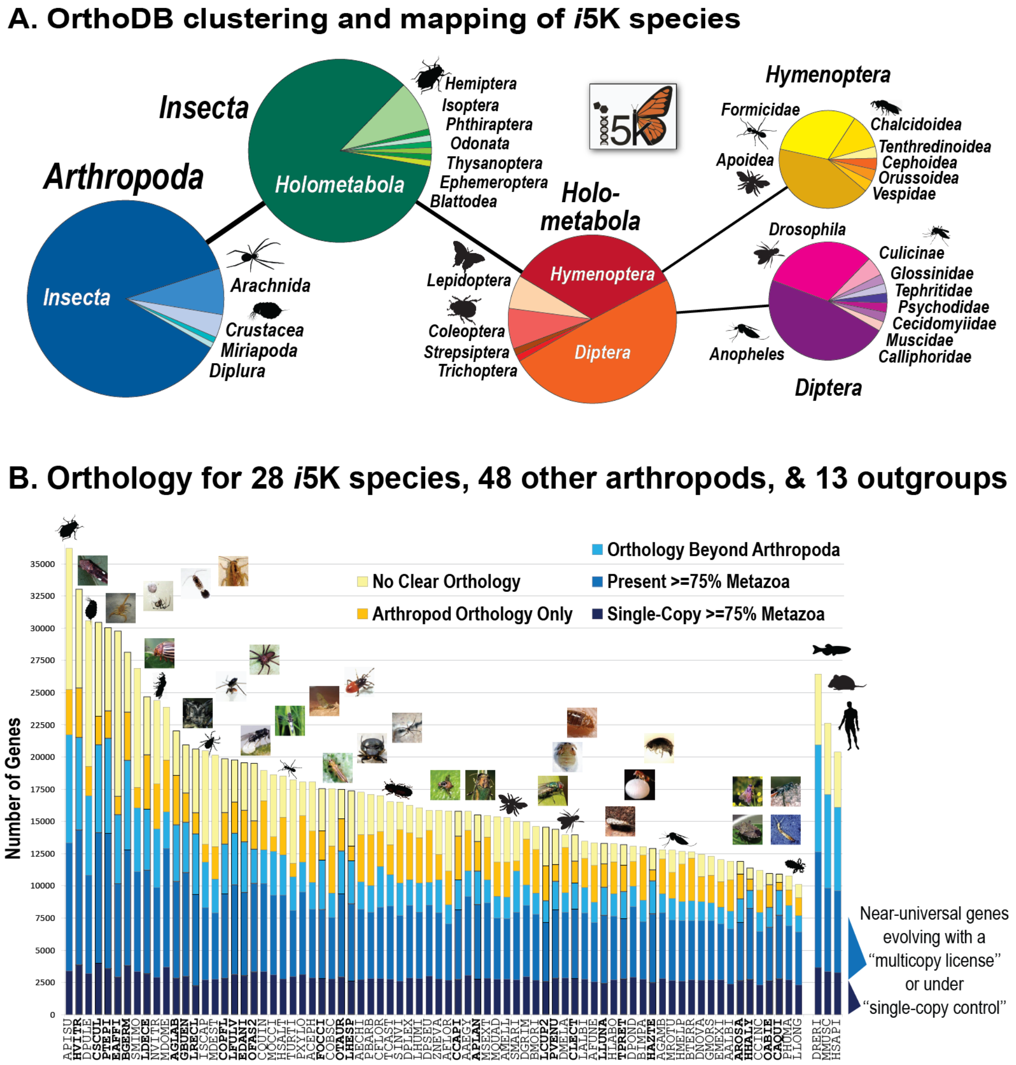

**Figure S5: Estimating gene counts at ancestral nodes.**

Gene content at ancestral nodes in the phylogeny is estimated by summing up the number of inferred proteins at any given node. This number, however, does not account for unobserved extinctions. This leads to a negative correlation between the amount time that has passed and the inferred number of genes at a given internal node (**A**). To correct for this, we used that correlation to estimate the number of genes expected in an arthropod at any time if no gene changes have occurred (i.e. the number observed at Split time = 0) and accounted for the stochasticity by translating the observed residuals around that point (**B**).

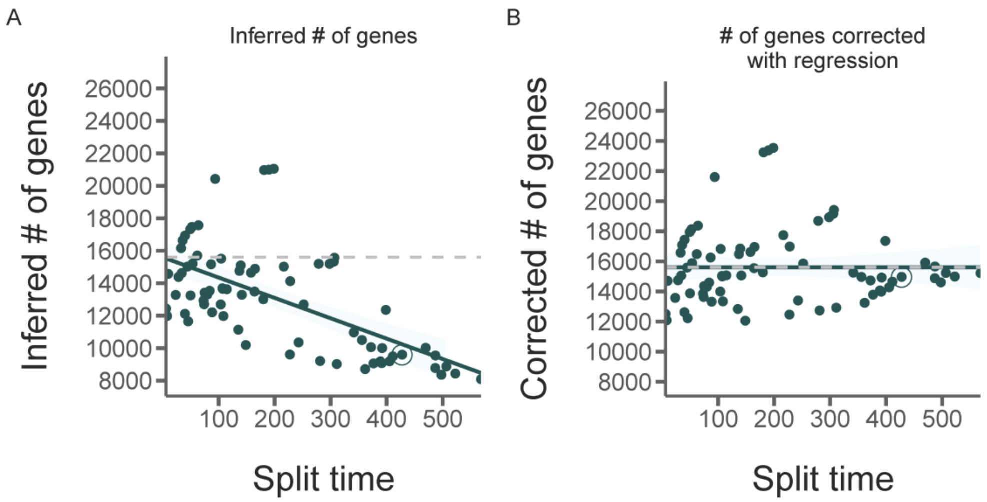

**Figure S6. Protein domain reconstruction and rearrangement event inference.**

Given a phylogenetic tree and annotated protein domains for all studied species (**A**) it becomes possible to reconstruct the ancestral domain content for every inner node, by comparing the content of the child nodes and infer the ancestral state based on a parsimony principle (red box). (**B**) In the reconstructed tree different rearrangement events can be inferred by comparing every node to its parental node (green boxes). (**C**) Six different event types are considered and inferred based on different parsimony principles (see methods). All arrangements are inferred by a Fitch Parsimony (orange), while all single domain states are inferred by a Dollo Parsimony (blue).

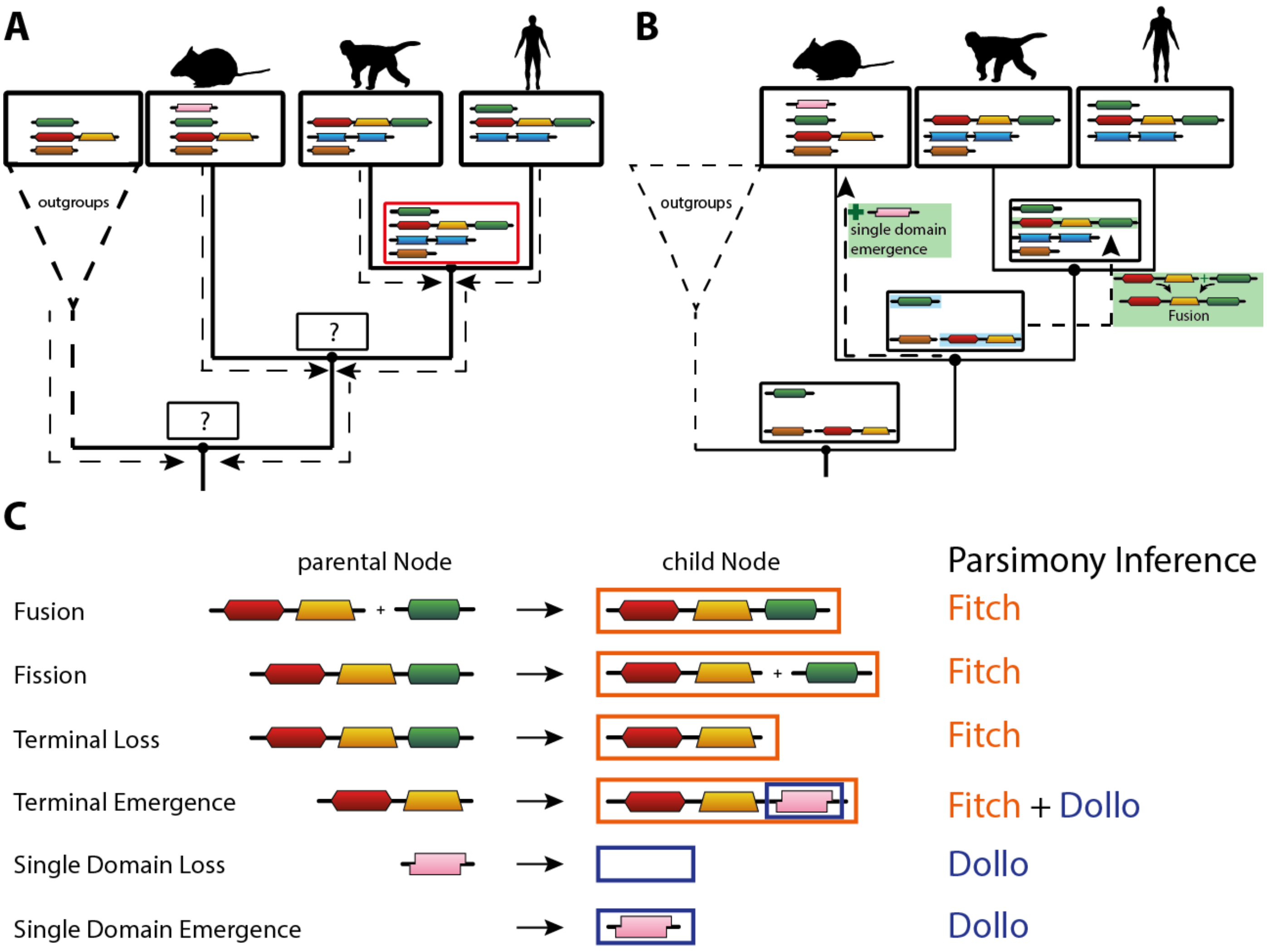

**Figure S7. Presence of DNA methylation machinery across the arthropods.**

Evidence of DNA methylation and DNA methylation machinery across 76 arthropod taxa. Top four panels: E-values indicating the presence of DNMT1, DNMT3, MBD2, and UHRF1, respectively. For DNMT genes, each box represents the best hit e-value for six annotated DNMT orthologs (3 in insects, 1 in basal invertebrates, 2 from vertebrates). The red lines indicate an e-value of 1e-10. Strongly negative values suggest a match and indicate the presence of the focal gene in the target species. Estimated gene copy number is provided above each panel. Bottom panel: Mean coding sequence CpG o/e for each species. CpG o/e is negatively correlated with DNA methylation. Therefore, low values of CpG o/e suggest the presence of DNA methylation in the genome.

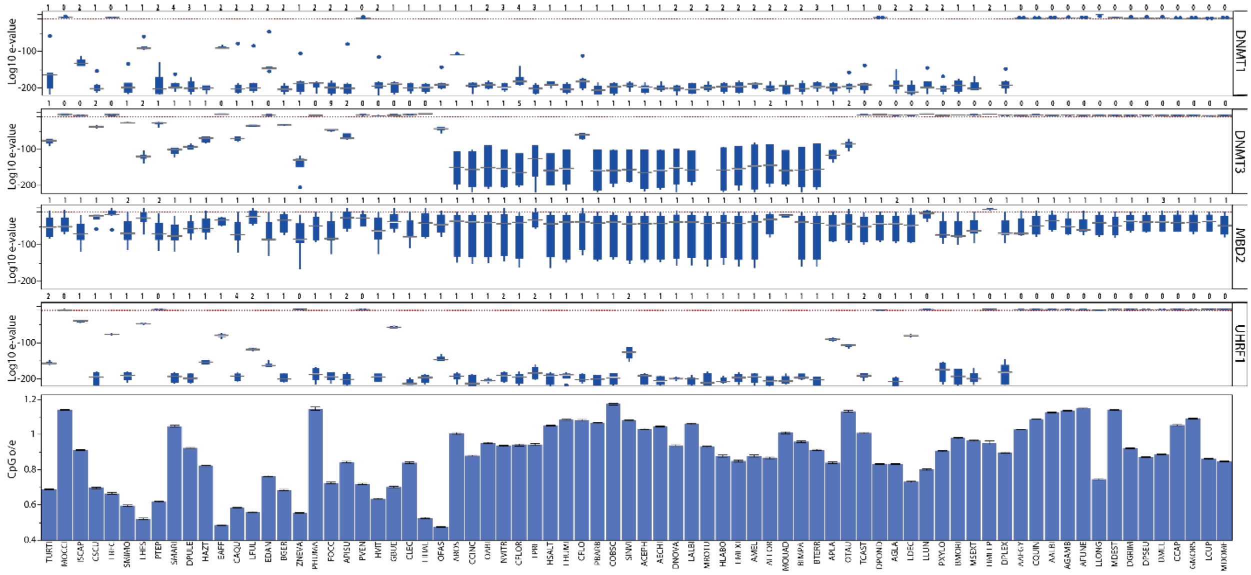

**Figure S8. Patterns of DNA methylation CpG 0/e signals of 72 arthropod species.**

The first two panels next to each taxon provide the mean level and distribution of CpG o/e in exons (green dashed line), introns (black dotted line), genomic windows (non-genic background, light blue dashed line), and gene frames (exons + introns, red solid line). The third panel illustrates CpG o/e levels across genic positions for 'high' and 'low' CpG o/e genes (*i.e.*, genes falling above or below the mean CpG o/e of all genes for a given species, respectively).

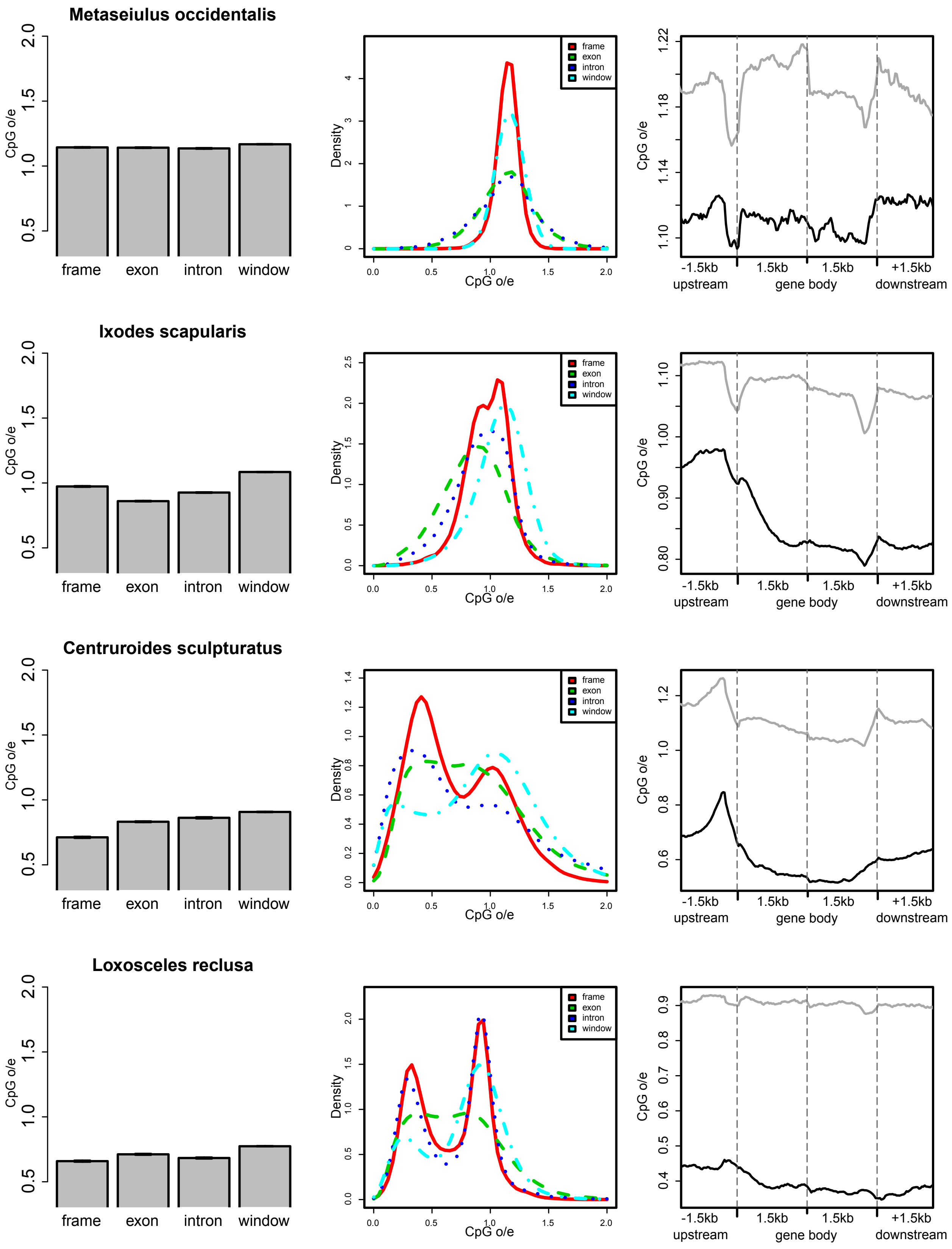
Metaseiulus occidentalis
Ixodes scapularis
Centruroides sculpturatus
Loxosceles reclusa
CpG o/e
frame
exon
intron
window
Density
-1.5kb
upstream
1.5kb
gene body
+1.5kb
downstream

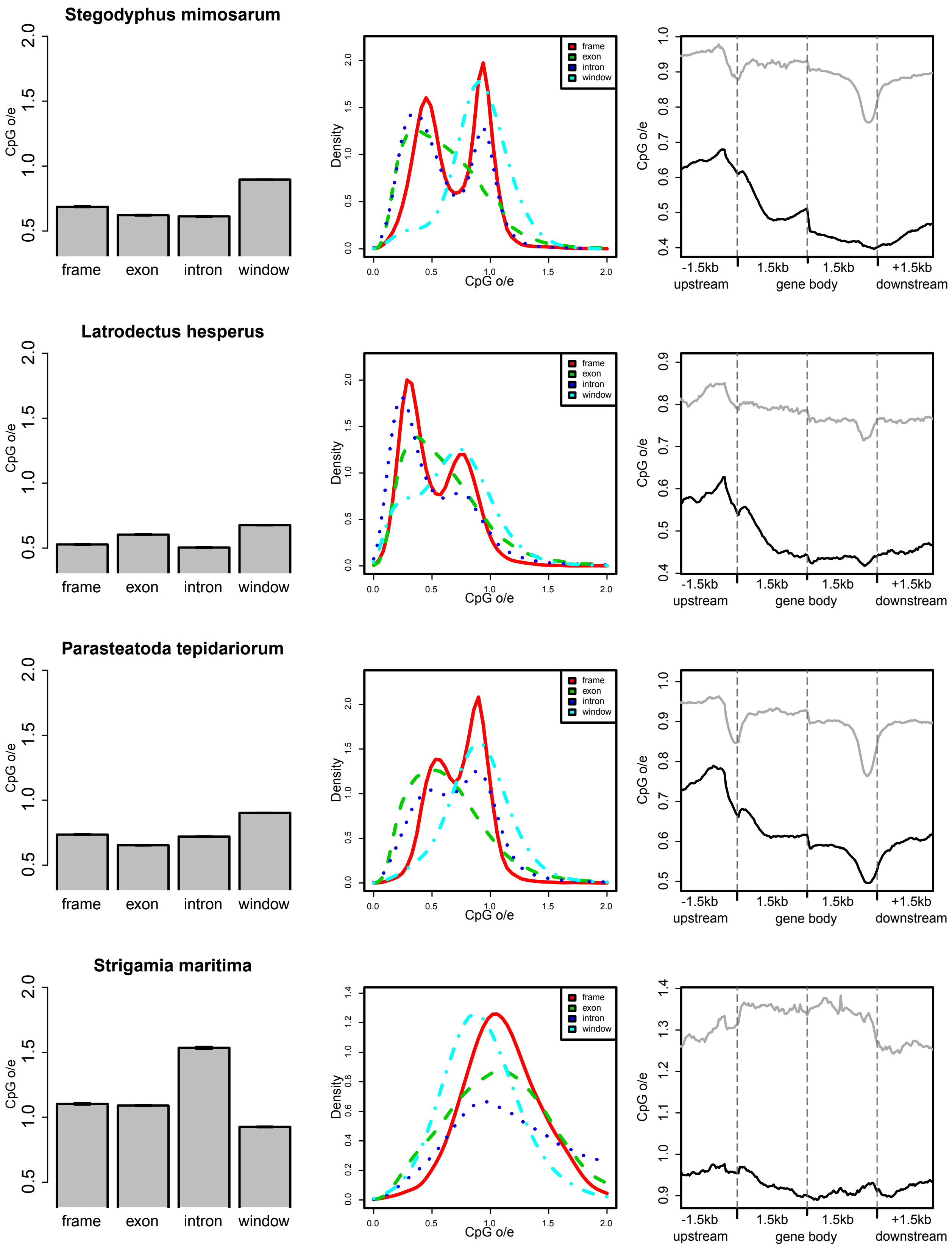
Stegodyphus mimosarum
Latrodectus hesperus
Parasteatoda tepidariorum
Strigamia maritima
CpG o/e
frame
exon
intron
window
Density
-1.5kb
upstream
1.5kb
gene body
+1.5kb
downstream

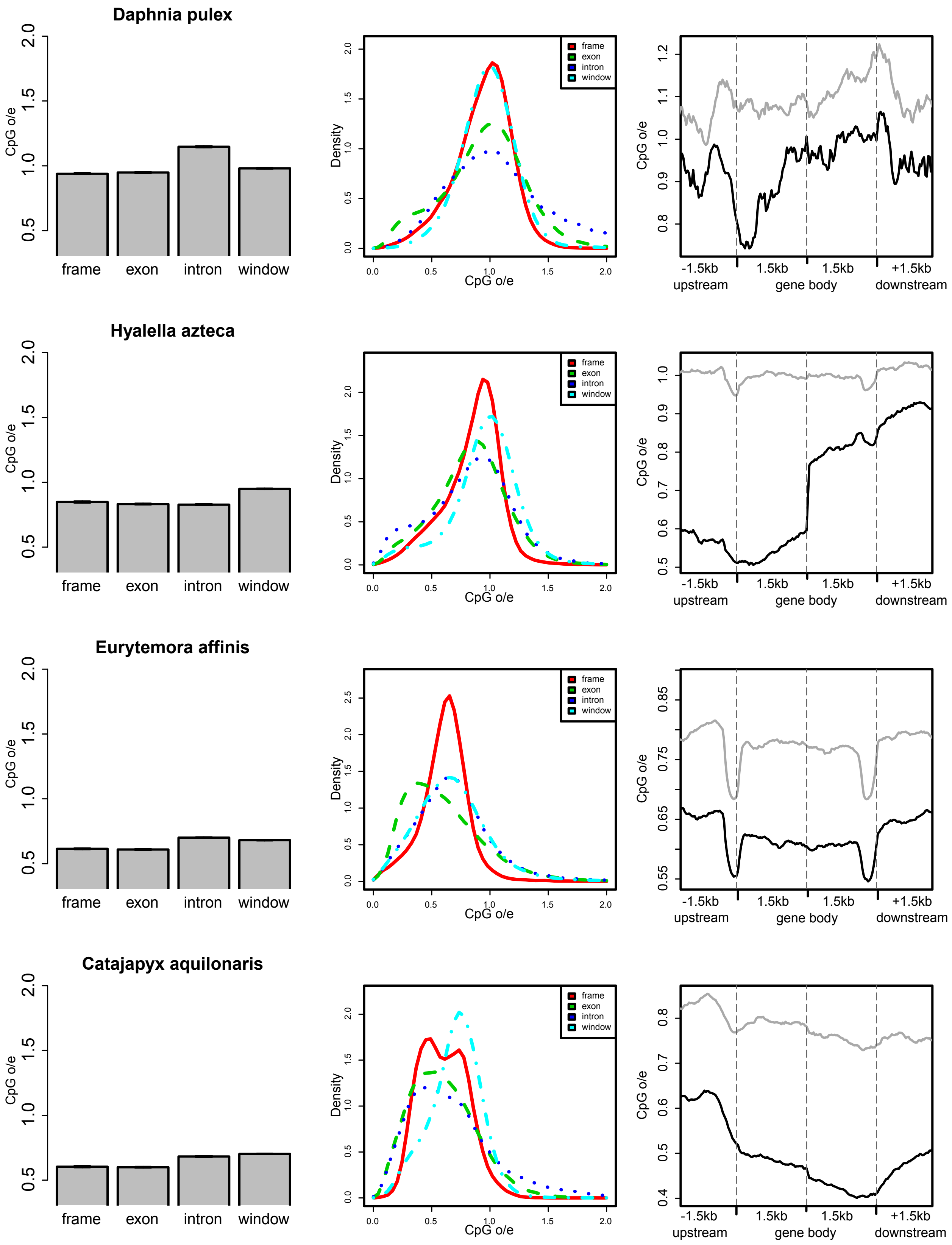
Daphnia pulex
Hyalella azteca
Eurytemora affinis
Catajapyx aquilonaris
CpG o/e
frame
exon
intron
window
Density
-1.5kb
upstream
1.5kb
gene body
+1.5kb
downstream

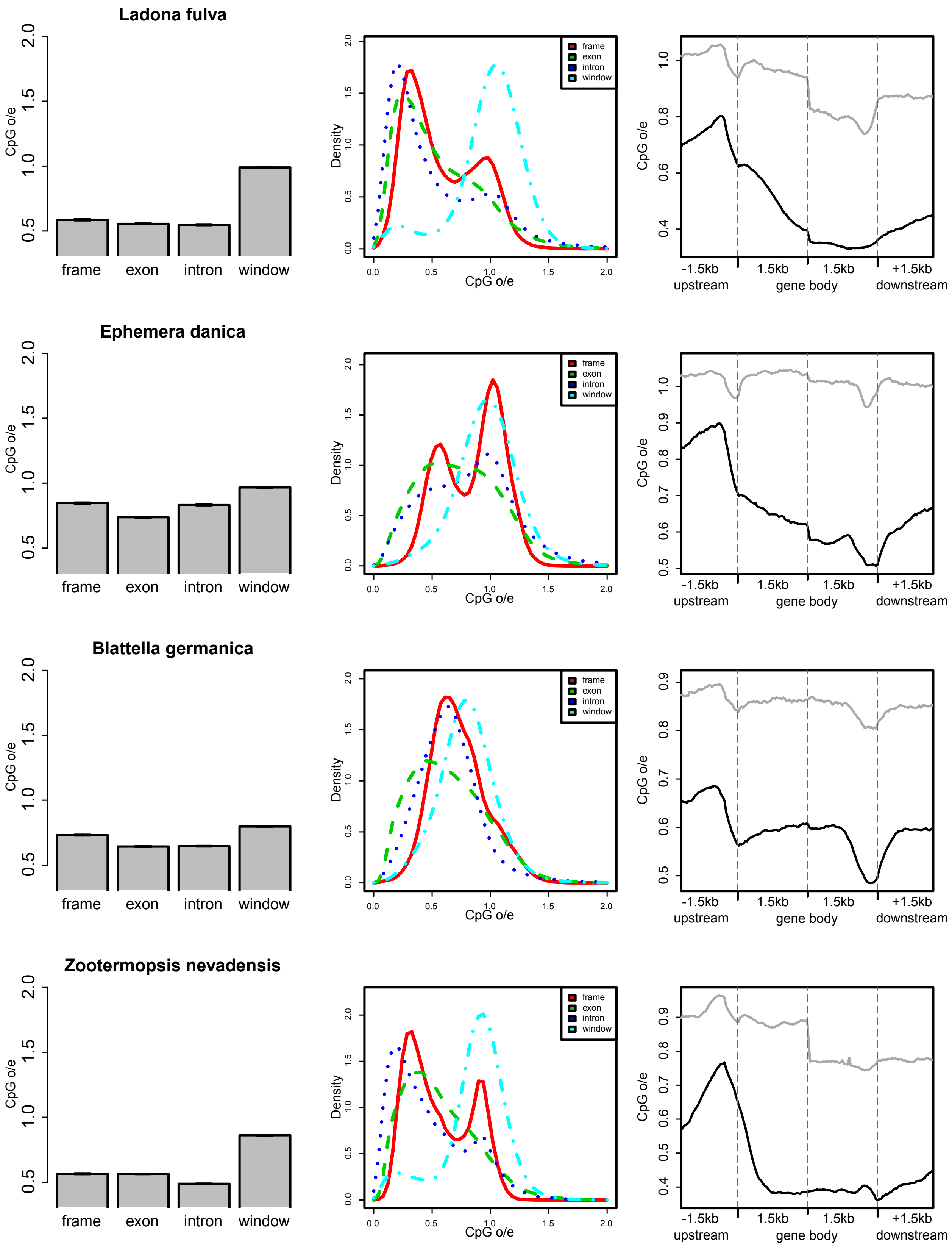
Ladona fulva
Ephemera danica
Blattella germanica
Zootermopsis nevadensis
CpG o/e
frame
exon
intron
window
Density
CpG o/e
-1.5kb
upstream
1.5kb
gene body
1.5kb
+1.5kb
downstream

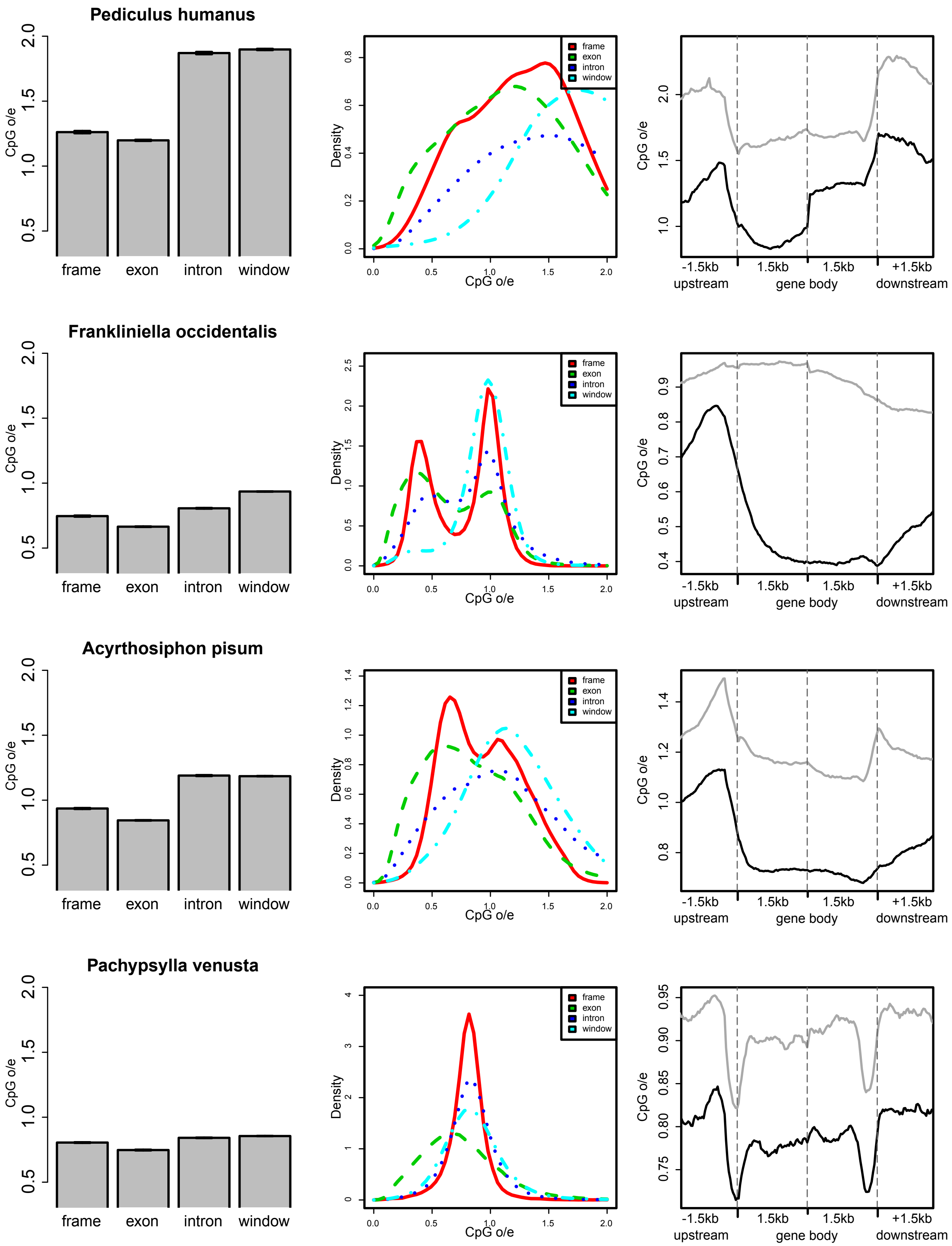
Pediculus humanus
Frankliniella occidentalis
Acyrthosiphon pisum
Pachypsylla venusta
CpG o/e
frame
exon
intron
window
Density
upstream
gene body
downstream
-1.5kb
1.5kb
+1.5kb

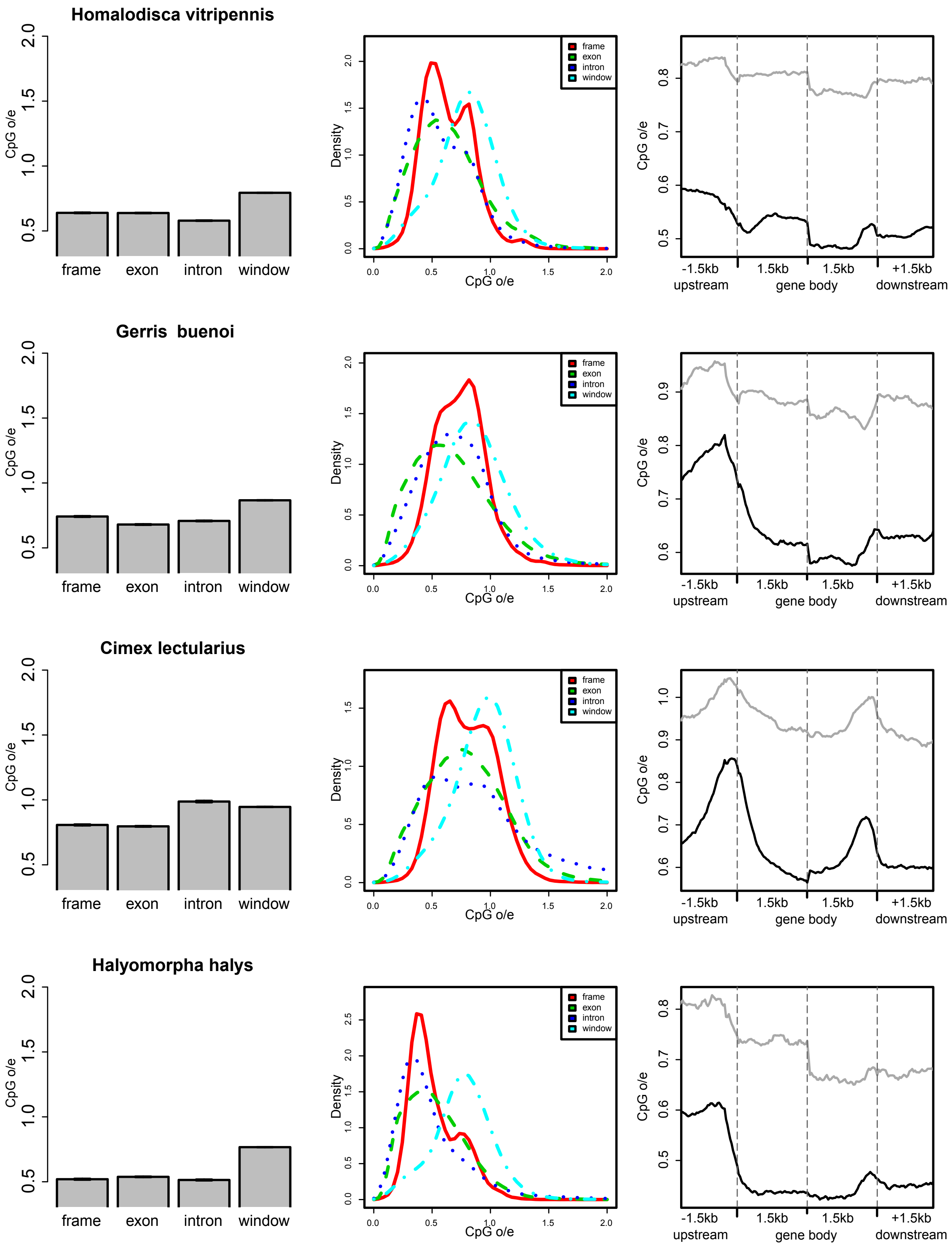
Homalodisca vitripennis
Gerris buenoi
Cimex lectularius
Halyomorpha halys
CpG o/e
frame
exon
intron
window
Density
CpG o/e
-1.5kb
upstream
1.5kb
gene body
1.5kb
+1.5kb
downstream

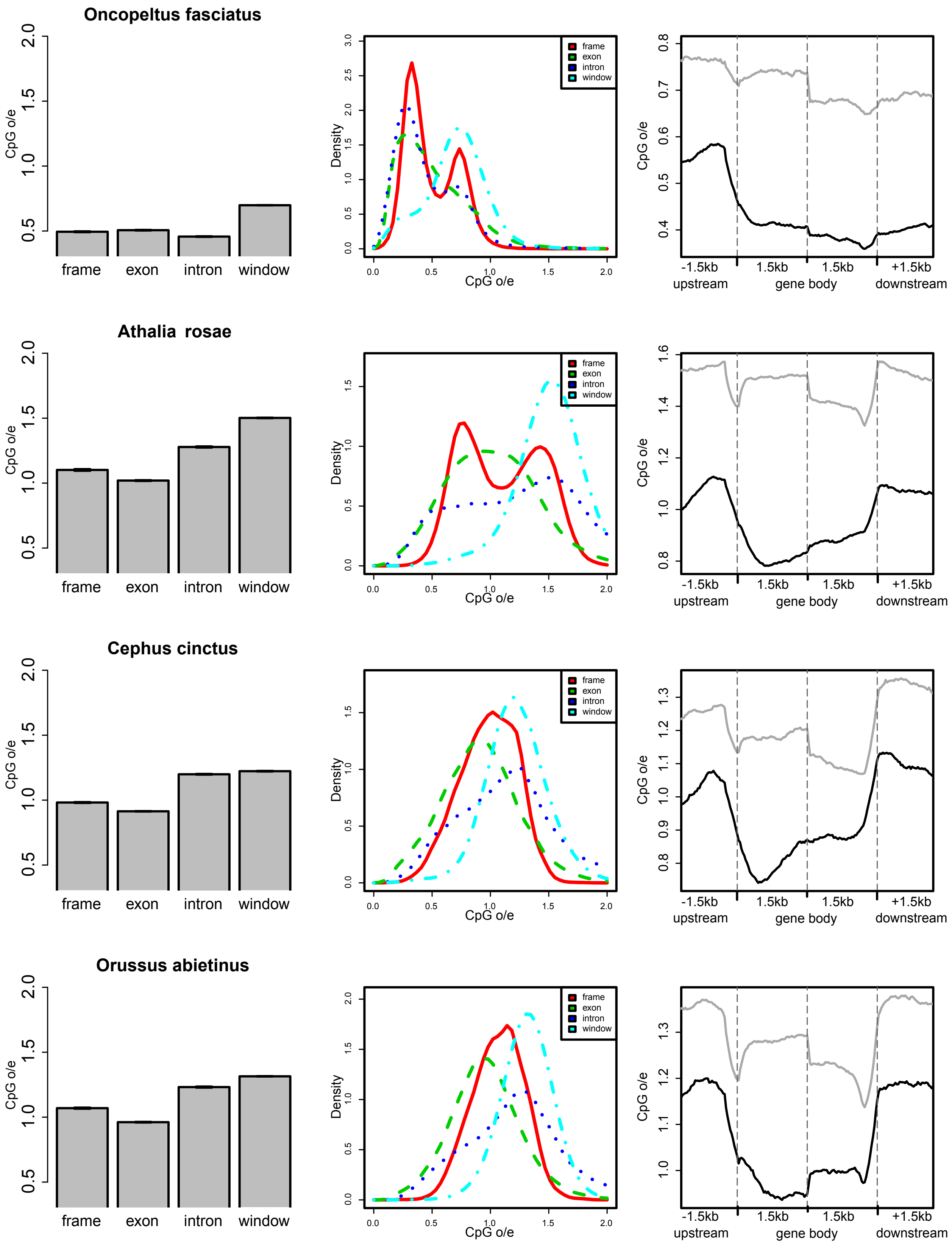

Oncopeltus fasciatus
Athalia rosae
Cephus cinctus
Orussus abietinus
CpG o/e
frame
exon
intron
window
Density
-1.5kb
upstream
1.5kb
gene body
+1.5kb
downstream

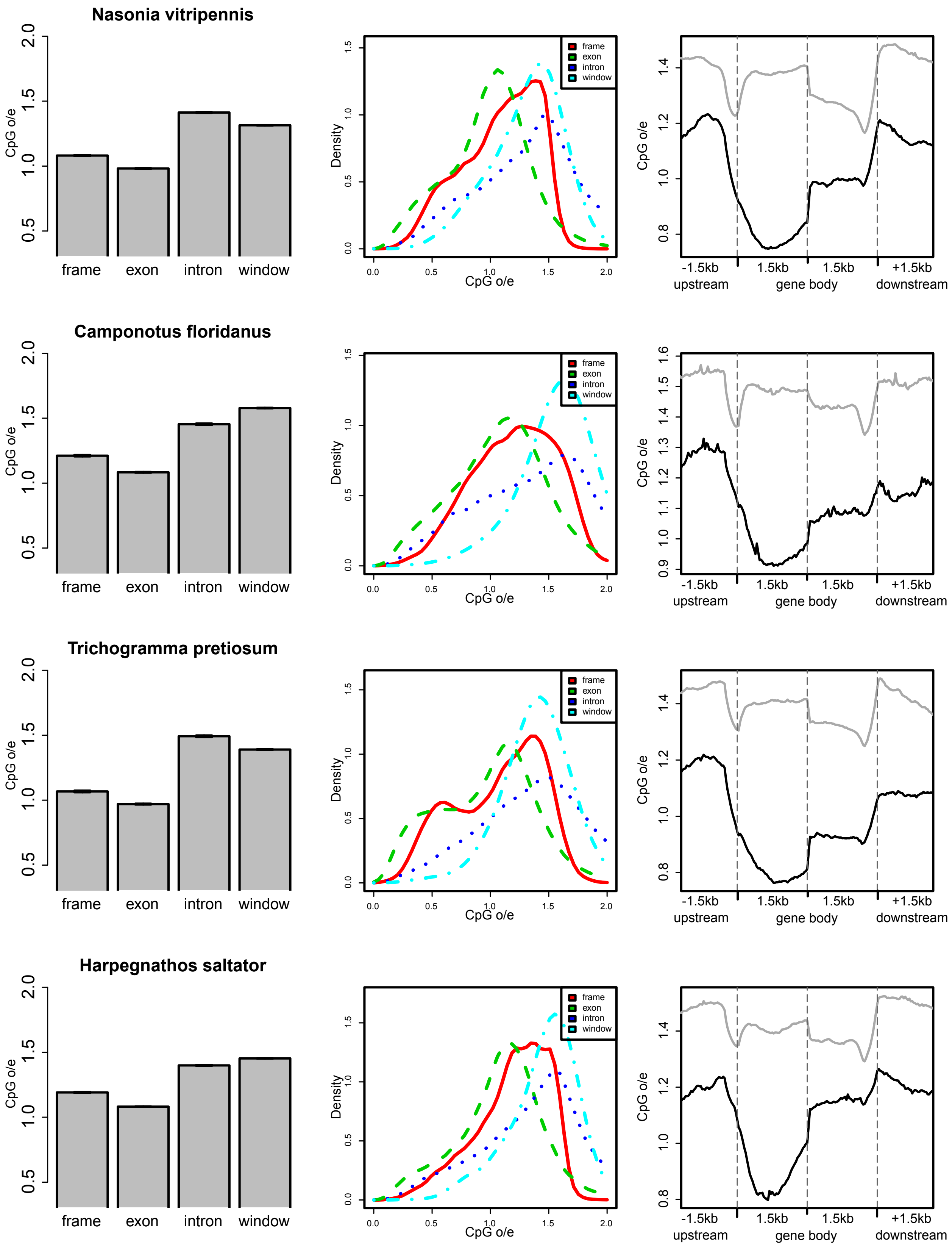
Nasonia vitripennis
Camponotus floridanus
Trichogramma pretiosum
Harpegnathos saltator
CpG o/e
frame
exon
intron
window
Density
-1.5kb
upstream
1.5kb
gene body
+1.5kb
downstream

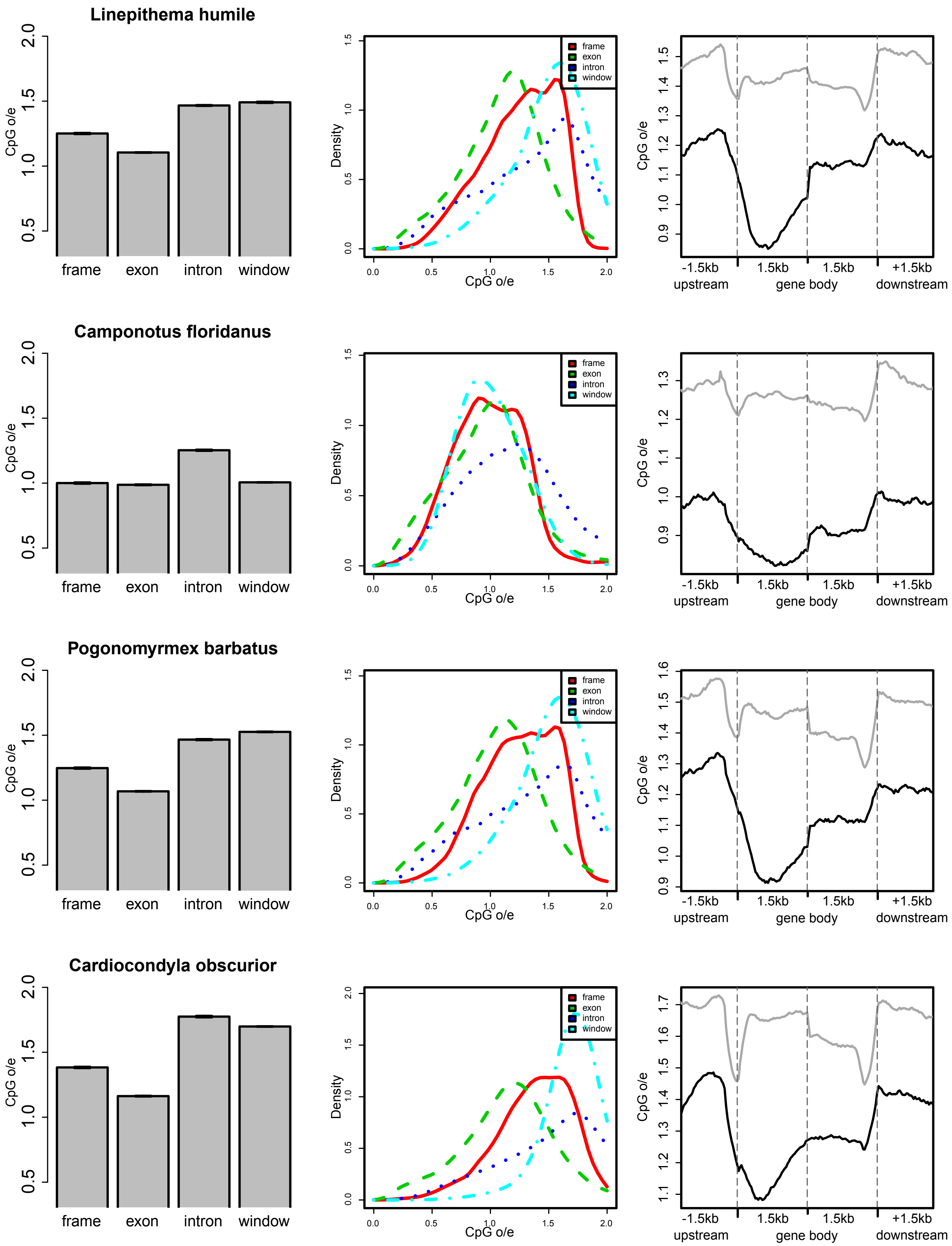
Linepithema humile
Camponotus floridanus
Pogonomyrmex barbatus
Cardiocondyla obscurior
CpG o/e
frame
exon
intron
window
Density
-1.5kb
upstream
1.5kb
gene body
+1.5kb
downstream

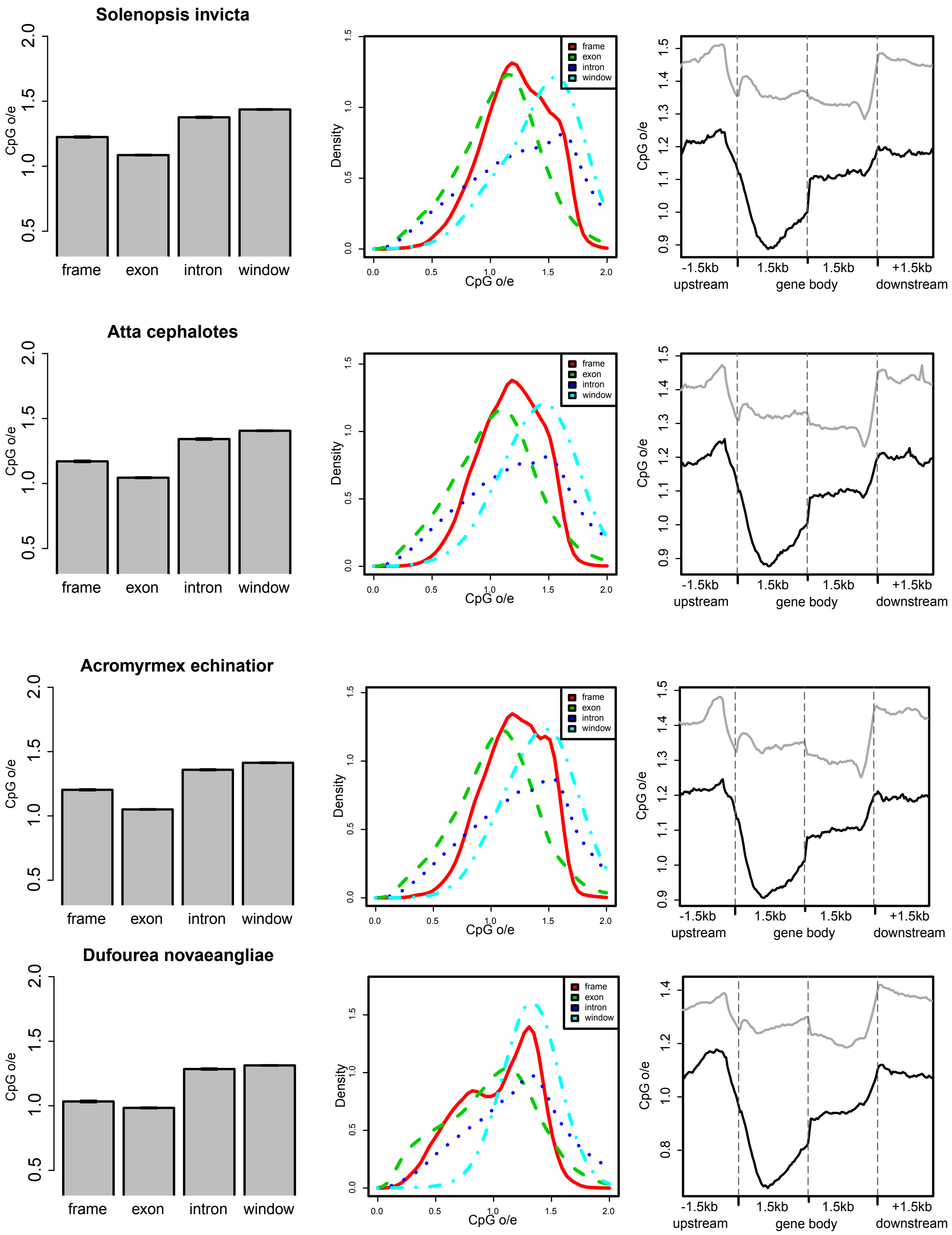
Solenopsis invicta
Atta cephalotes
Acromyrmex echinatior
Dufourea novaeangliae
CpG o/e
frame
exon
intron
window
Density
upstream
gene body
downstream
-1.5kb
1.5kb
+1.5kb

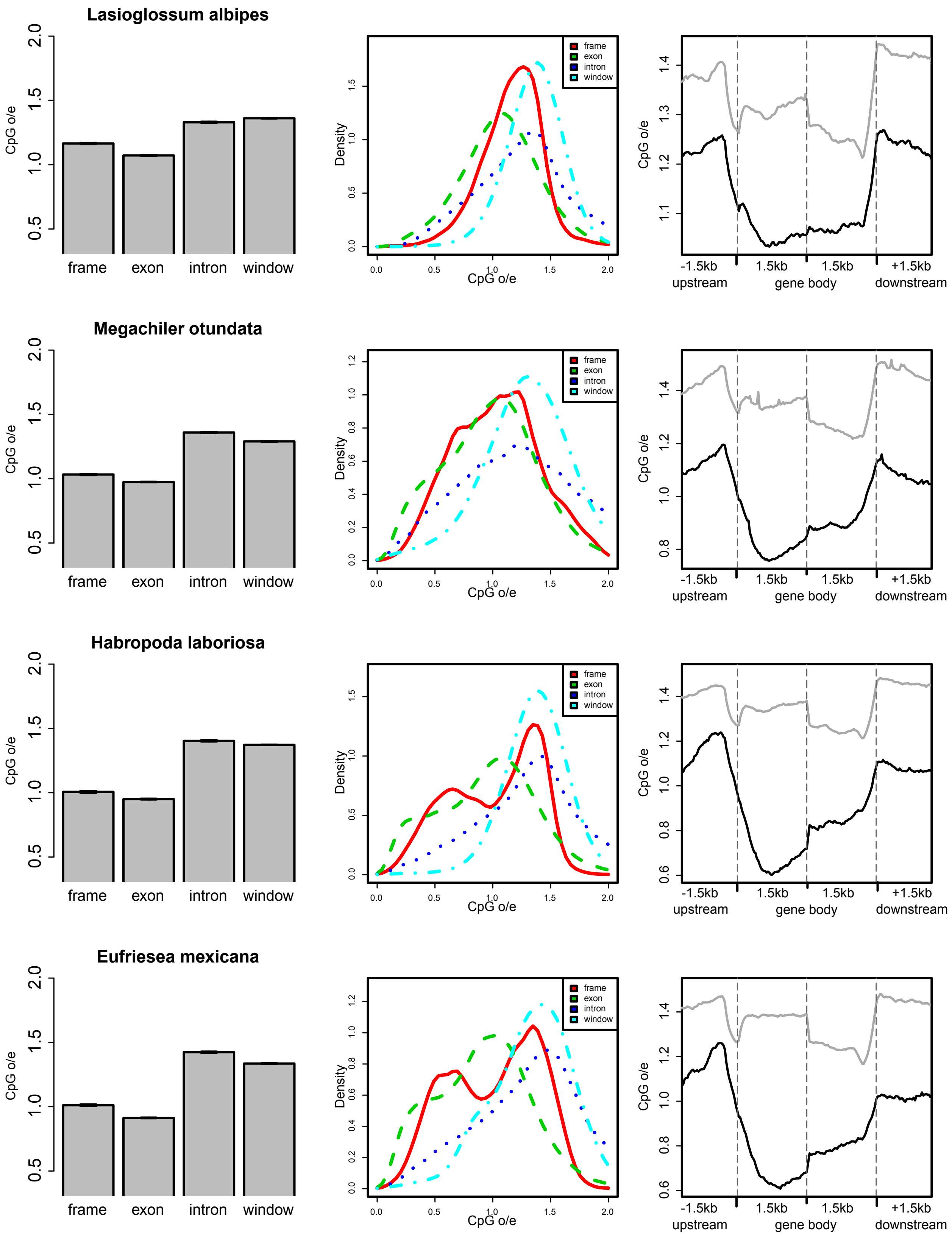
Lasioglossum albipes
Megachiler otundata
Habropoda laboriosa
Eufriesea mexicana
CpG o/e
frame
exon
intron
window
Density
CpG o/e
frame
exon
intron
window
CpG o/e
-1.5kb
upstream
1.5kb
1.5kb
gene body
+1.5kb
downstream

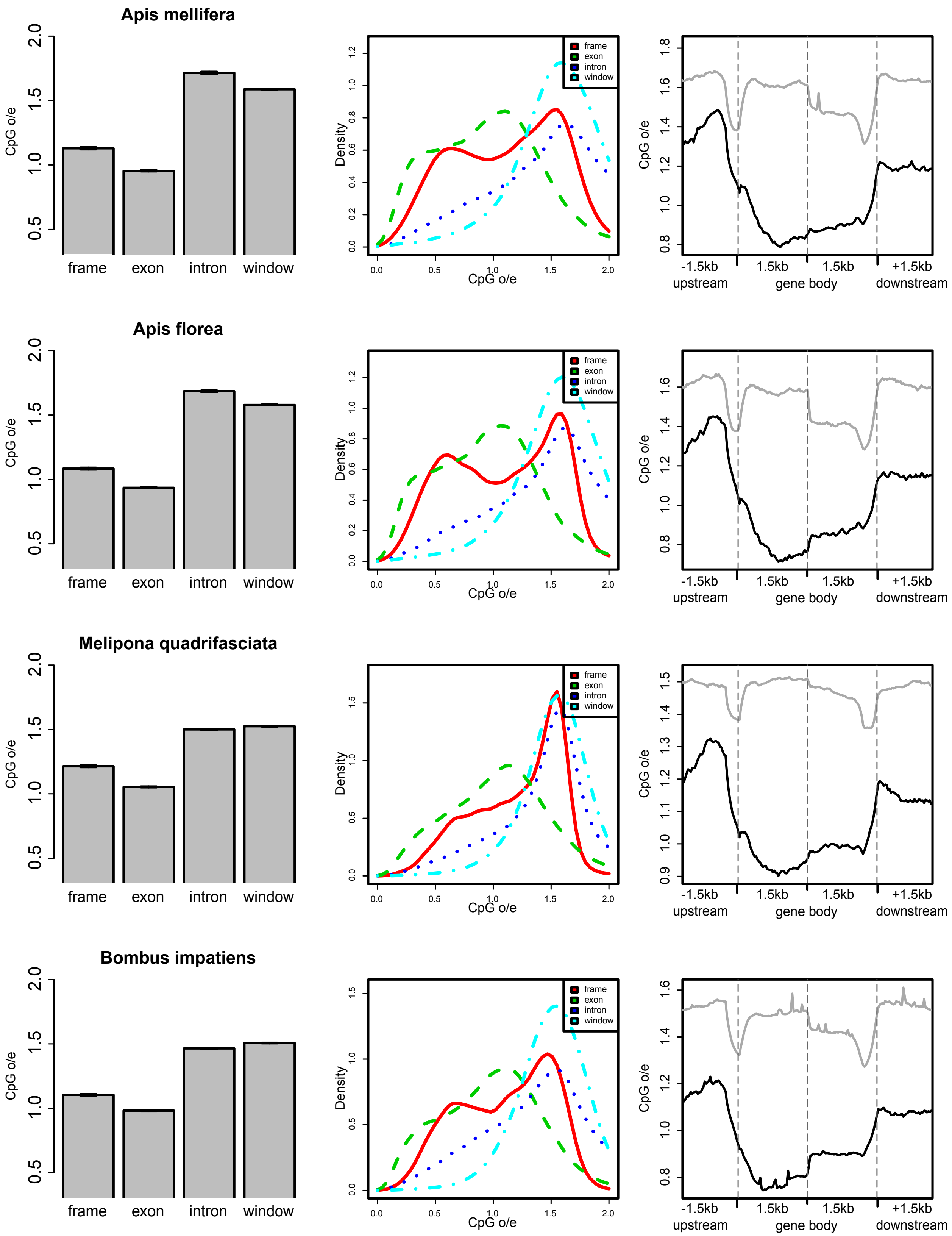
Apis mellifera
Apis florea
Melipona quadrifasciata
Bombus impatiens
CpG o/e
frame
exon
intron
window
Density
CpG o/e
-1.5kb
upstream
1.5kb
gene body
1.5kb
+1.5kb
downstream

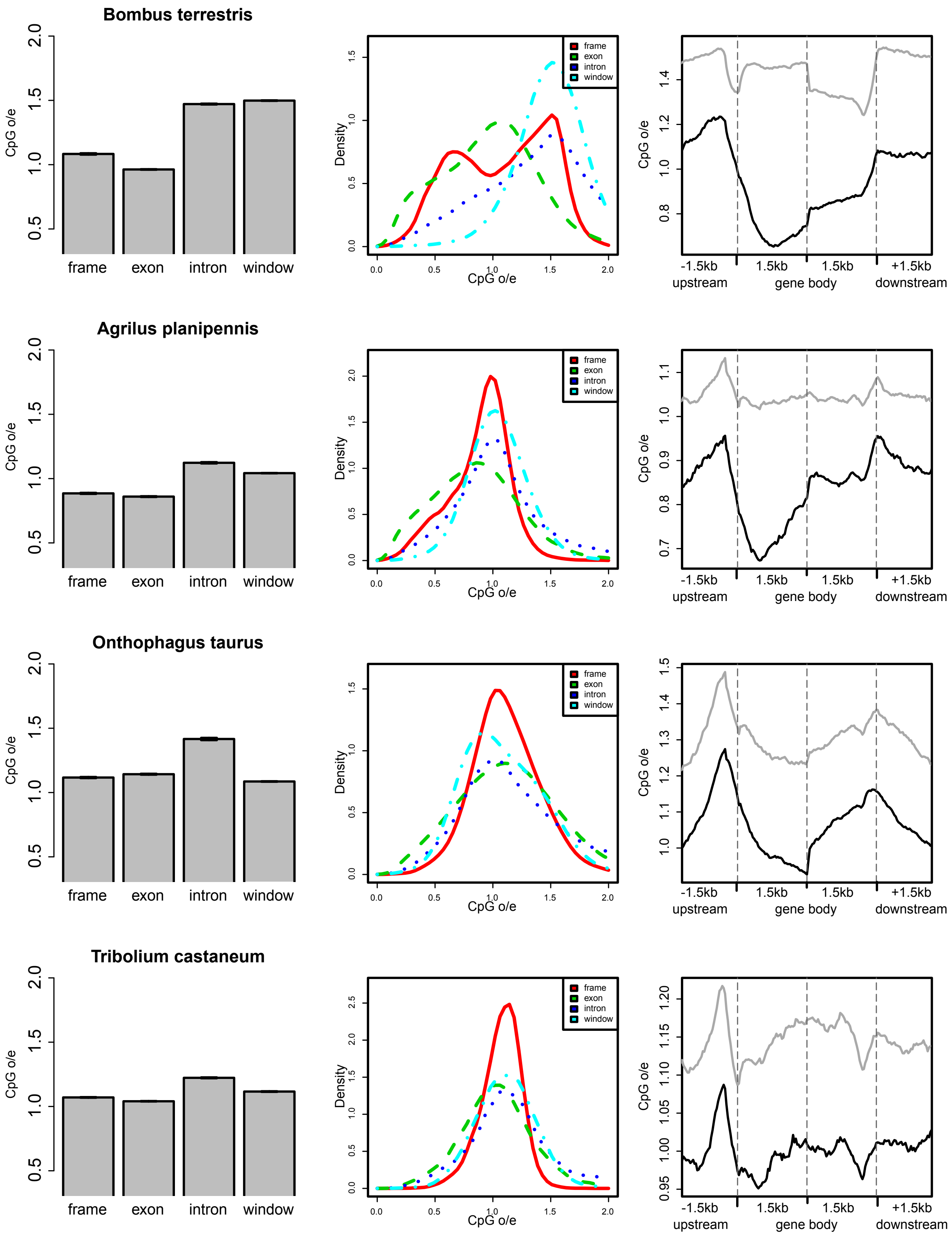
Bombus terrestris
Agrilus planipennis
Onthophagus taurus
Tribolium castaneum
CpG o/e
frame
exon
intron
window
Density
upstream
gene body
downstream
-1.5kb
1.5kb
1.5kb
+1.5kb

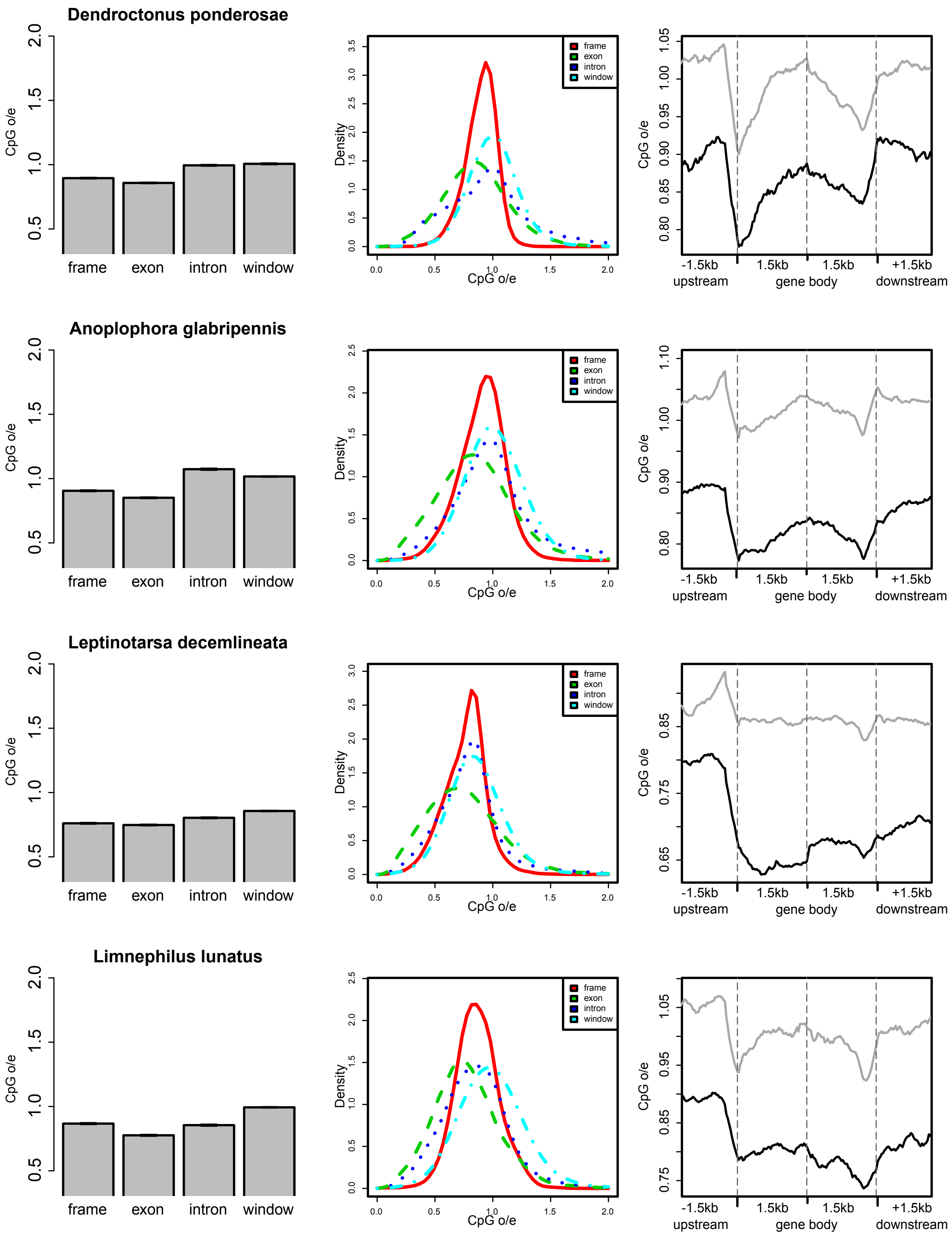
Dendroctonus ponderosae
Anoplophora glabripennis
Leptinotarsa decemlineata
Limnephilus lunatus
CpG o/e
frame
exon
intron
window
Density
-1.5kb
upstream
1.5kb
gene body
1.5kb
+1.5kb
downstream

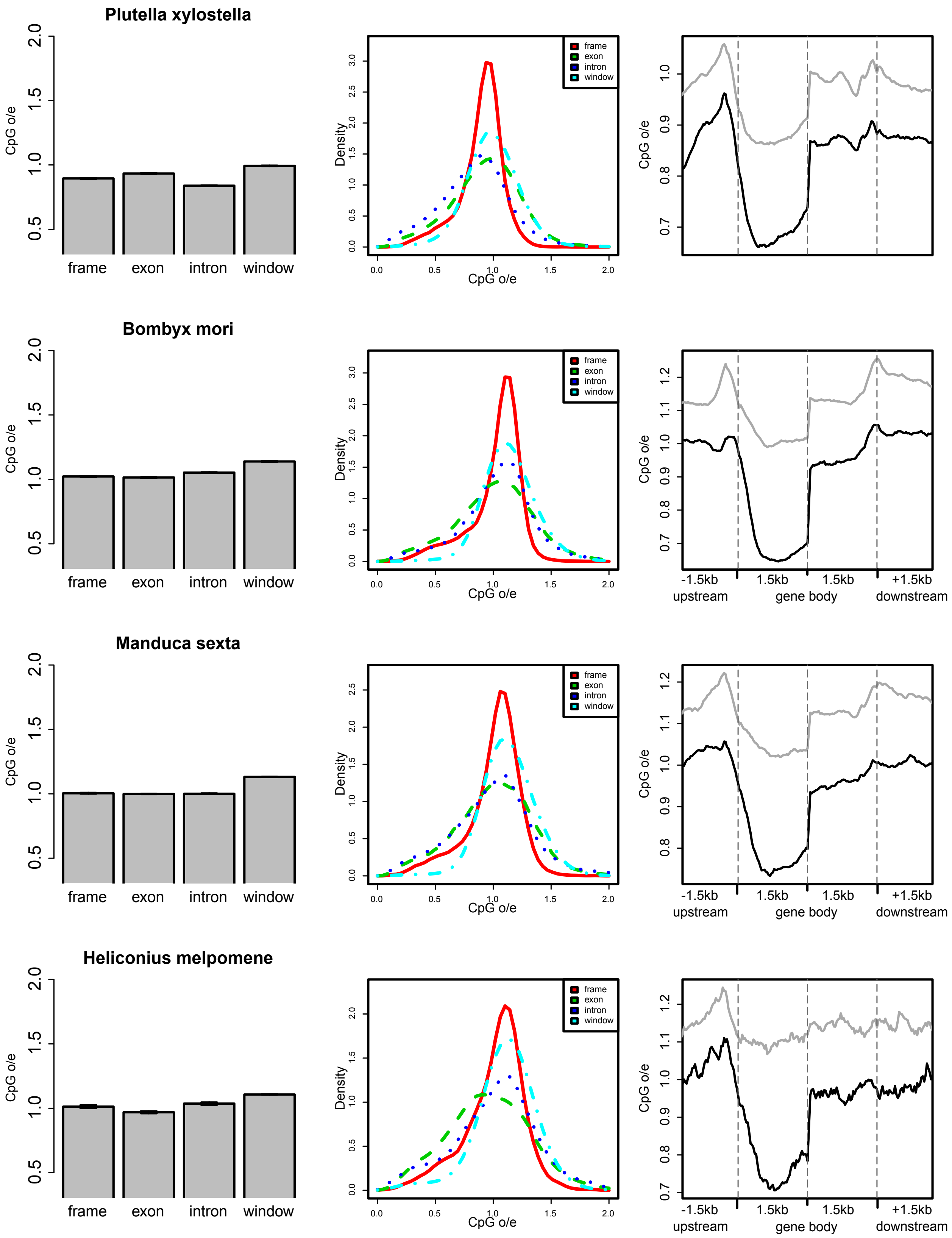
Plutella xylostella
Bombyx mori
Manduca sexta
Heliconius melpomene
CpG o/e
frame
exon
intron
window
Density
-1.5kb
upstream
1.5kb
gene body
1.5kb
+1.5kb
downstream

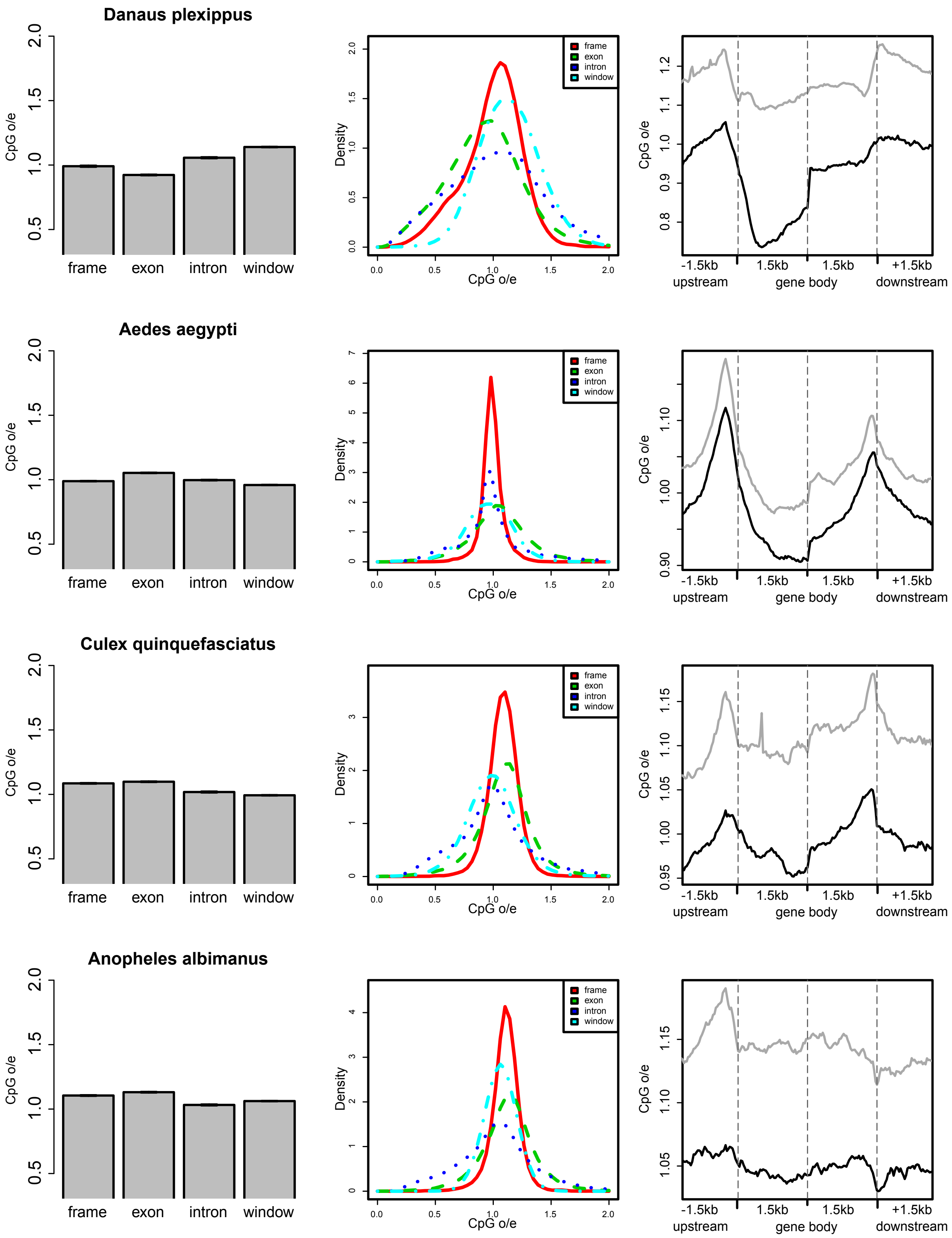
Danaus plexippus
Aedes aegypti
Culex quinquefasciatus
Anopheles albimanus
CpG o/e
frame
exon
intron
window
Density
CpG o/e
frame
exon
intron
window
CpG o/e
-1.5kb
upstream
1.5kb
1.5kb
gene body
+1.5kb
downstream

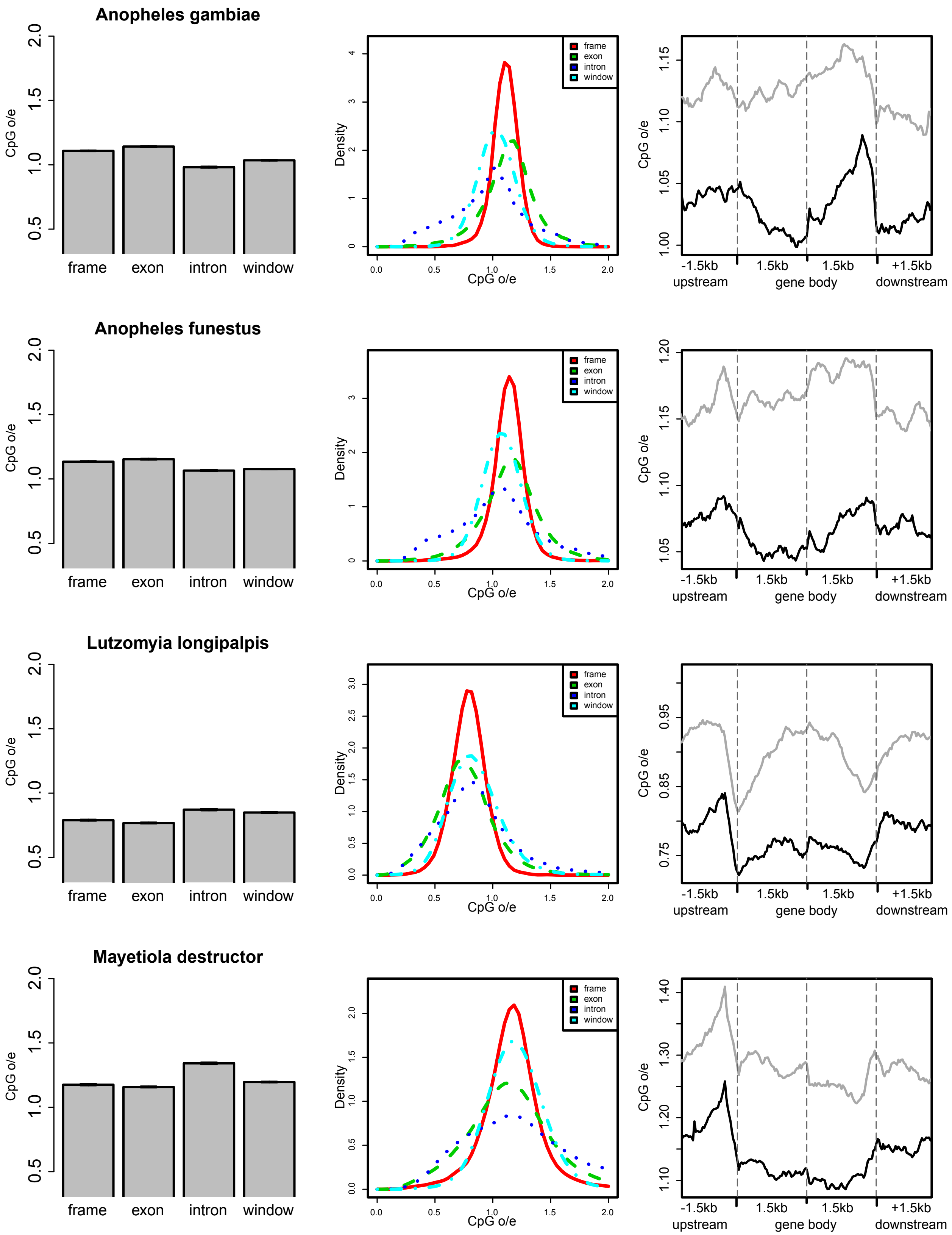
Anopheles gambiae
Anopheles funestus
Lutzomyia longipalpis
Mayetiola destructor
CpG o/e
frame
exon
intron
window
Density
-1.5kb
upstream
1.5kb
gene body
1.5kb
+1.5kb
downstream

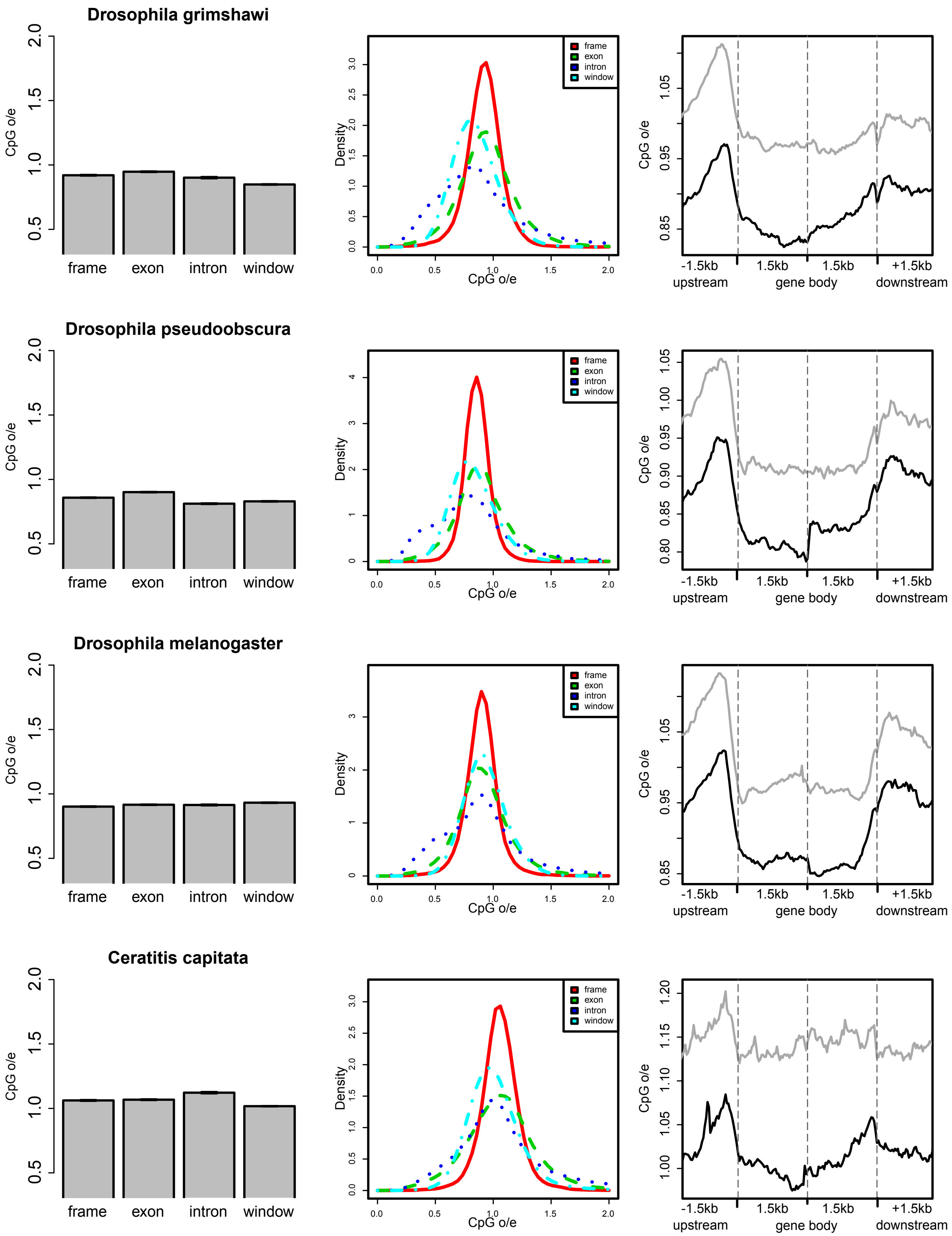
Drosophila grimshawi
Drosophila pseudoobscura
Drosophila melanogaster
Ceratitis capitata
CpG o/e
frame
exon
intron
window
Density
CpG o/e
frame
exon
intron
window
-1.5kb
upstream
1.5kb
1.5kb
gene body
+1.5kb
downstream

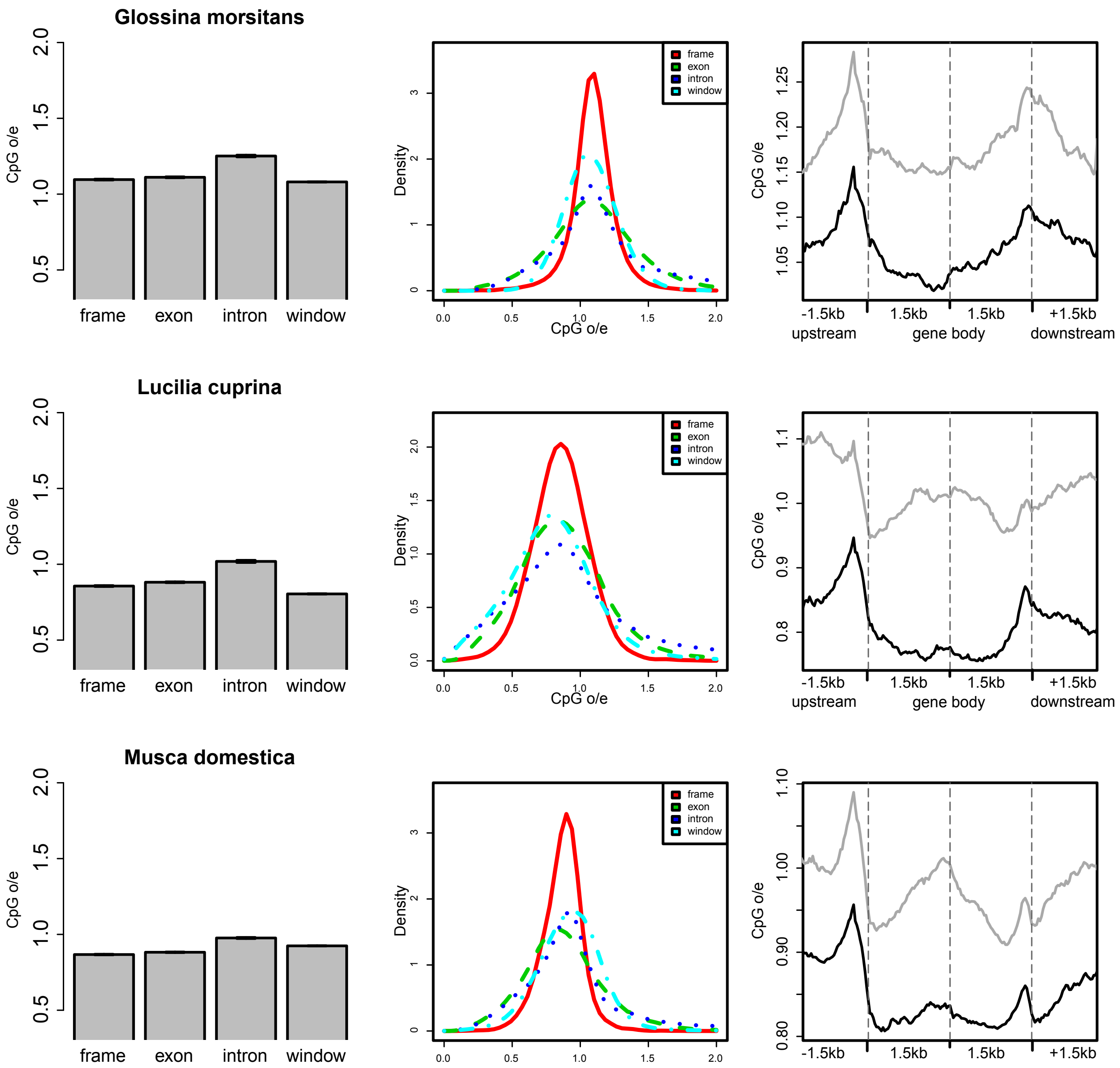
Glossina morsitans
Lucilia cuprina
Musca domestica
CpG o/e
frame
exon
intron
window
Density
-1.5kb
1.5kb
+1.5kb
upstream
gene body
downstream

**Figure S9. Support for 15 alternate crustacean topologies with 3 gene sets.**

Orthologous gene dataset 1 (DS1) consists of the fewest sequences but the most species, while Dataset 2 (DS2) consists of an intermediate number of both, and Dataset 3 (DS3) consists of the most sequences among the fewest species. Each dataset was used to estimate species trees with varying alignment methods, species tree reconstruction methods, and amino acid substitution models (see Methods) for a total 72 species tree estimations. Among the 15 possible topologies for the three crustacean orders (Cladocera [CL], Calanoida [CA], and Amphipoda [A]) and insects (I) we recover 12. A majority of methods support a monophyletic crustacea (T1, T2, and T3).

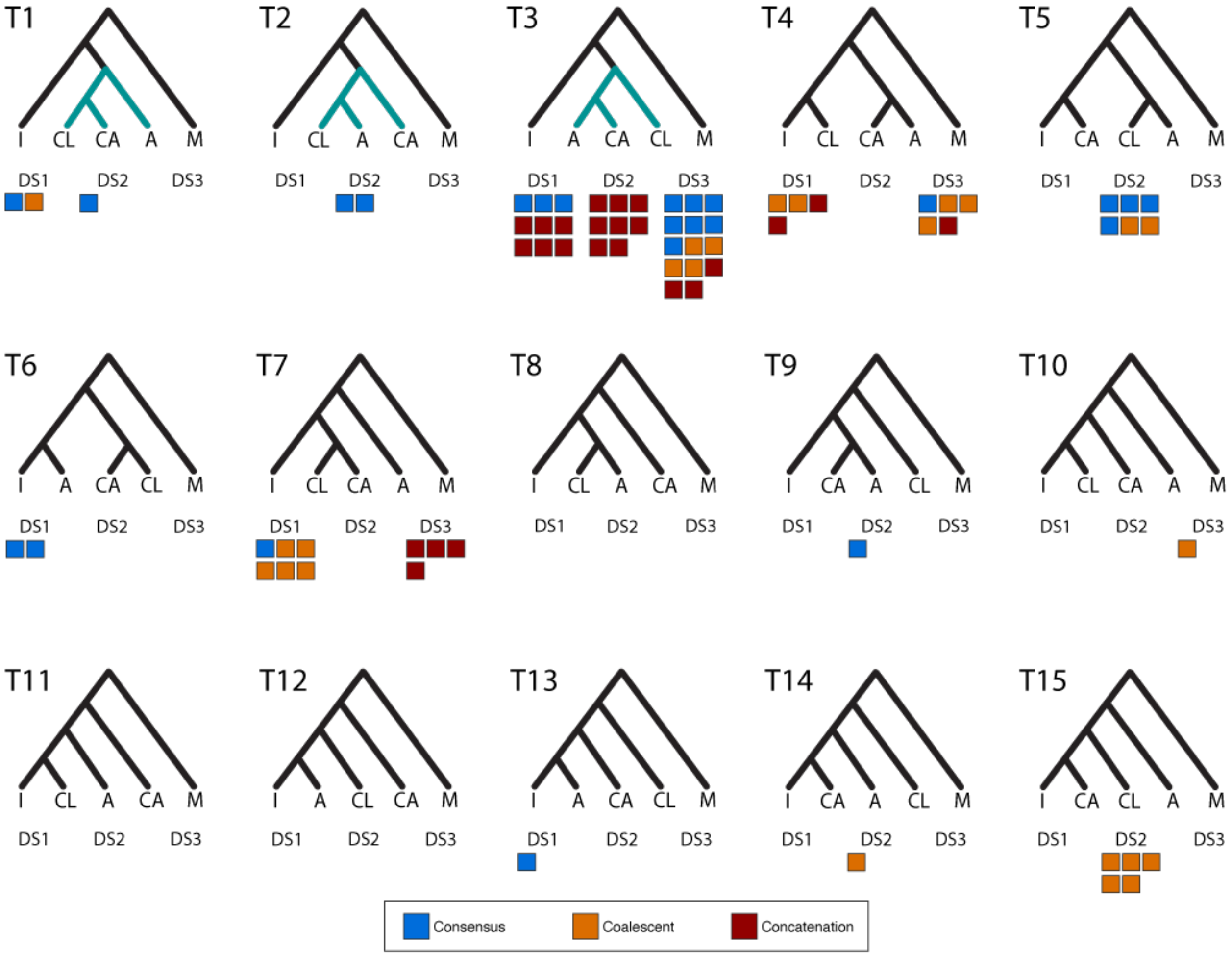

**Figure S10: Novel gene family expansions and extinctions.**

The raw number of gene family emergences (**A**) and extinctions (**B**) for every node in the arthropod phylogeny. Lineages leading to orders represented by multiple species are colored.

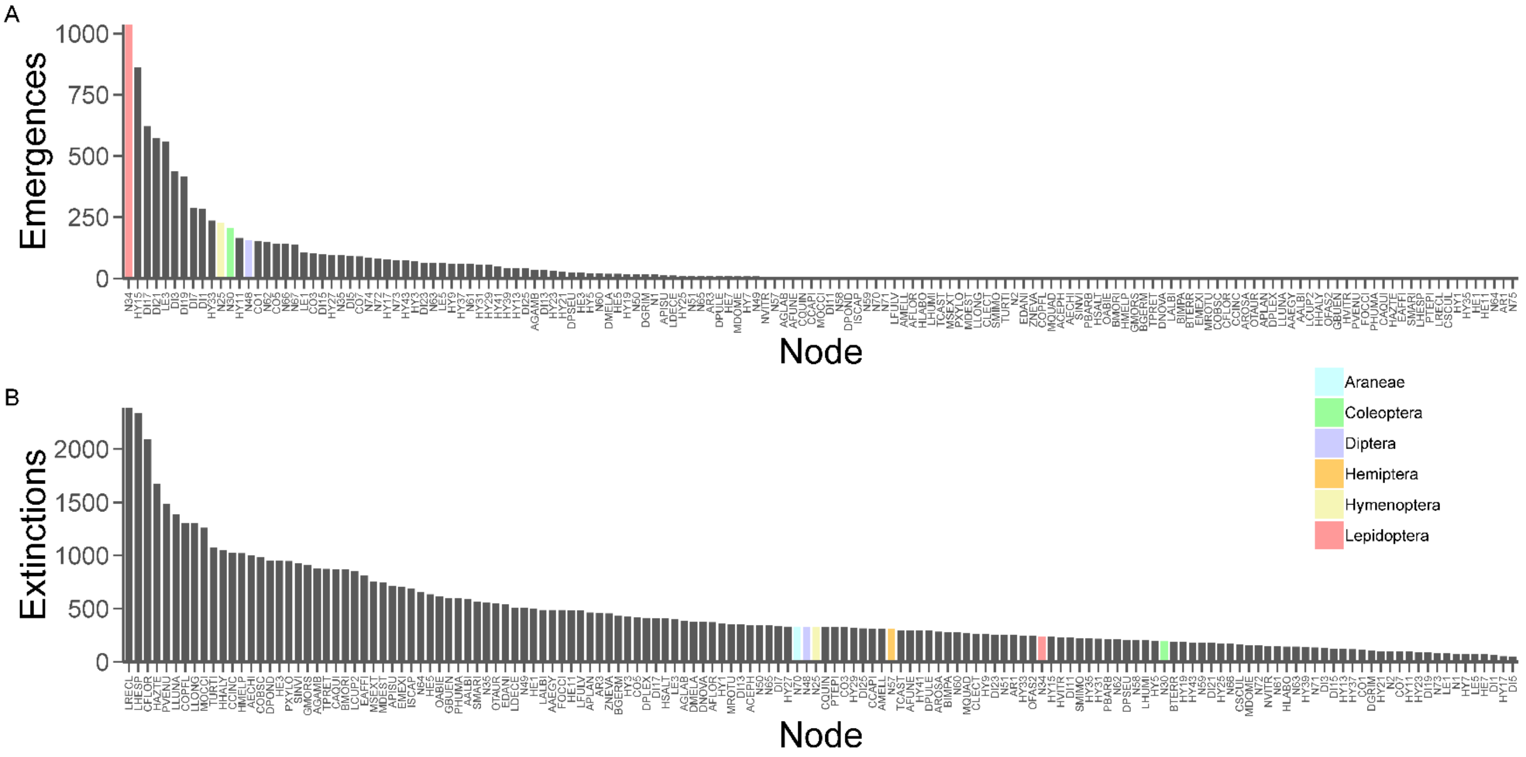

**Figure S11: Araneae tree.**

See Table S27 for Newick format.

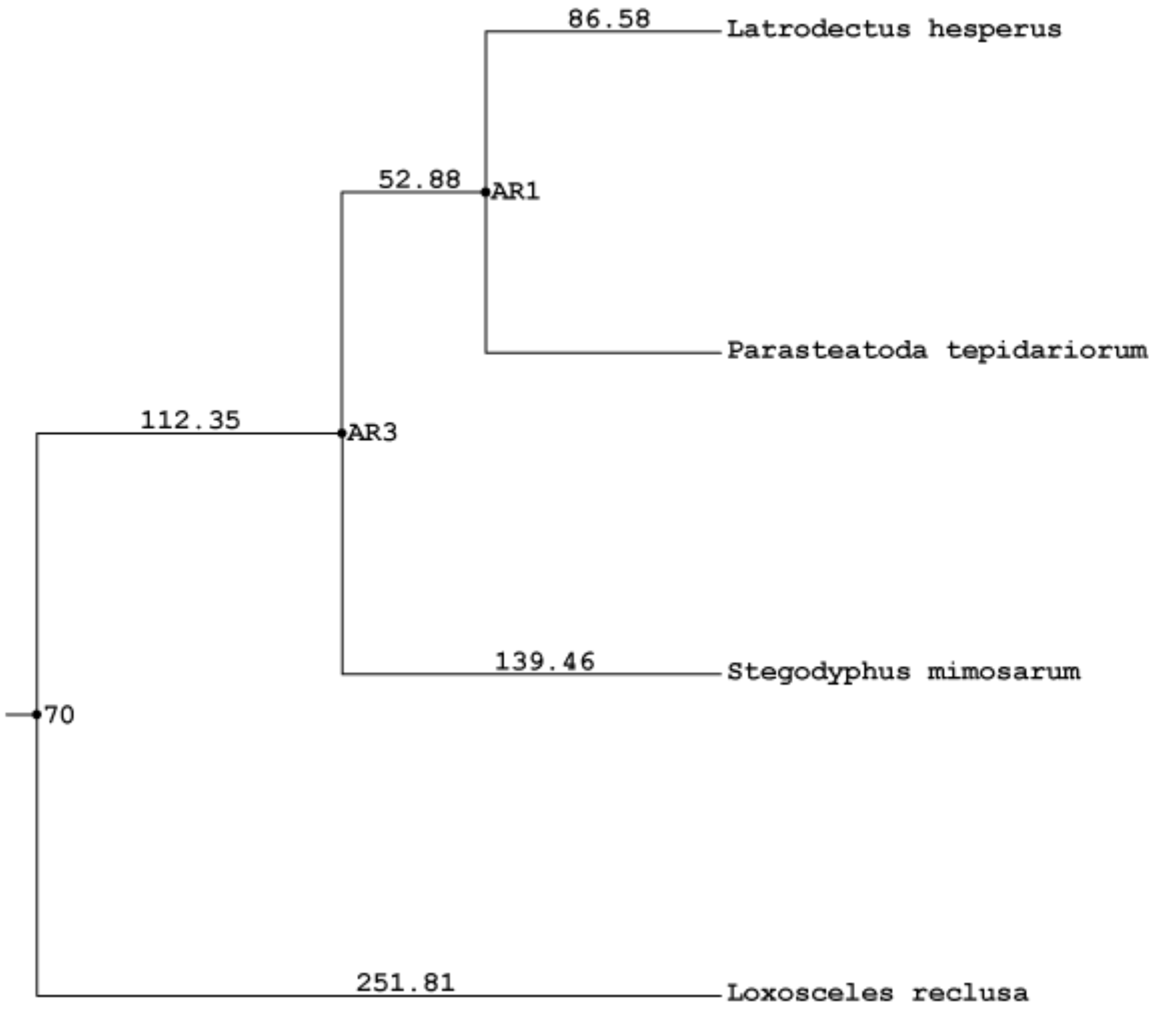

**Figure S12: Hemiptera tree.**

See Table S27 for Newick format.

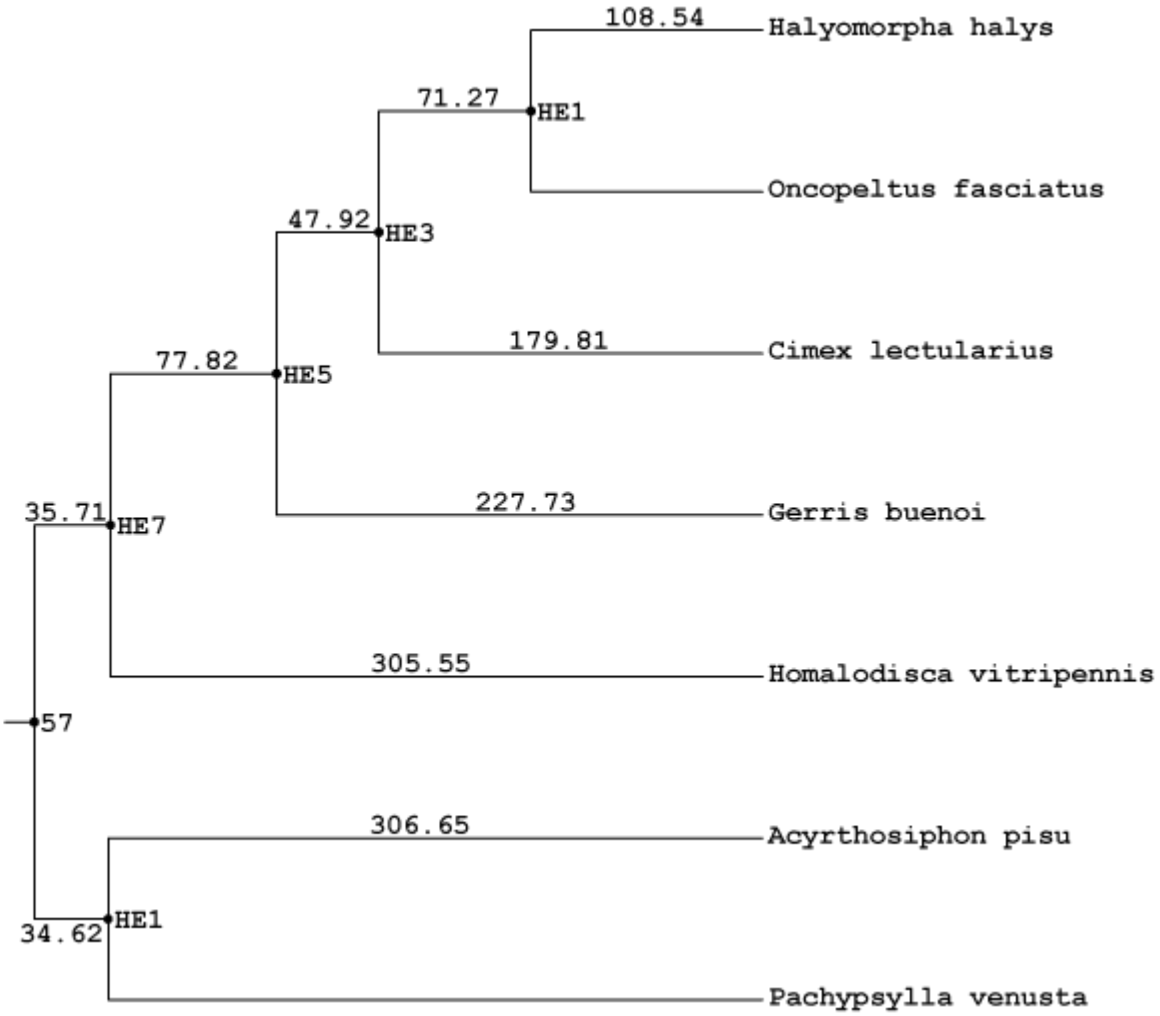

**Figure S13: Hymenoptera tree.**

See Table S27 for Newick format.

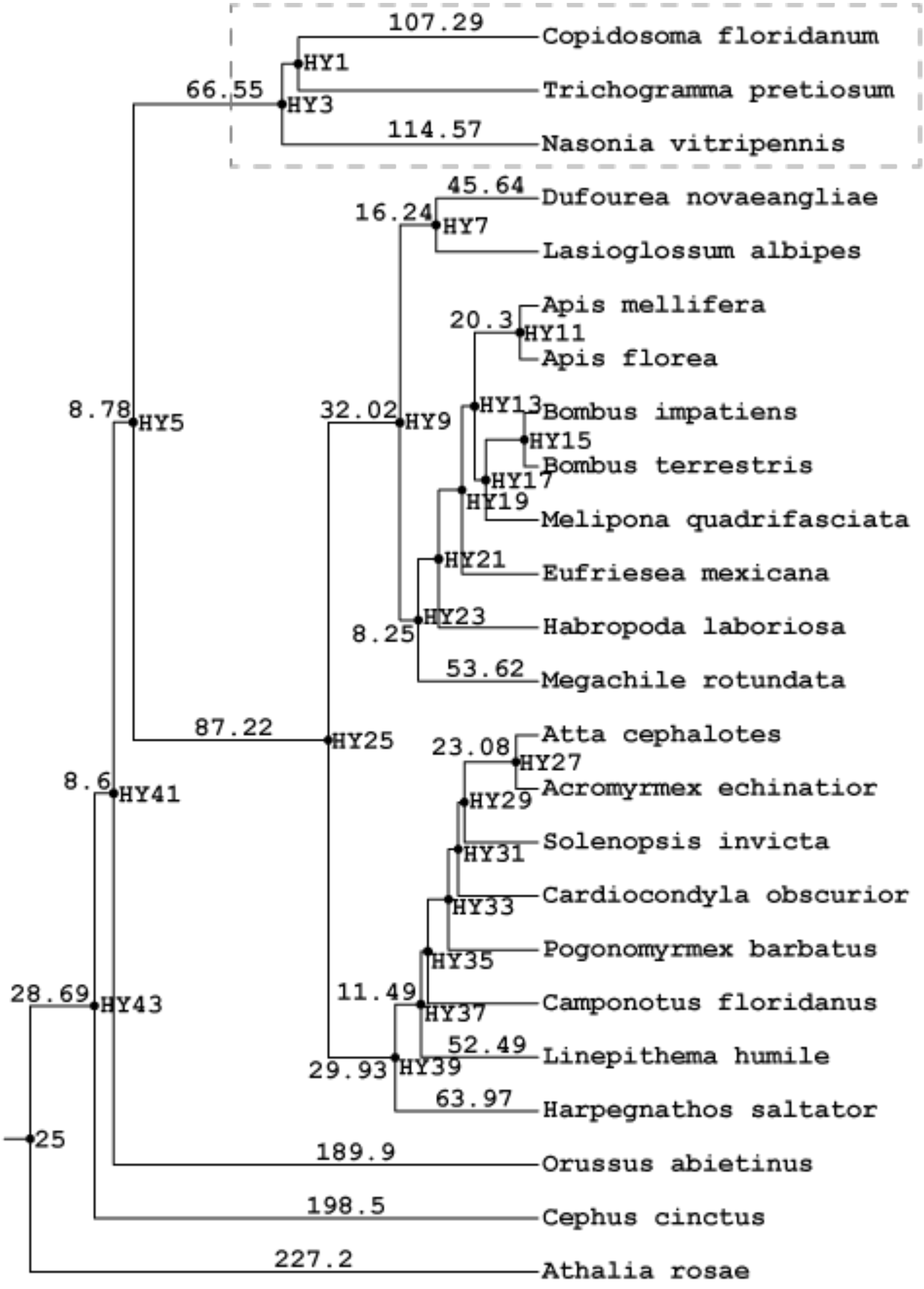

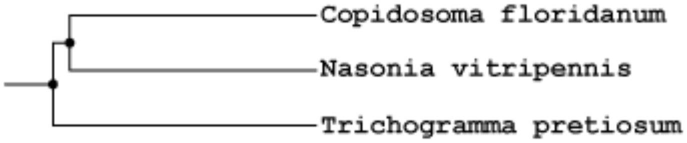

**Figure S14: Coleoptera tree.**

See Table S27 for Newick format.

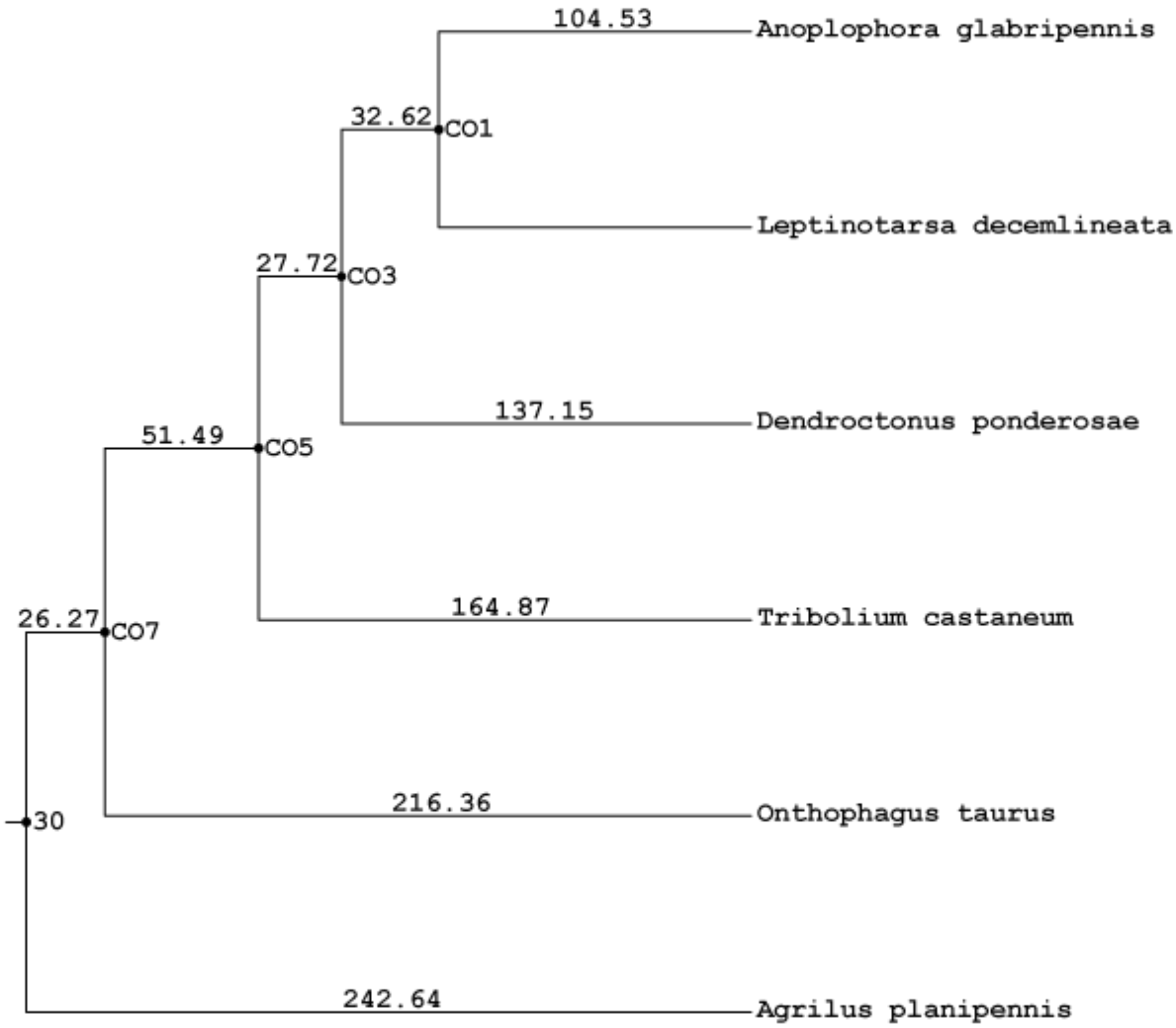

**Figure S15: Lepidoptera tree.**

See Table S27 for Newick format.

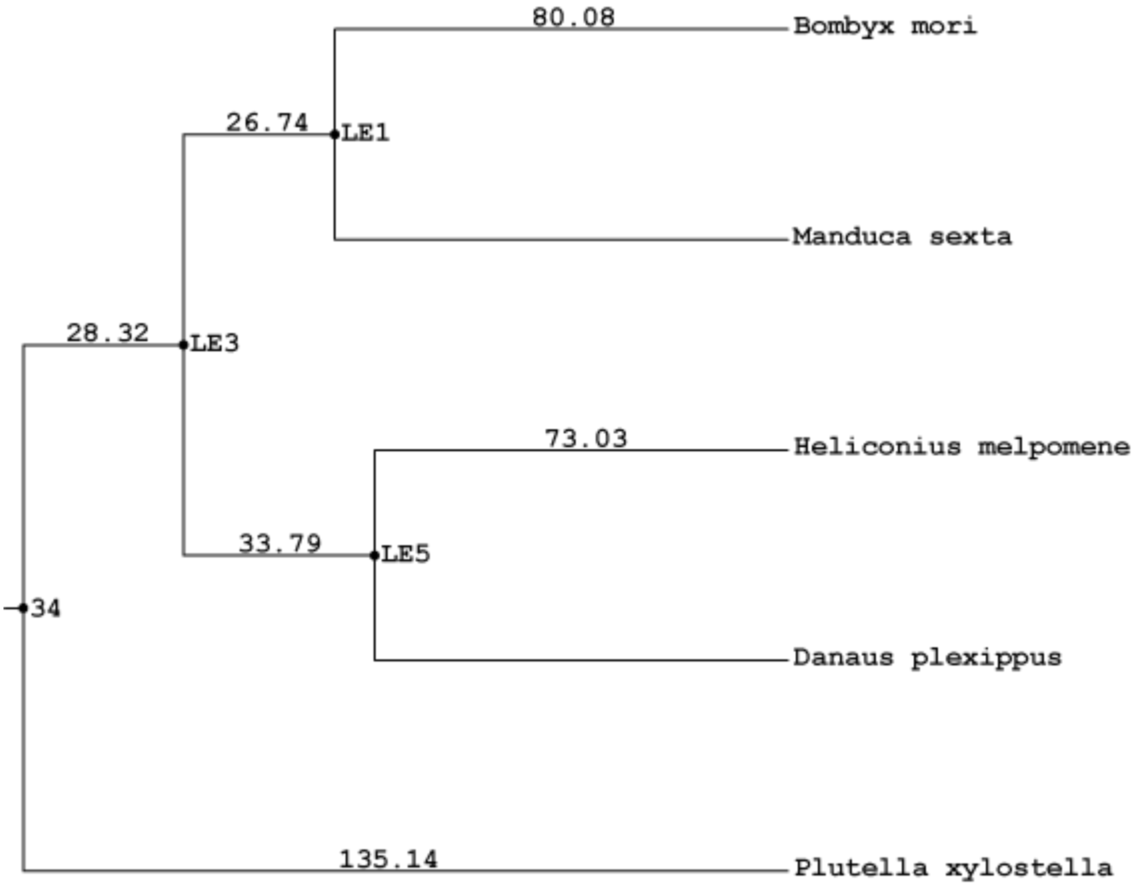

**Figure S16: Diptera tree.**

See Table S27 for Newick format.

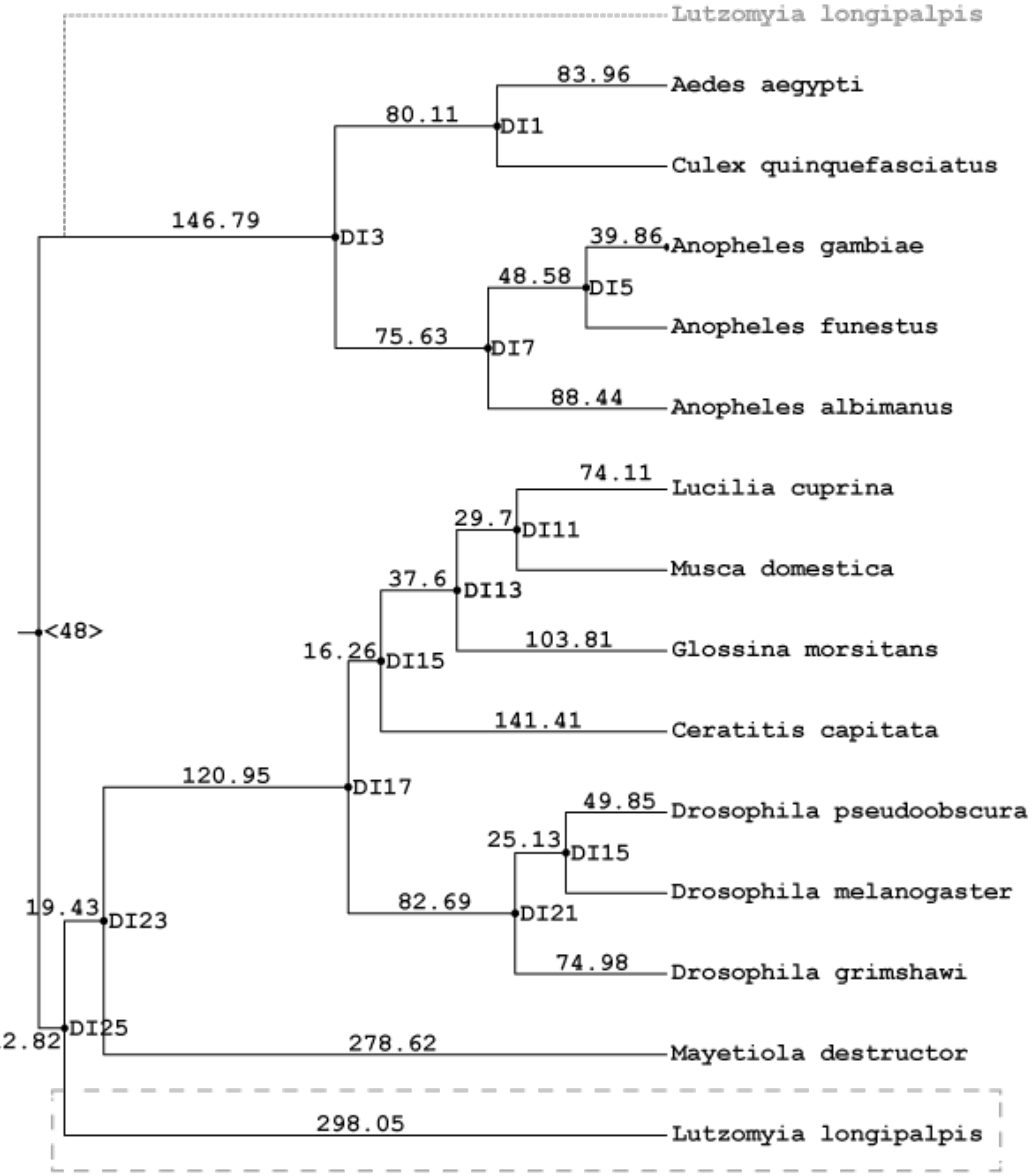

**Figure S17: Fig 2. with all nodes labeled.**

See Table S27 for Newick format.

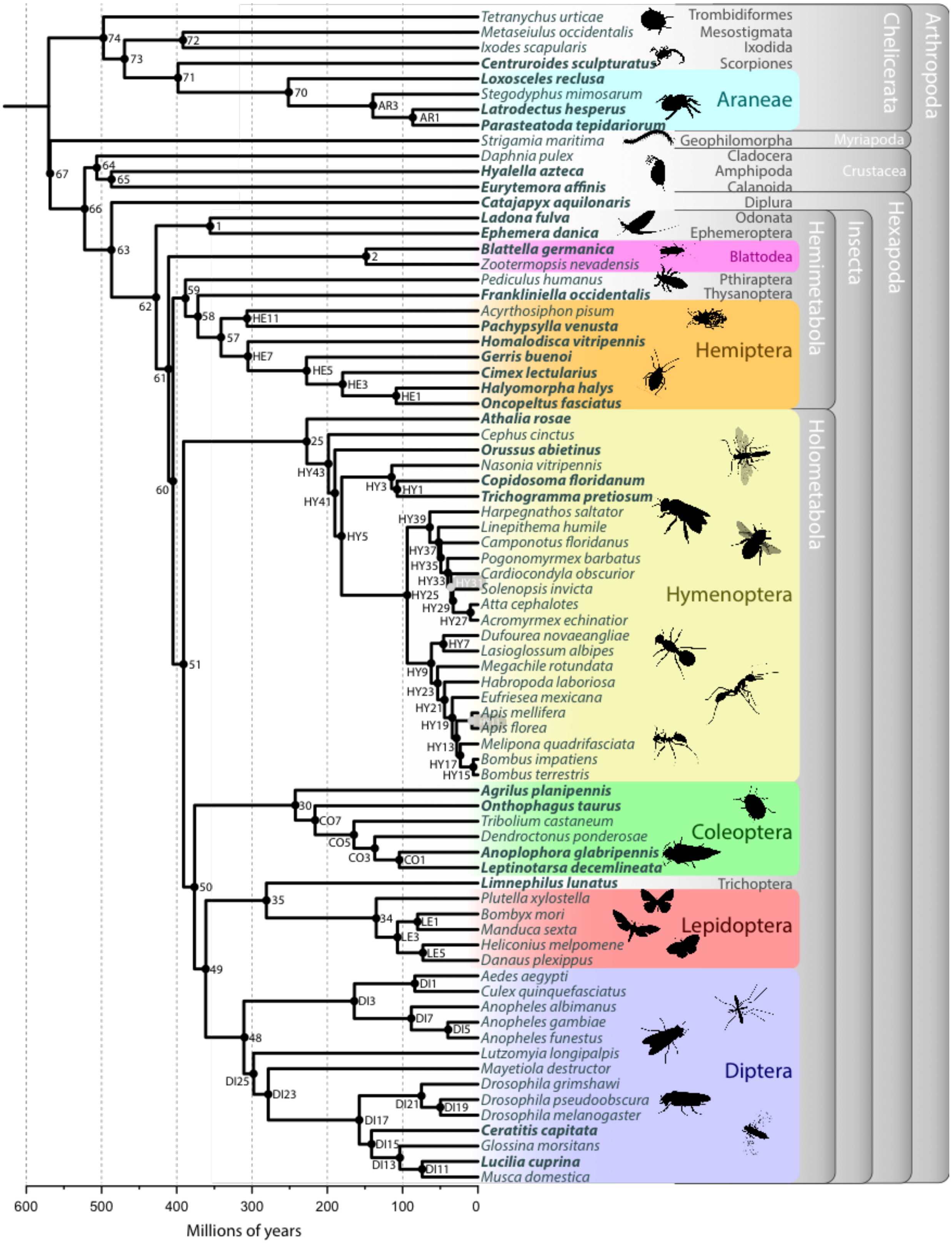

**Figure S18: Gene family emergences vs. gene family extinctions.**

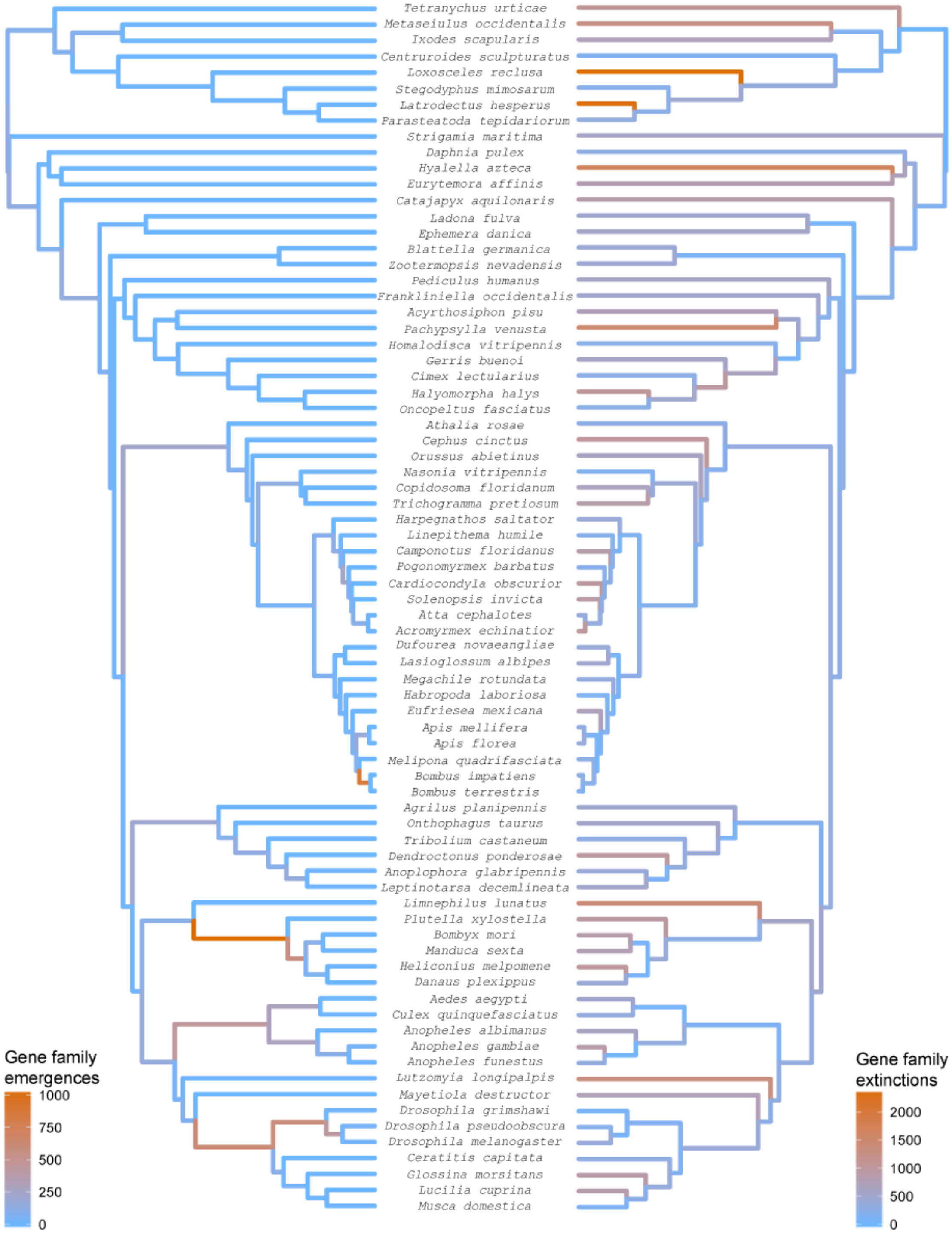

**Figure S19: Rapid gene family expansions vs. rapid gene family contractions.**

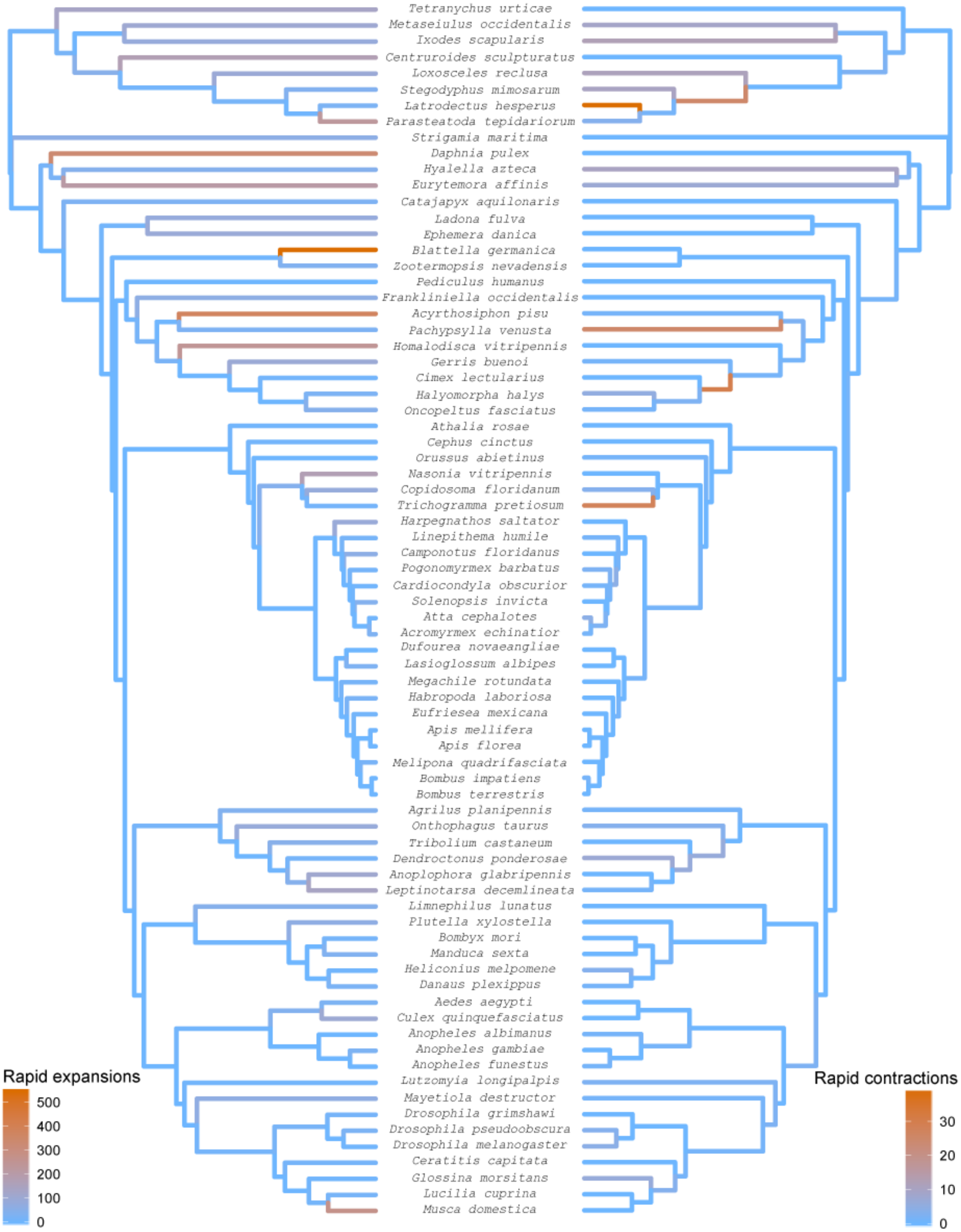

#### Figure S20. Rates of gene family emergences and extinctions.

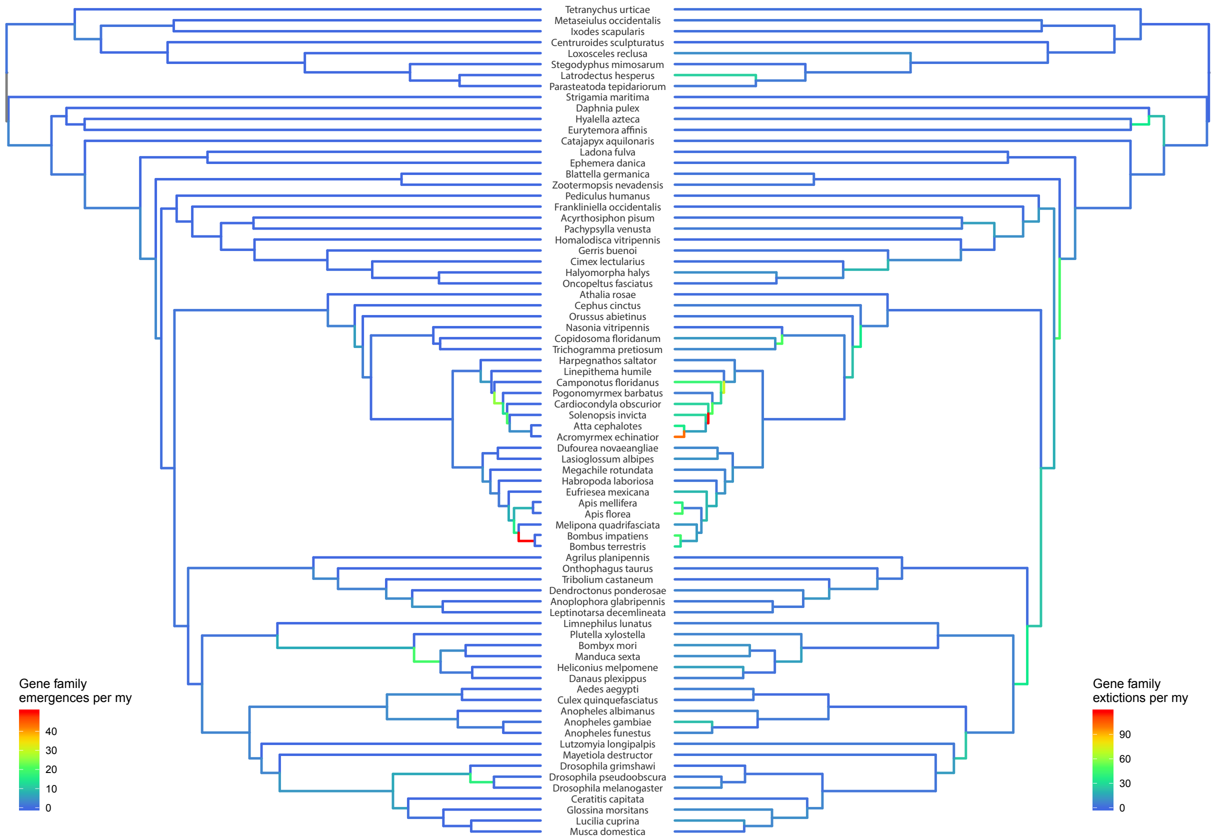

**Figure S21. Distribution of domain rearrangement events.**

Pie charts show the distribution of domain rearrangement events per node (according to color scheme at the top), while the number of reconstructed domain rearrangement events per node is shown in digit representation on the right.

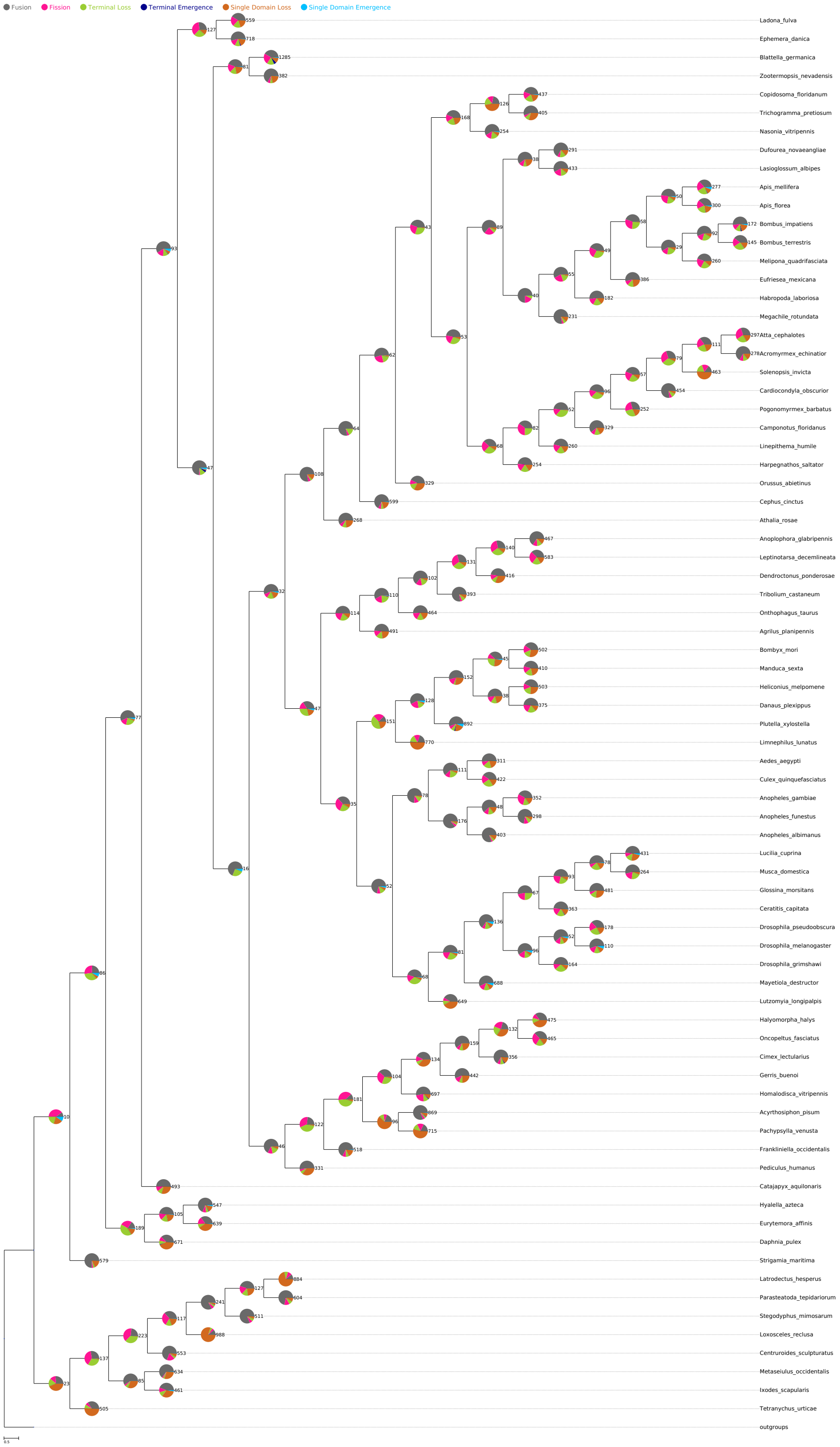

### Figure S22. Distribution of fusion events

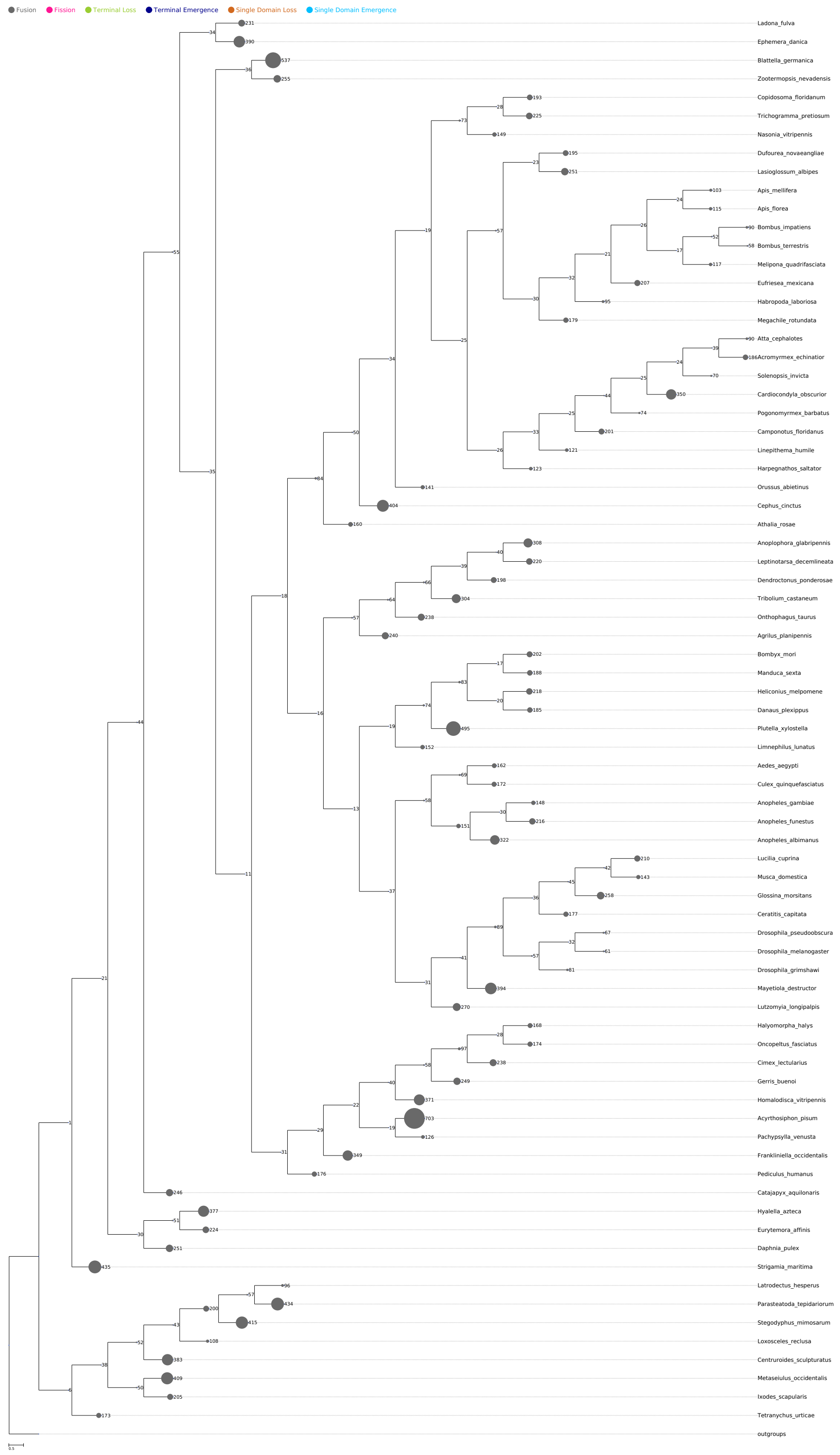

### Figure S23. Distribution of fission events.

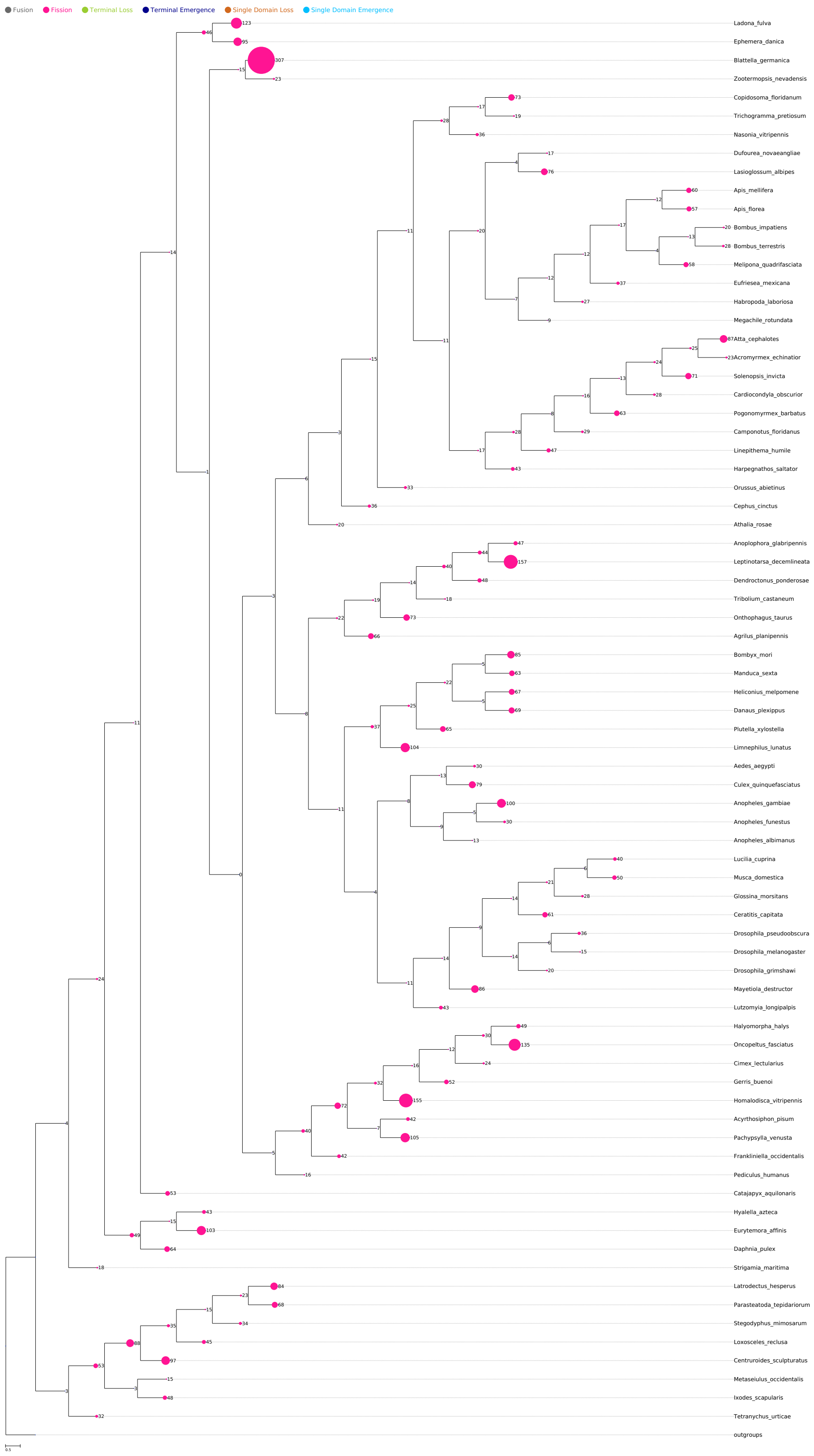

### Figure S24. Distribution of terminal loss events

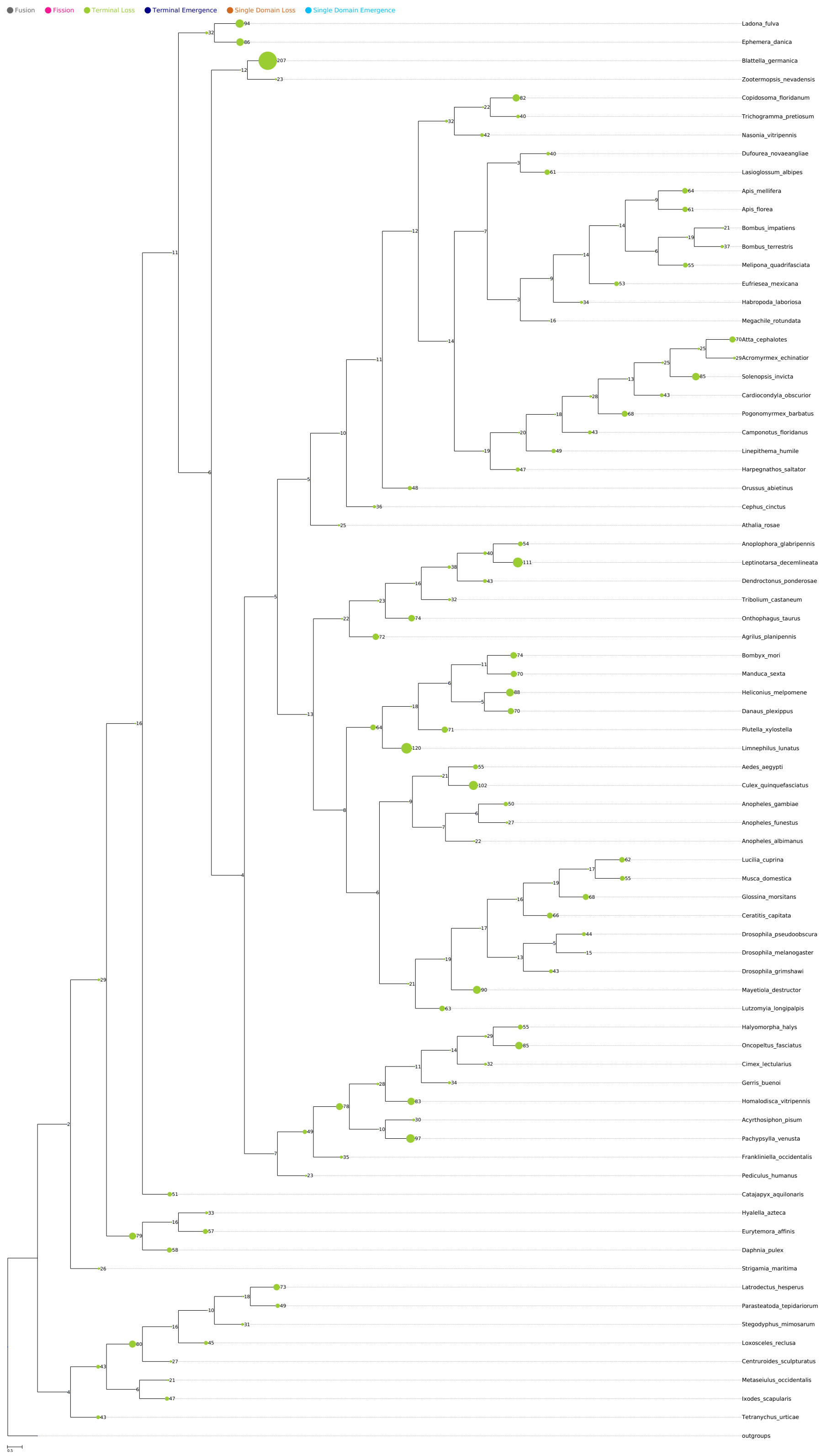

### Figure S25. Distribution of terminal emergence events

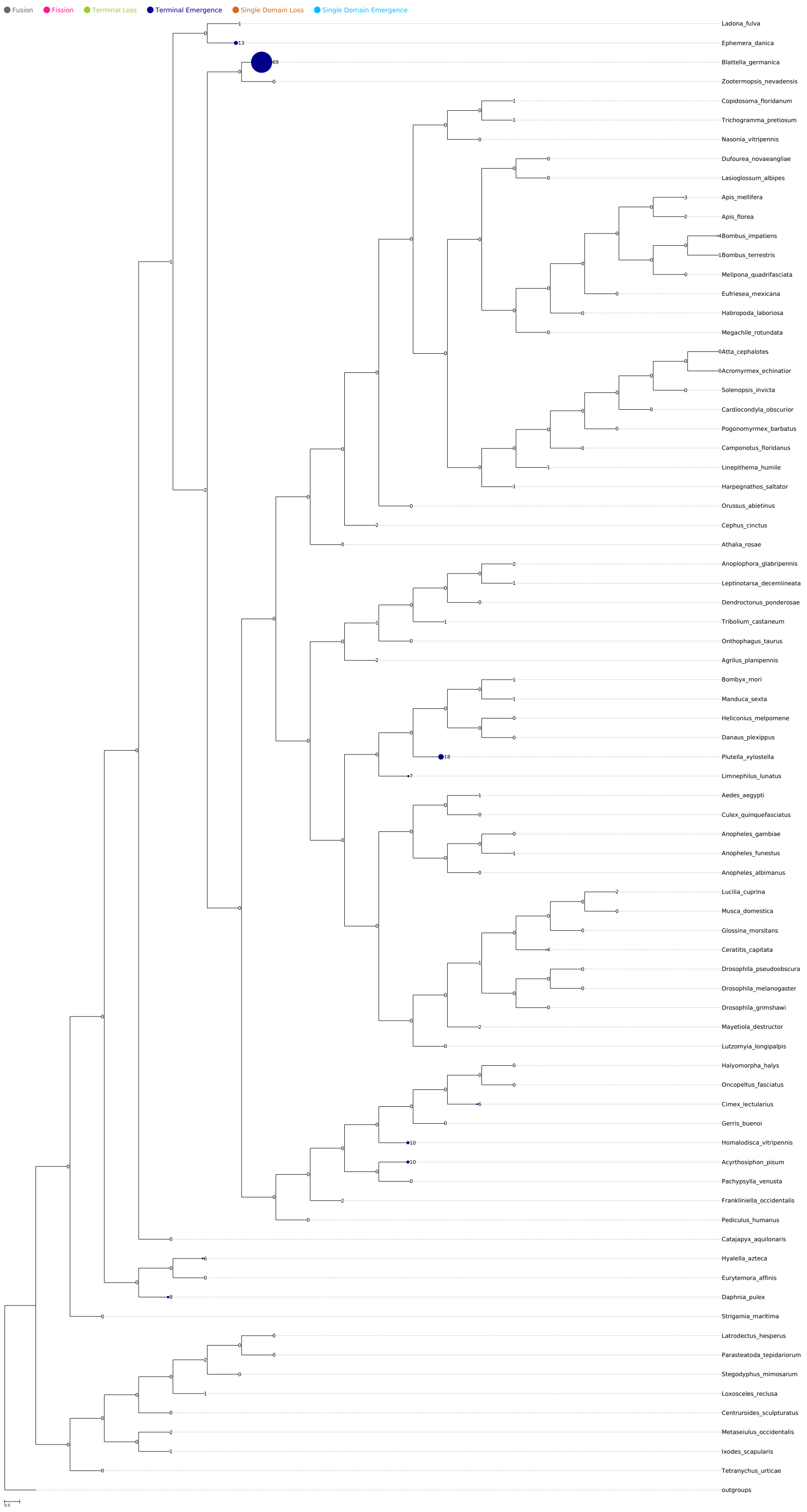

### Figure S26. Distribution of single domain loss events

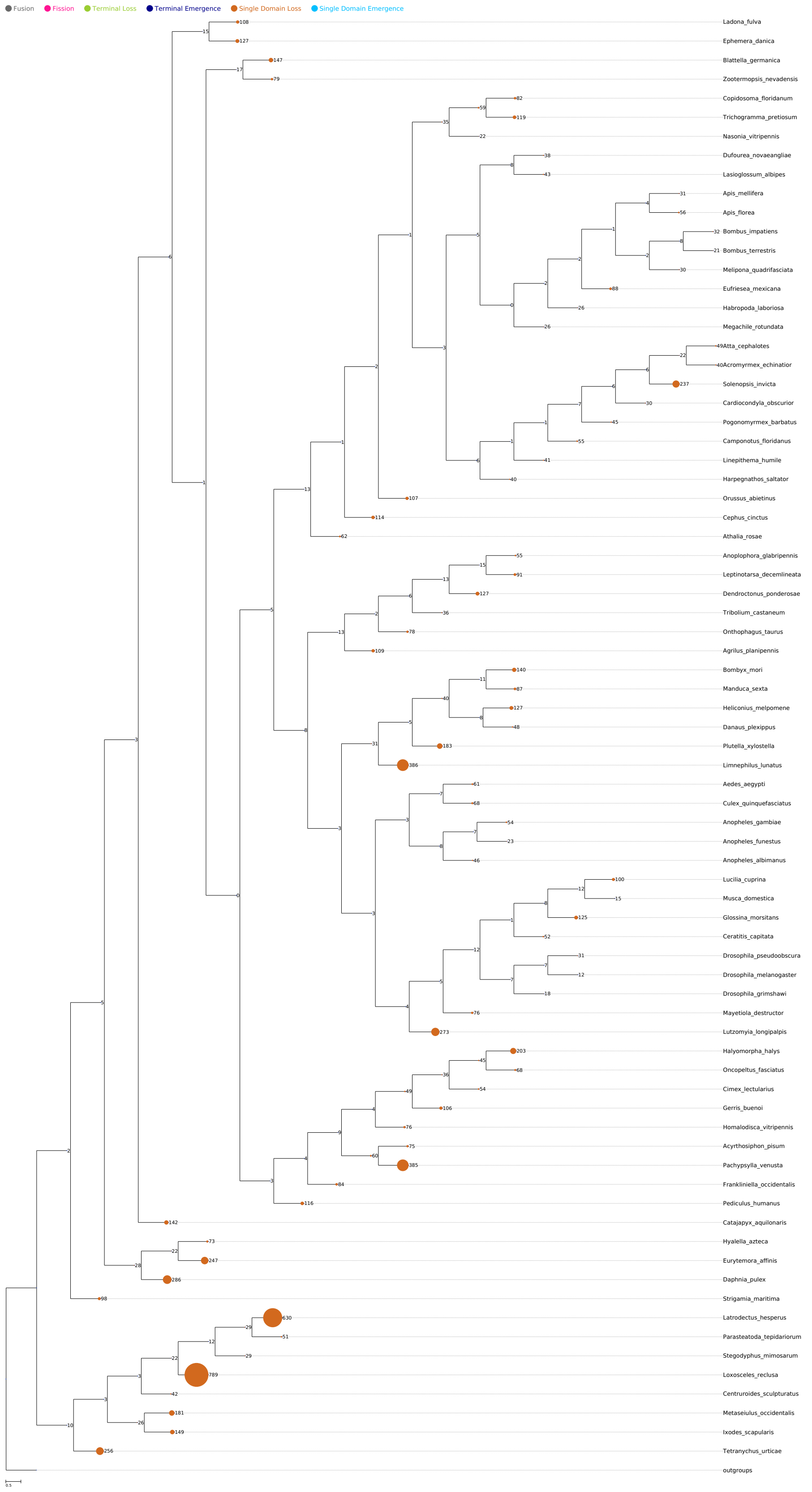

### Figure S27. Distribution of single domain emergence events

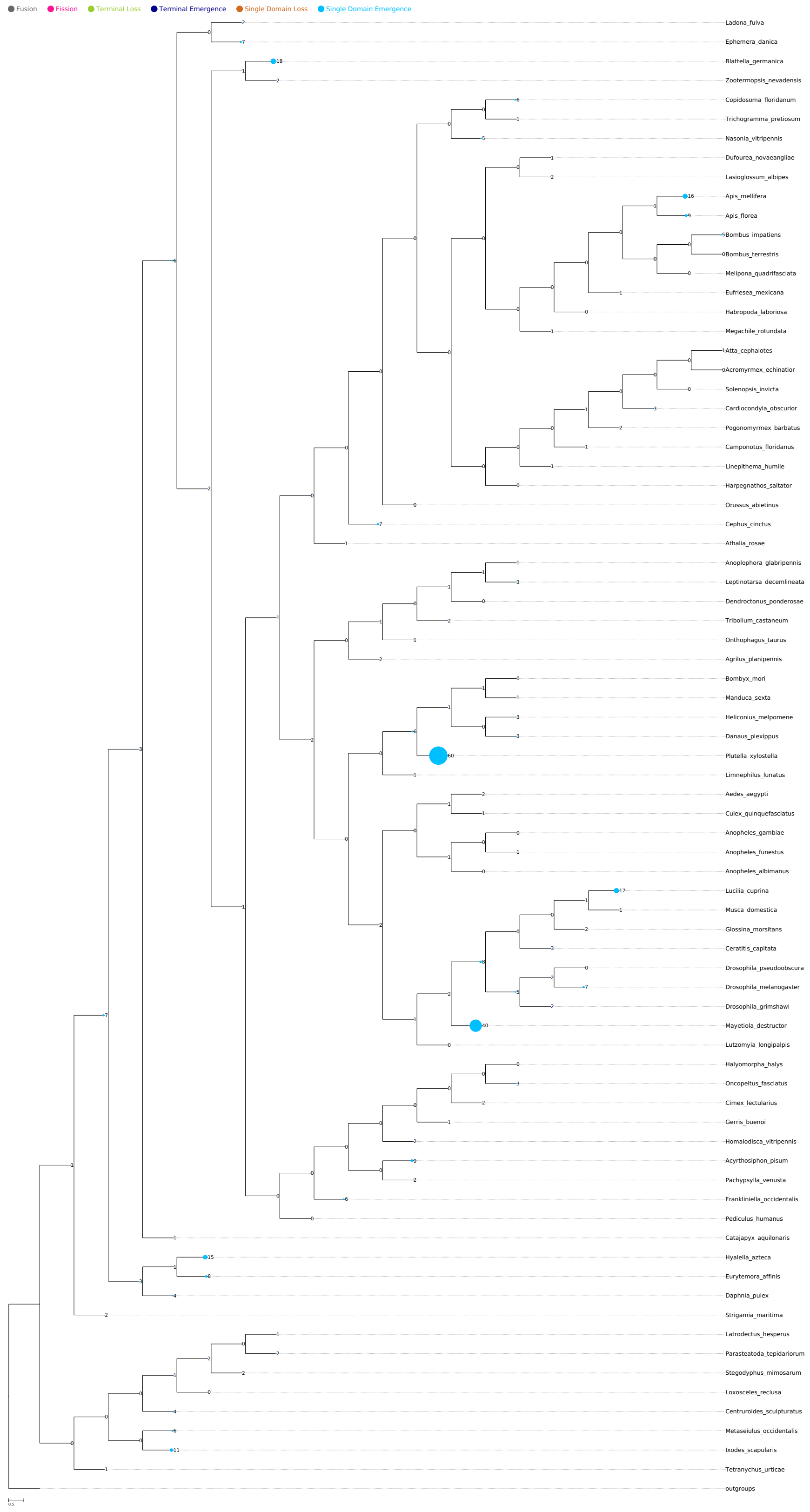

**Figure S28. Substitution, gene gain/loss, and domain rearrangement rates compared.**

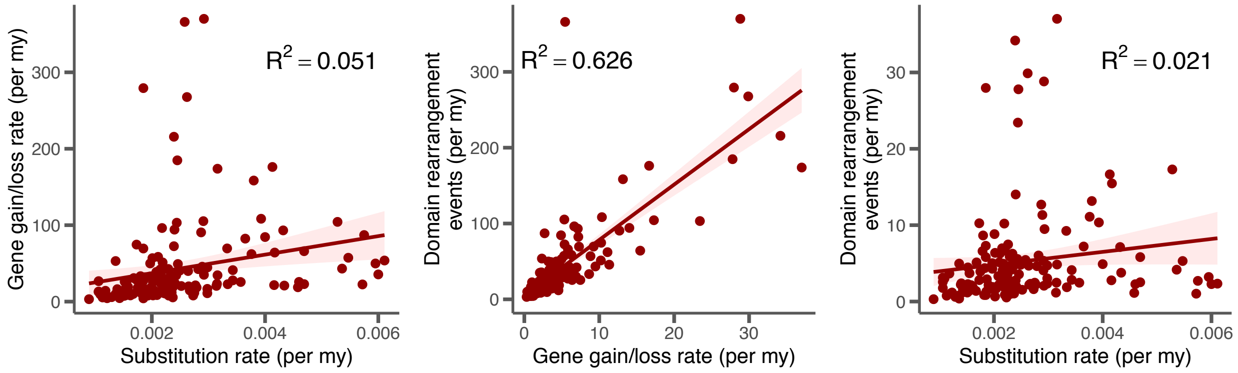

**Figure S29. Significant GO terms in gained domain arrangements.**

Terms of the ontology of "Biological process" (red) "Molecular function" (blue) are shown.

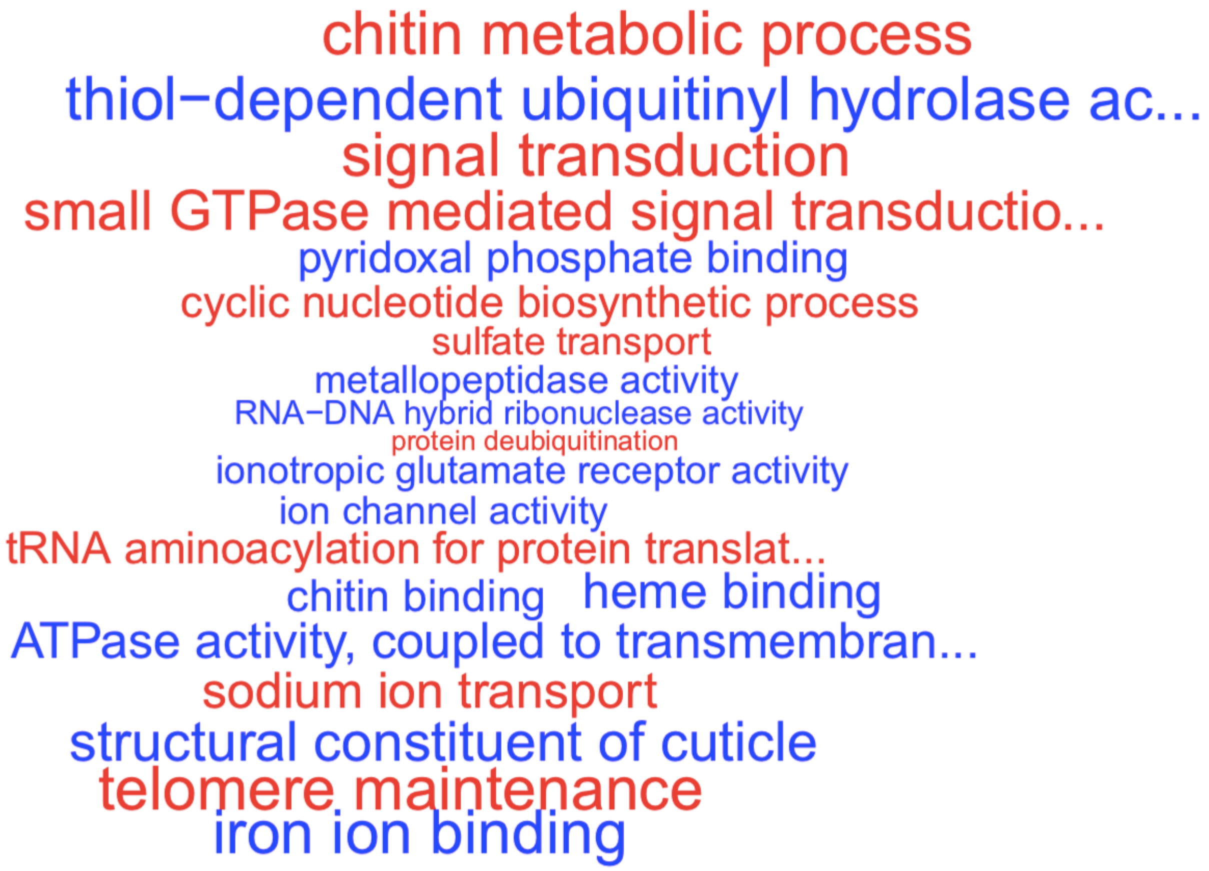

#### Figure S30. Dipteran gene content descriptive statistics.

**A**

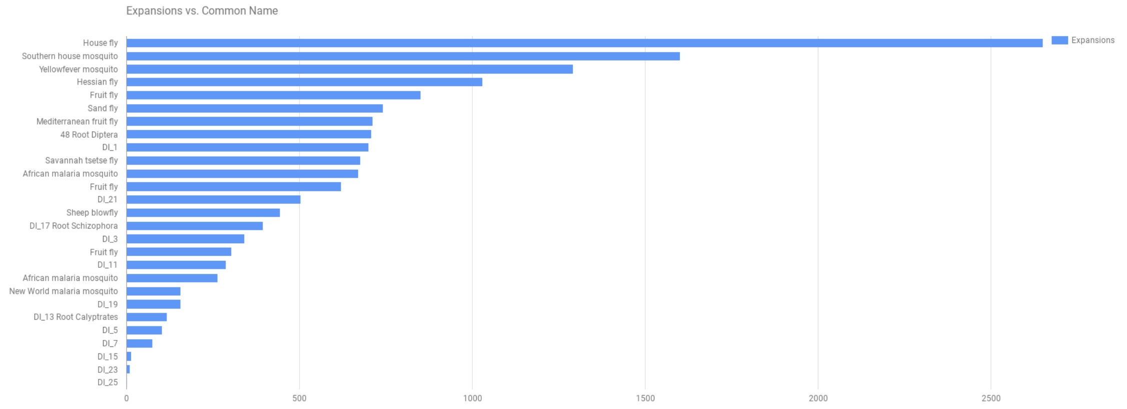

**B**

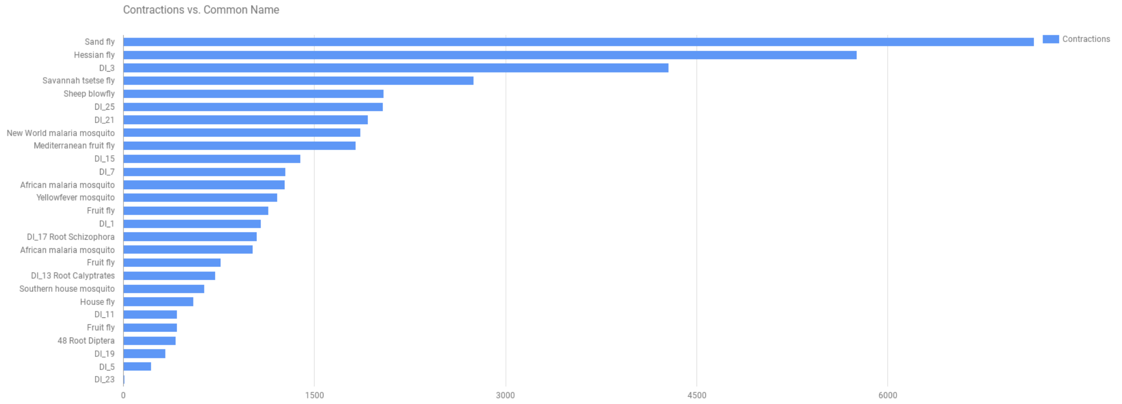

**C**

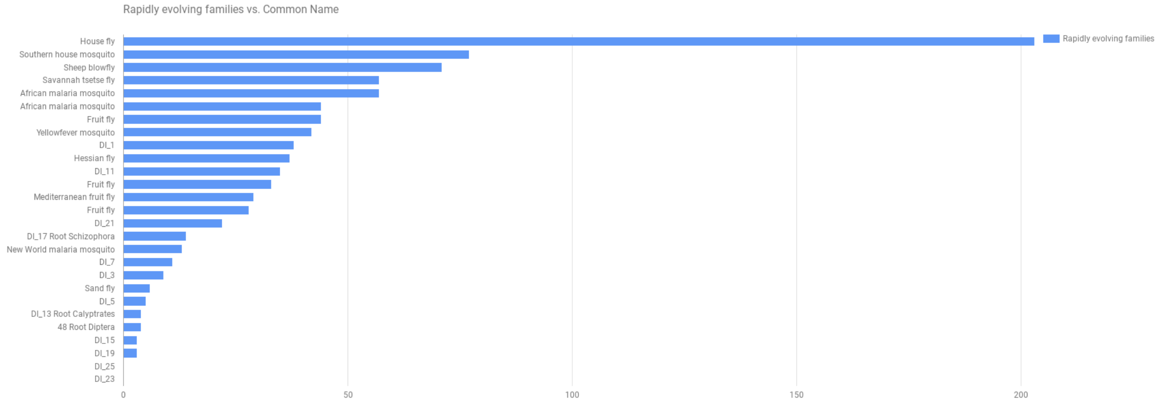

**Figure SG2_31: Comparison of branch lengths from 150 backbone or 1,000's of genes.**

Comparisons of branch lengths estimated by maximum likelihood on concatenated alignments across the Arthropod phylogeny using only the 150 single-copy backbone genes or hundreds to thousands of single-copy genes among taxonomic groups ($r^2$= 0.86).

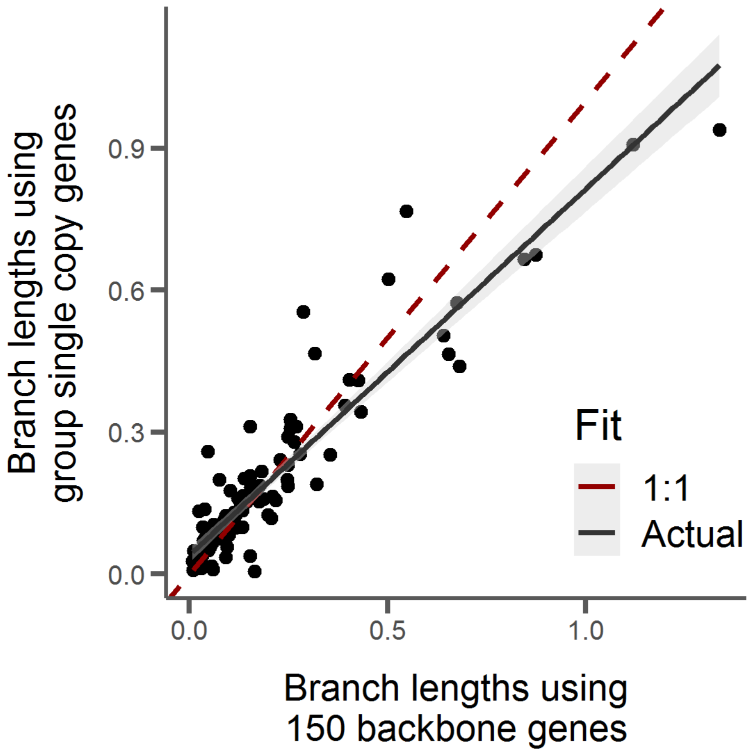

**Figure SG1_32: Jackknife test of gene gains/losses after randomly removing species.**

Distributions of gene gains and losses for every lineage after randomly removing 5 species and repeating for 100 replicates. Red notches indicate original counts with all species. Blue notches indicate counts after removal of three low quality species: LRECL, LHESP, and LLUNA. Note that in most cases red and blue notches overlap and are indiscernible.

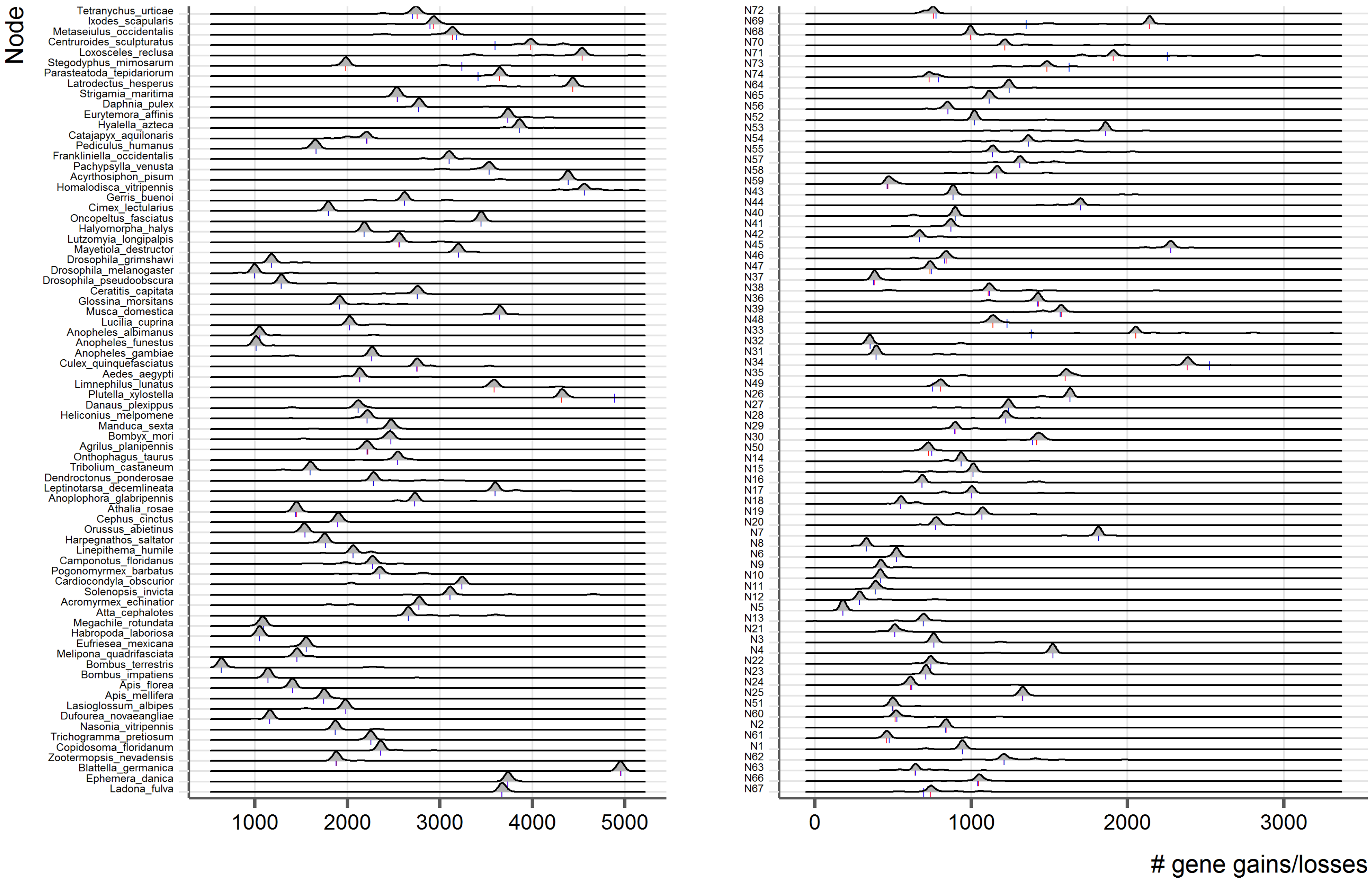

**Figure SG3_33: Jackknife resampling test of phylogeny branch divergence times.**

Distributions of divergence time estimations for every node in the Arthropod phylogeny generated by randomly removing 5 of 22 fossil calibration points 100 times. Red notches indicate estimates when all fossil calibrations used.

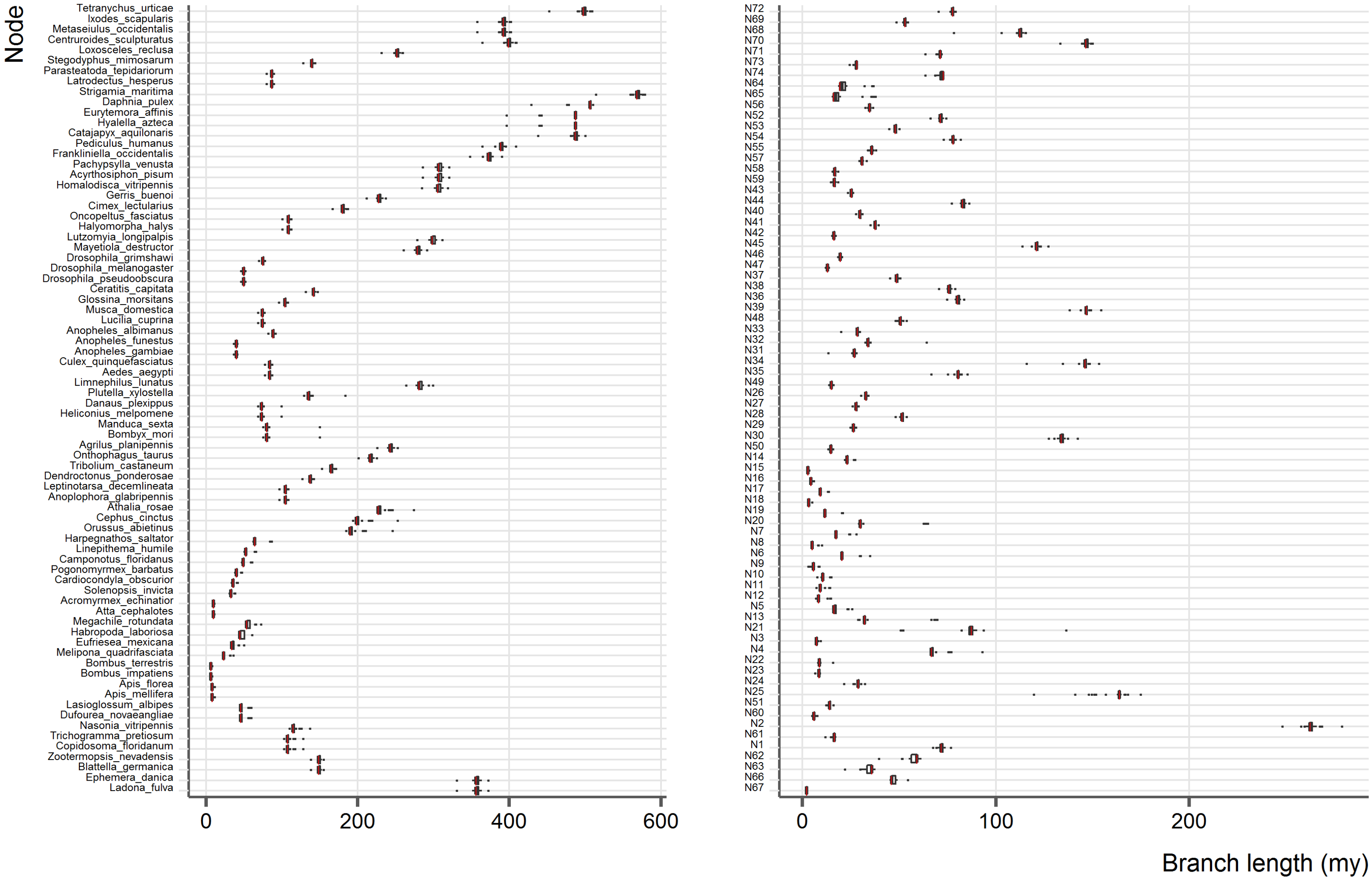

**Figure SE1_34: Jackknife resampling test of total protein domain change events**

Proportion of total protein domain change events (%) by event type for 100 replicates of a jackknife test which removes randomly 5 species from the original phylogeny. Shown are box plots with overlapping swarm plots indicating data point distribution. Red horizontal line represents the value of the original phylogeny, yellow line the value for removal of the three low quality species (LRECL, LHESP, LLUNA).

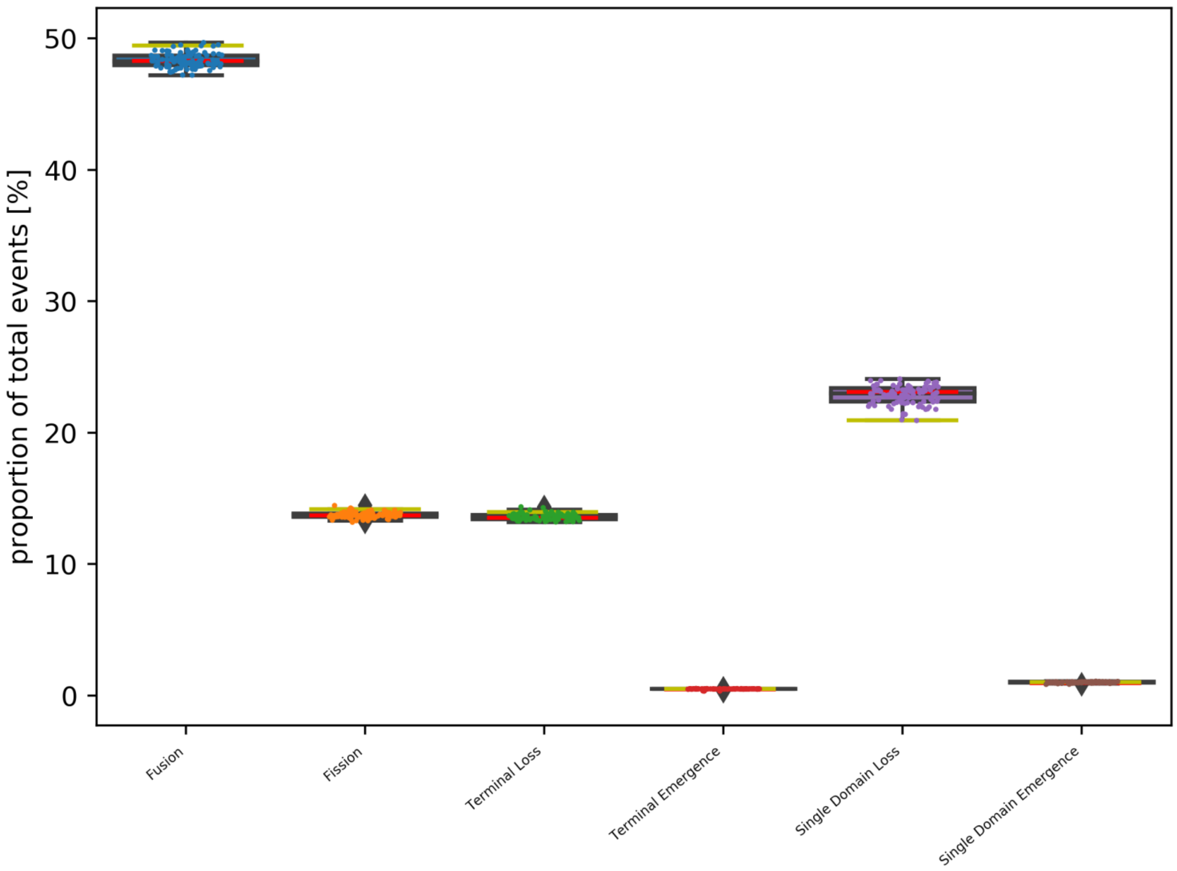
